# Probing the dynamic structure-function and structure-free energy relationships of the corona virus main protease with Biodynamics theory

**DOI:** 10.1101/2020.07.19.211185

**Authors:** Hongbin Wan, Vibhas Aravamuthan, Robert A. Pearlstein

## Abstract

The SARS-CoV-2 Main protease (M^pro^) is of major interest as an anti-viral drug target. Structure-based virtual screening efforts, fueled by a growing list of apo and inhibitor-bound SARS-CoV/CoV-2 M^pro^ crystal structures, are underway in many labs. However, little is known about the dynamic enzyme mechanism, which is needed to inform both structure-based design and assay development. Here, we apply Biodynamics theory to characterize the structural dynamics of substrate-induced M^pro^ activation, and explore the implications thereof for efficacious inhibition under non-equilibrium conditions. The catalytic cycle (including tetrahedral intermediate formation and hydrolysis) is governed by <u>concerted</u> dynamic structural rearrangements of domain 3 and the m-shaped loop (residues 132-147) on which Cys145 (comprising the thiolate nucleophile and one-half of the oxyanion hole) and Gly143 reside (comprising the other half of the oxyanion hole). In particular:

1. Domain 3 undergoes dynamic rigid-body rotations about the domain 2-3 linker, alternately visiting two conformational states (denoted as 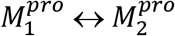).
2. The Gly143-containing crest of the m-shaped loop (denoted as crest B) undergoes up and down translations in concert with the domain 3 rotations (denoted as 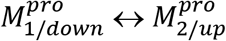, whereas the Cys145-containing crest (denoted as crest A) remains statically in the up position. The crest B translations are driven by conformational transitions within the rising leg of the loop (Lys137-Asn142).

We propose that substrates associate to the 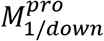 state, which promotes the 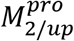 state, dimerization (denoted as 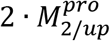-substrate), and catalysis. The structure resets to the dynamic monomeric form upon dissociation of the N-terminal product. We describe the energetics of the aforementioned state transitions, and address the implications of our proposed mechanism for efficacious M^pro^ inhibition under native-like conditions.

## Introduction

M^pro^ is of current interest as an anti-viral drug target, and experimental and *in silico* efforts toward the discovery of potent, efficacious inhibitors are currently underway in many labs across the globe. However, drug discovery is typically a trial-and-error/hit-and-miss undertaking due in no small measure to key deficiencies in the fundamental understanding of molecular and cellular structure-free energy relationships, as well as reliance on <u>equilibrium</u> potency metrics (e.g. IC_50_, K_d_) that are of limited applicability to <u>non-equilibrium</u> conditions *in vivo* [1,2].

In this work, we break from traditional screening and structure-based drug design approaches, and examine M^pro^ inhibition from a <u>theoretical</u>, *in vivo*-relevant perspective, based on multi-scale Biodynamics principles described in our previous work [1,2]. Our theory addresses the fundamental nature of dynamic structure-free energy relationships under aqueous cellular conditions (powered principally by de-solvation and re-solvation costs [2,3]), and the general means by which cellular function is derived from interacting molecular species undergoing time-dependent cycles of exponential buildup and decay. As such, enzyme structure-function is necessarily considered in the overall context of cellular function and dysfunction (consisting of viral infection in this case), and in particular:

1. Synchrony between substrate k_1_, k_cat_, and k_-1_, in which the bound substrate lifetime (t_1/2_) is comparable to 1/k_cat_ (a general kinetic paradigm that was first described by van Slyke and Cullen [4,5]), and product inhibition is circumvented via fast leaving group dissociation.
2. Synchrony between the rates of enzyme and substrate buildup and product formation.

We assume that infection proceeds in the following general phases [6,7]:

1. Virion capture:

a. Receptor binding and internalization.
b. RNA unpacking.
2. Virion “factory” construction:

a. Translation of ORF1a and ORF1ab into polyproteins pp1a containing non-structural proteins (nsp) 1 to 11 and pp1ab containing nsp1 to 16, respectively.
b. Cleavage of the pp1a and pp1ab polyproteins into their constituent nsps:

i. Auto-cleavage of nsp3 (papain-like protease, PPL^pro^) in cis, followed by nsp3-mediated trans cleavage of nsp4.
ii. Auto-cleavage of nsp5 (M^pro^) in cis, followed by nsp5-mediated trans cleavage of nsp6 through nsp11/16. As such, M^pro^ and its substrates build together, which has important implications for therapeutic inhibition.
c. Buildup of the replication-transcription complex (RTC) within cytoplasmic endosome-derived double membrane vesicles [8–11].
3. Virion production:

a. RNA replication.
b. Structural protein translation/processing.
c. Virion assembly/export [12].

Therapeutic intervention is optimally targeted at druggable proteins, including M^pro^, that participate in the earliest steps of infection prior to, or during, the factory construction phase. Clinical anti-viral success depends on reducing the active M^pro^ population below that required for RTC buildup and virion production at a threshold fractional inhibition of the protein population over time, which may be relatively high given that many substrate copies can be cleaved by each free enzyme copy (equating to “leakage” from the inhibited system). As we showed previously, efficacious dynamic occupancy <u>under non-equilibrium conditions</u> is achieved <u>at the lowest possible exposure</u> when the rates of drug association and dissociation are tuned to the rates of target or binding site buildup and decay [1]. In the case of enzymes, including M^pro^, fractional inhibitor occupancy depends on the inhibitor on-rate relative to that of the substrate (denoted as k_1_×[substrate](t) · [M^pro^](t) and k_on_ · [inhibitor](t) · [M^pro^](t), respectively), where [M^pro^](t) is denoted henceforth as k_i_. We assume that M^pro^ builds approximately in tandem with expression of the full length polyprotein via auto-cleavage in cis (noting that trans cleavage of M^pro^ is unlikely, as explained in a later section), and that a relatively constant 1:1 ratio of substrate to post-cleaved enzyme is maintained throughout the factory construction stage due to the simultaneous expression of M^pro^ and its downstream substrates. Two general scenarios for non-covalent competitive inhibition are possible under these conditions:

1. The worst case scenario, in which post-cleaved M^pro^ and its substrates build at approximately the same rate as polyprotein expression (consistent with tuned kinetics). Efficacy depends on high inhibitor occupancy under this scenario (Figure 1A), which in turn, depends on parity between k_on_ and k_i_.
2. The best case scenario (which seems less likely), in which the cleaved products lag behind the polyprotein (consistent with mistuned kinetics) (Figure 1B). Ideal fractional inhibition of the M^pro^ population depends on k_on_ ≈ k_i_, whereas minimal efficacious inhibition depends on k_on_ > k_1_ (where both rate constants are < k_i_).

**Figure 1.**
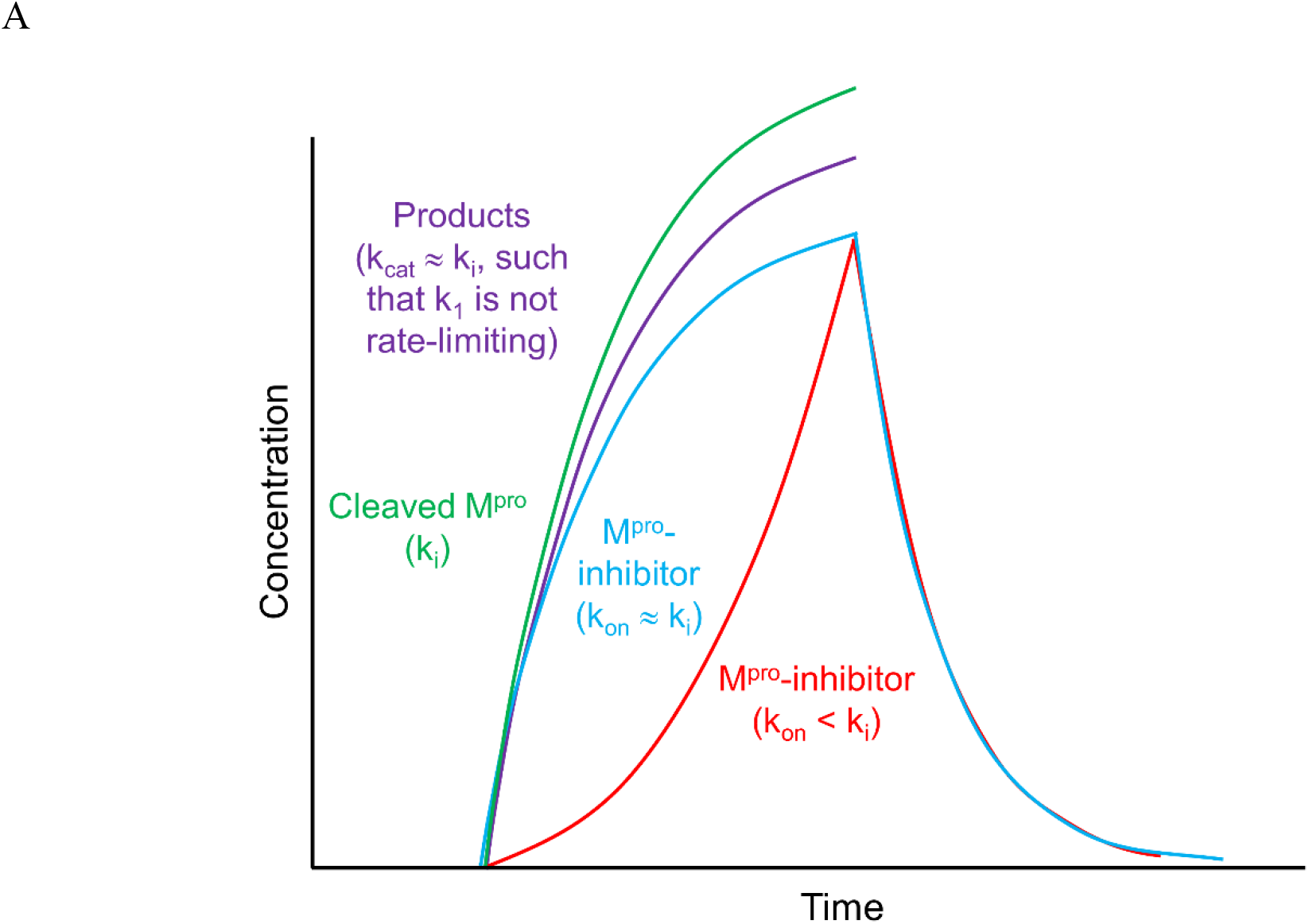
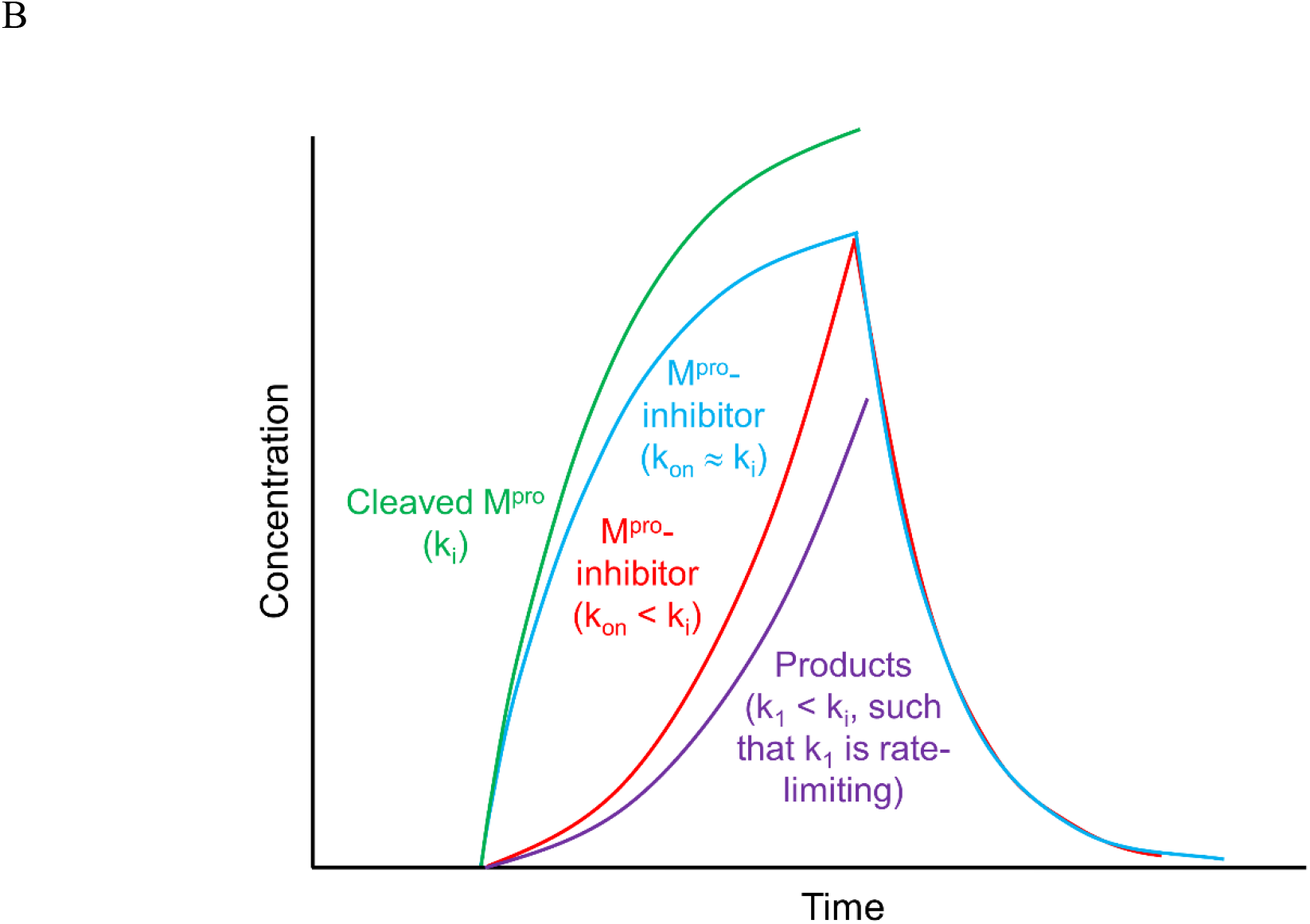
<u>Hypothetical</u> examples of post-cleaved M^pro^ buildup and decay and downstream cleavage products (reflecting substrate association, dissociation, and turnover in aggregate) under the two general scenarios described in the text (the mathematical basis of these plots is explained in [1]). (A) The worst case scenario, in which the rate of product buildup is comparable to k_i_. The plot includes the following quantities: auto-cleaved M^pro^ buildup (green tracing) (noting that M^pro^ decay depends on the existence of a degradation pathway), collective product buildup (purple tracing), and buildup and decay of inhibitor-bound M^pro^ under conditions in which the inhibitor k_on_ ≈ k_i_ (blue tracing), and inhibitor k_on_ < k_i_ (red tracing). (B) Same as A, except for the best case scenario, in which the rate of product buildup < k_i_.

The kinetic tuning requirement may be partially relaxed in the case of covalent inhibition, in which the inhibited enzyme fraction <u>accumulates</u> over time. Covalent inhibition has been used successfully with other anti-viral targets, including hepatitis C NS3 protease [13,14]. However, accumulation rates << k_i_ can likewise result in “leakage” of uninhibited M^pro^ and its downstream products. Understanding the mechanism and dynamics of M^pro^ cleavage and its subsequent activation is essential for differentiating among these scenarios, and informing *in vivo*-relevant inhibitor design.

We collected, classified, and overlaid representative dimeric and monomeric ligand-bound and apo SARS-CoV and CoV-2 M^pro^ crystal structures (see Materials and methods). We then explored and compared the structures using an integrated approach, consisting of 3D visualization and molecular dynamics (MD)-based solvation analysis (WATMD [2,15]) to <u>qualitatively</u> assess the free energy <u>barriers</u> governing the intra-molecular states of M^pro^ (denoted as monomeric 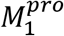 and 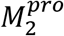, and dimeric 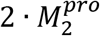, as well as the association and dissociation barriers governing substrate and inhibitor occupancy. We then investigated the structure and function of M^pro^, focusing on the interrelationship between the catalytic and substrate/inhibitor-binding mechanisms. Questions of interest include:

1. The basis of substrate- and dimer-induced activation and specificity of the catalytic site.
2. The interplay between covalent/mechanism-based inhibitor binding kinetics and the structural dynamics of the protein.
3. The interplay between catalytic turnover and viral dynamics governing the buildup of post-cleaved M^pro^ and its substrates.

### Overview of M^pro^ structure and catalytic function

Monomeric M^pro^ consists of three domains, including domain 3 and domains 1 and 2 that together subserve the protease function (Figure 2A) [16]. Domains 1, 2, and 3 respectively comprise the “ceiling”, “floor”, and “basement” of the active site (AS), and are organized in a loosely packed arrangement that promotes high sensitivity of protein structure-function to the monomeric versus dimeric forms, the pre-versus post-cleaved species, and the unbound versus substrate-bound states. Domains 1 and 2 versus domains 2 and 3 are connected by short and long flexible linkers, respectively, resulting in a tighter 1-2 inter-domain relationship compared with the looser, more dynamic relationship among domains 2 and 3. We denote this hierarchical inter-domain relationship as {1-2}-3, consistent with the crystallographically observed conformations in the monomeric 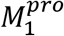 and 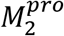 states (PDB code = 2QCY) and the dimeric 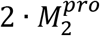 form (PDB code = 6M03, 2Q6G, and many others) (noting that <u>monomeric</u> 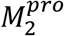 is unobserved experimentally). The backbone NH groups comprising the oxyanion hole (contributed by Cys145 and Gly 143) reside on a three-dimensional double-crested m-shaped loop, hereinafter denoted as the “m-shaped loop” (Figure 2B). The N-terminal leader sequence (denoted as NTL) and C-terminal tail sequence (denoted as CTT), the latter of which includes a small C-terminal helix, play key roles in organizing the AS, m-shaped loop, and dimer interface. We assume in this work that the substrate binding and catalytic <u>machineries</u> are well-conserved in CoV and CoV-2, which differ by 12 residues, half of which are located in domain 1 (including one located at the upper boundary of the AS), two in domain 2, and four in domain 3 [17]). As such, CoV and CoV-2 structures were used interchangeably in our analysis.

**Figure 2.**
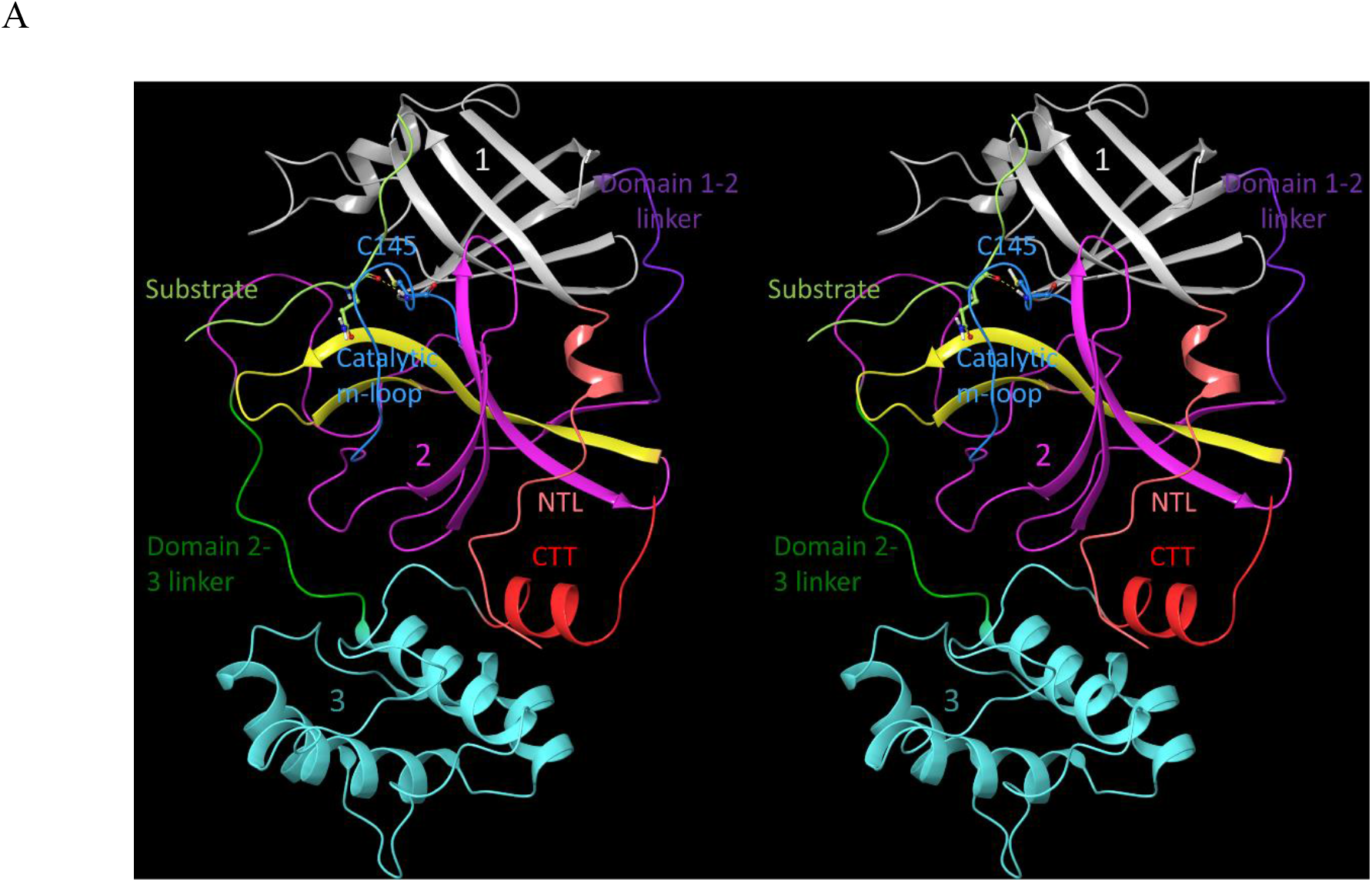
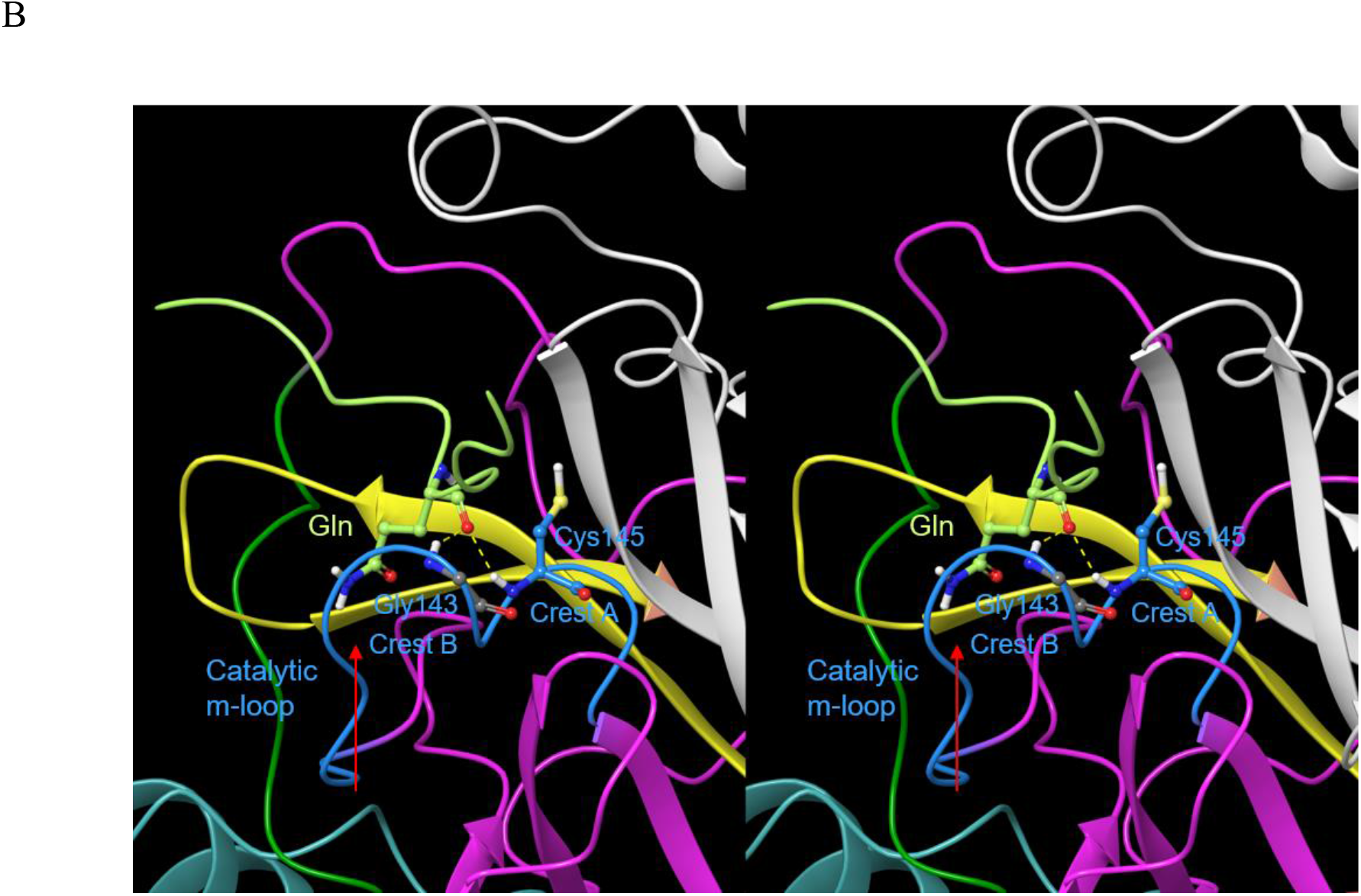
Stereo views of key M^pro^ structural features. (A) Domains 1 (white), 2 (magenta), and 3 (cyan) exemplified by PDB 2Q6G (chain B), illustrating the canonical S-shaped topological inter-domain architecture and key structural features of M^pro^. The three domains are interconnected by flexible linkers (domain 2-3 and 1-2 linkers shown in dark green and purple, respectively). The substrate peptide (light green) binds to the upper strand of a β-hairpin loop (yellow) via the backbone NH and C=O groups of Glu166. The catalytic m-shaped loop (blue) contains Cys145 and the backbone NH groups of Cys145 and Gly 143 comprising the oxyanion hole (one residue per crest). The NTL (coral), denoted by others as the “finger peptide” [18], projects into the dimer interface, together with the CTT (red). (B) Close-up view of the AS and oxyanion hole, showing the positioning of the substrate P1 Gln side chain (light green) in the monomeric S1 pocket, together with the backbone NH-substrate H-bonds. Cys145 and Gly143 reside respectively on crests A and B of the m-shaped loop. The N-to C-terminal directionality of the rising leg of the m-shaped loop is denoted by the red arrow.

Serine proteases function via a common catalytic mechanism conveyed by an Asp-His-Ser triad. However, a His-Cys dyad appears sufficient for proton abstraction from the more acidic Cys (relative to Ser) of cysteine proteases, leading to an activated thiolate-His ion pair [19–21]. The M^pro^ catalytic mechanism may be summarized as follows (the residue numbering throughout this work refers to the SARS-CoV-2 variant):

1. Abstraction of the Cys145 proton by His41, resulting in a nucleophilic thiolate moiety (stage 1 proton transfer).
2. Substrate binding, followed by nucleophilic attack on the scissile bond, resulting in a transient tetrahedral intermediate (TI) which is stabilized by the backbone NH dyad comprised of Gly143 and Cys145 (referred to as the “oxyanion hole”). This step is claimed to be extremely fast in other cysteine proteases (requiring stopped flow measurement) [20].
3. Spontaneous TI decay to the N-terminal leaving group (product 1) and thioester adduct.
4. Hydrolysis of the thioester adduct (stage 2 proton transfer), resulting in the C-terminal leaving group (product 2). This step is claimed to be rate-determining in other cysteine proteases [22].

However, alternate catalytic <u>triad</u>-based mechanisms have been proposed M^pro^, including:

1. Substitution of the canonical Asp of the catalytic triad by a high occupancy water molecule observed near His41 in many M^pro^ crystal structures and our WATMD calculations (possibly a weaker surrogate for Asp) [21,23], noting the absence of this water in subunit B of 2Q6G due to repositioning of Asp187 (which if catalytically essential, would result in enzyme inactivation).
2. Rearrangement of Asp187 from its observed pairing with Arg40 to His41 [13]. Given the strategic location of the Arg40-Asp187 ion pair opposite to the domain 1-2 linker in all of the structures that we examined (exhibiting a latch-like appearance), stabilization of the domain 1-2 interface by this shielded ion pair is the more likely scenario (Figure 3).

**Figure 3.**
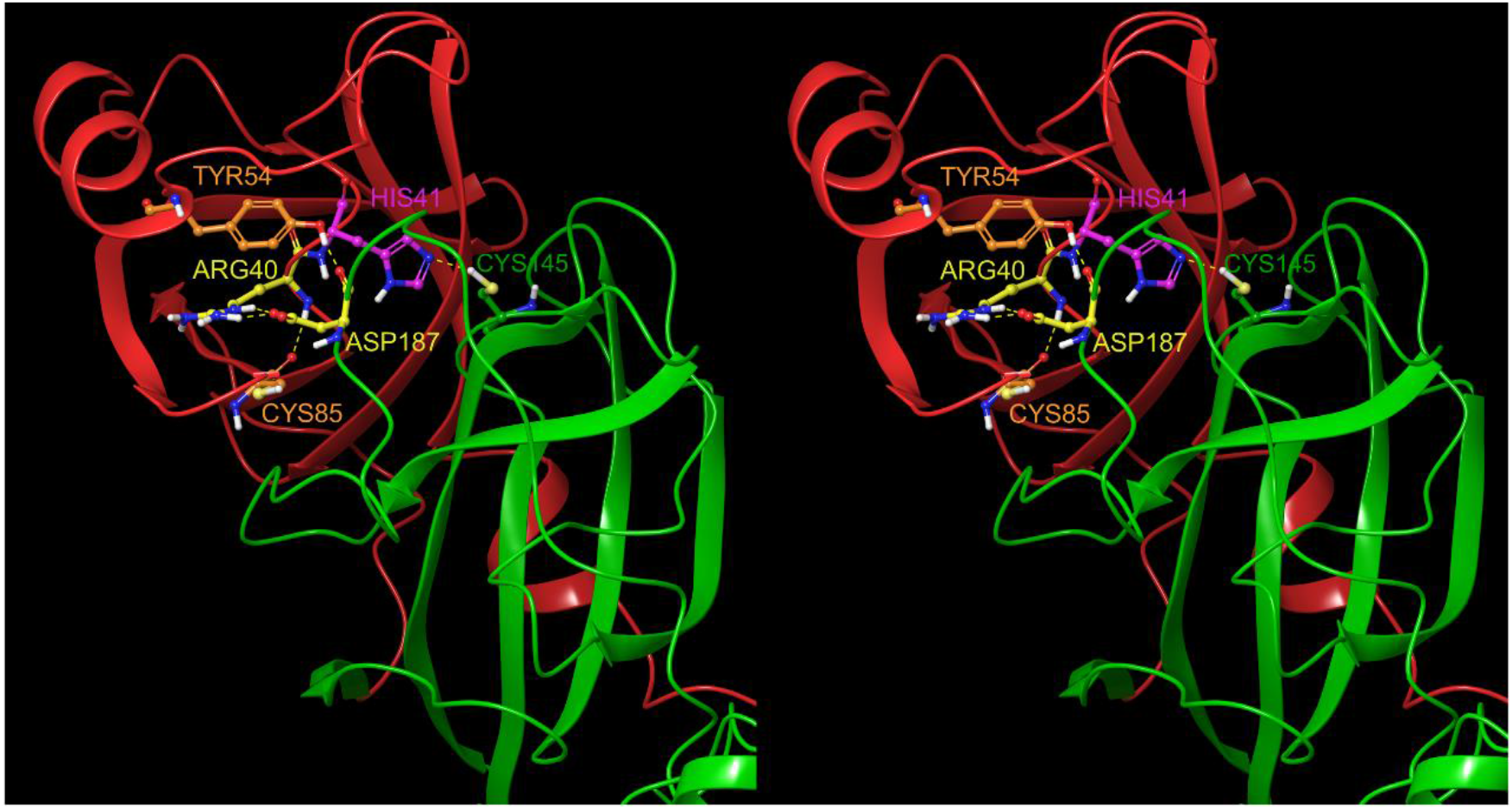
The Arg40-Asp187 ion pair (yellow side chains), which is shielded between Tyr54 and Cys85, likely stabilizes the domain 1-2 interface and upper region of the domain 2-3 linker.

## Materials and methods

### Structural data and visualization

All M^pro^ structures used in our study were obtained from the RCSB databank [24], and grouped according to species, site-directed mutants, ligand/substrate-bound forms, and dimeric and mutant monomeric forms (Table 1).

**Table 1.** Structures used in our analysis [23,25–28] (those on which we performed WATMD calculations are denoted by a ‡).

| PDB code | Species | Form | Mutation(s) | Ligand/substrate-bound | Crystallization pH |
| --- | --- | --- | --- | --- | --- |
| 2QCY‡ | SARS-CoV | Monomer | R298A | - | 6 |
| 2Q6G‡ | SARS-CoV | Dimer | H41A | Substrate | 6 |
| 6M03‡ | SARS-CoV2 | Dimer | - | - | 8.1 |
| 6WNP‡ | SARS-CoV2 | Dimer | - | Boceprevir | 7.5 |
| 2BX3 | SARS-CoV | Dimer | - | - | 5.9 |
| 6LU7 | SARS-CoV2 | Dimer | - | N3 | 6 |
| 4KTC | Hepatitis C virus NS3 protease | Dimer | A156T | 1X3 | 6.2 |
| 4CHA | α-chymotrypsin | Dimer | - | Substrate | NA |

All calculations and structural visualizations were performed using WATMD [2,15,29], Maestro version 2020-1 (Schrodinger, LLC), and PyMol version 2.0 (Schrodinger, LLC). 2QCY, 2Q6G, 6M03, and 6WNP were prepared for WATMD calculations using the pprep tool in Maestro, and the resulting structures were aligned using the structalign tool in Maestro. The aligned dimeric structures and their disassembled A and B chains were compared visually using Maestro. We emphasize that this is a first principles theoretical study which involves limited use of conventional molecular modeling techniques.

### General non-equilibrium structure-free energy relationships used in this work

Whereas drug-target binding and efficacy are considered in terms of equilibrium free energy models throughout mainstream cell biology and pharmacology, it is apparent that living systems (including virally infected cells) depend to a very large degree on non-equilibrium operation. We previously reported a first principles theoretical treatment of non-equilibrium structure-function-free energy relationships referred to as Biodynamics [1,2]. Spontaneous non-covalent intra- and inter-molecular interactions, by definition, lower the total system free energy (i.e. ΔG = G_non-interacting_ – G_interacting_ < 0). However, ΔG is a non-sequitur under conditions in which the concentrations of the participating species are changing over time (non-equilibrium conditions, by definition), in which case binding free energy is partitioned into distinct barriers governing the rates of entry and exit to/from each available structural state (denoted as 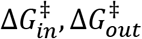. Fractional occupancy of a given state is proportional to the relative rates of entry and exit to/from that state (versus ΔG = −RT · ln(K_d_)), wherein the entry-exit process can cycle throughout the lifetimes of the participating species (see below). We proposed previously that, under aqueous conditions, 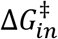 and 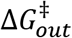 respectively equate to the de-solvation and re-solvation costs of entering and exiting non-covalent states [2]. The de-solvation cost equates to the total loss of hydrogen bond (H-bond) free energy from the expulsion of H-bond enriched solvation from the intra- or inter-molecular interface to bulk solvent, whereas the re-solvation cost equates to the total loss of H-bond free energy of water returning from bulk solvent to H-bond depleted positions of the interface. Polar/charged solute surfaces promote H-bond enriched solvation relative to bulk solvent, resulting in free energy gains. The de-solvation cost of such water during intra- or inter-molecular rearrangements depends on the extent to which the disrupted H-bonds are replaced by intra- or inter-molecular counterparts (typically, a zero sum game at best). The rate of entry to a given state is proportional to the <u>de-solvation</u> cost of that state, and occupancy increases with increasing rate of entry at a constant rate of exit. The rate of exit from a given state is proportional to the <u>resolvation</u> cost of that state, and occupancy increases with decreasing rate of exit at a constant rate of entry. As such, the occupancy of a given state may accumulate via fast entry, slow exit, or both (to the limit of the total 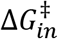 and 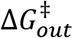. Energetically favorable charged/polar interactions, including charge-charge, charge-neutral, and neutral-neutral, drive protein folding and rearrangements of folded macromolecules under aqueous conditions by:

1. Lowering the total de-solvation cost via optimal mutual water H-bond replacements (thereby speeding the rate of entry).
2. Electrostatic free energy gains in <u>shielded</u> charge-charge scenarios (thereby slowing the rate of exit).

The energetic driving force of all non-covalent rearrangements (including protein folding) consists principally of the expulsion of H-bond depleted solvation. The persistence of a given state depends on the total free energy of the expelled solvation (amounting to the re-solvation cost incurred upon exiting). Persistent states result from complete expulsion of H-bond depleted solvation, whereas cyclic rearrangeability depends on a “whack-a-mole-like” scenario, in which expulsion of H-bond depleted solvation from one region is accompanied by the generation of an equivalent amount at another (resulting in an oscillation between the contributing states—a kind of molecular “motor”).

### WATMD calculations

Solvation calculations were performed using our in-house WATMD software, as described previously [2]. Briefly, the prepared protein structures (corrected for proton states, Asn/Gln and His flips, missing atoms, and net charge) were simulated in AMBER 16 with ff14SB force-field [30] at 300° K without restraints using periodic boundary conditions in a TIP3 water box of 8 Å between any protein atom and the edge of the box. The simulations consisted of a 0.5 ns equilibration phase, followed by a 10 ns production phase sufficient to achieve convergence of the solvating water, resulting in 40,000 frames per trajectory. pH-dependent M^pro^ structure and substrate recognition, and the possibility of pH-driven structure switching has been suggested by other workers on the basis of the observed pH dependence of M^pro^ structure [16,31]. However, M^pro^ appears to operate exclusively within the cytoplasmic double membrane vesicle environment, the pH of which is likely ~7.0-7.4 in the absence of proton pumps. As such, M^pro^ simulations at pH 7.0 seem justified.

The 40,000 frame AMBER trajectories were first aligned to a common reference frame about a specified set of template residues, and analyzed in terms of time-dependent water occupancy (i.e. the frequency of water visits) within each 1 Å voxel of a three-dimensional lattice containing the time-averaged protein structure. H-bond enriched and depleted solvation regions were assigned on the basis of high (un-trapped or trapped) and low occupancy voxels relative to bulk solvent, respectively. The resulting water occupancies >> bulk and << bulk (denoted as hotspots) are represented by color-coded spheres, the radii of which are proportional to the occupancy level. The color-coding is based on a gradient representing the strength and type of the protein H-bond partner(s): strong donor (bright red) ⟶ weak/deficient partner (light pink/light blue/white) ⟶ strong acceptor (bright blue). Solvating water was assumed to be stationary relative to flexible protein substructures (analogous to a boundary layer effect), necessitating separate substructure-specific water occupancy analyses (specified by substructure-specific alignment templates). For example, the Met164-Glu166 β-strand in domain 2, a key AS element that undergoes significant motion during the simulations (shown in yellow in Figure 2), was used as the template for generating the solvation structure within and around the binding pocket (noting however that bound ligands in all crystal structures are agnostic to this motion). We calculated the solvation structures of the following M^pro^ states, from which we inferred the solvation free energy barriers governing the 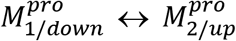 and 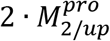 state transitions, together with substrate and inhibitor association and dissociation barriers:

1. The <u>apo</u> form of <u>monomeric</u> 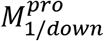 (2QCY) and putative <u>substrate-bound</u> form of <u>monomeric</u> 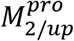 (PDB code = 2Q6G <u>with one chain removed</u>), focusing on:

a. The <u>AS</u> solvation structure in 2QCY, which informs about substrate k_1_, k_-1_ and inhibitor k_on_, k_off_. We examined the correspondences between the solvation hotspots and a set of representative inhibitors (Table 2) extracted from additional CoV and CoV-2 M^pro^ structures, which we overlaid on the 2QCY time-averaged protein structure.
b. The <u>domain 2-3 interface</u> in 2QCY and 2Q6G, which informs about the 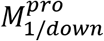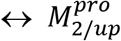 transition barrier.
c. The <u>pre-dimer interface</u> in a single subunit of 2Q6G, which informs about the monomer ⟷ dimer transition barrier.
2. The <u>apo</u> form of <u>dimeric</u> 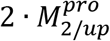 in 6M03, focusing on the <u>AS</u> solvation structure (hypothetical).
3. The <u>boceprevir-bound</u> form of <u>dimeric</u> 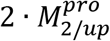 (PDB code = 6WNP), focusing on the <u>occupied AS</u> solvation structure, which informs about the residual solvation of the bound state (H-bond enriched slows substrate k_-1_; inhibitor k_off_).

**Table 2.** Crystallized inhibitor structures used in our study.

| PDB code | Inhibitor |
| --- | --- |
| 1WOF | I12 |
| 2HOB | PRD 002214 |
| 3TNT | G85 |
| 4MDS | 23H |
| 6W63 | X77 |
| 7BRR | K36 |

## Results

### Overview of M^pro^ structural dynamics

Analysis of the M^pro^ crystal structures in our study suggests the existence of a complex substrate-binding mechanism in both CoV and CoV-2 variants. This mechanism can be dissected into four interdependent switchable dynamic contributions (Figure 4) consisting of:

1. Rigid-body rotation of domain 3 relative to domains {1-2}, in which domain 3 oscillates between the 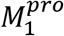 and 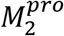 states (noting that dimerization occurs <u>fastest</u> in the substrate-bound 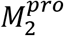 state). The trajectory is guided by transient rearrangements within a large H-bond network spanning within and between the dimeric subunits.
2. Cooperative state transitions between domain 3 and the rising leg of the m-shaped loop, in which the lower energy 3_10_ helix melts into the higher energy extended chain (denoted together as 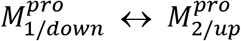. The free energy difference between these states is attributable to solvation-mediated rearrangements (see below). Monomeric 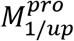 is ruled out by our mechanism, and monomeric 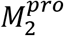 is highly transient (noting that neither of these states is observed experimentally).
3. Cognate substrate and inhibitor binding to 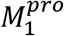, which transiently stabilizes both the dimerization-competent monomeric 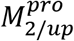 state and the dimeric 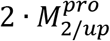 state.
4. Dimerization (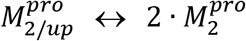 and 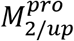-substrate 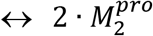-substrate). We postulate that dimerization occurs more slowly in the unbound 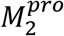 state, which is consistent with the higher observed substrate-independent K_d_ [32] (see below).
5. Catalytic turnover from the substrate-bound 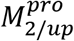 state:

a. Thioester adduct formation.
b. Amide bond cleavage.
c. Dissociation of the C-terminal product.
d. Hydrolysis of the adduct.
e. Dissociation of the N-terminal product.
6. Dimer dissociation, and return to step 1.

**Figure 4.**
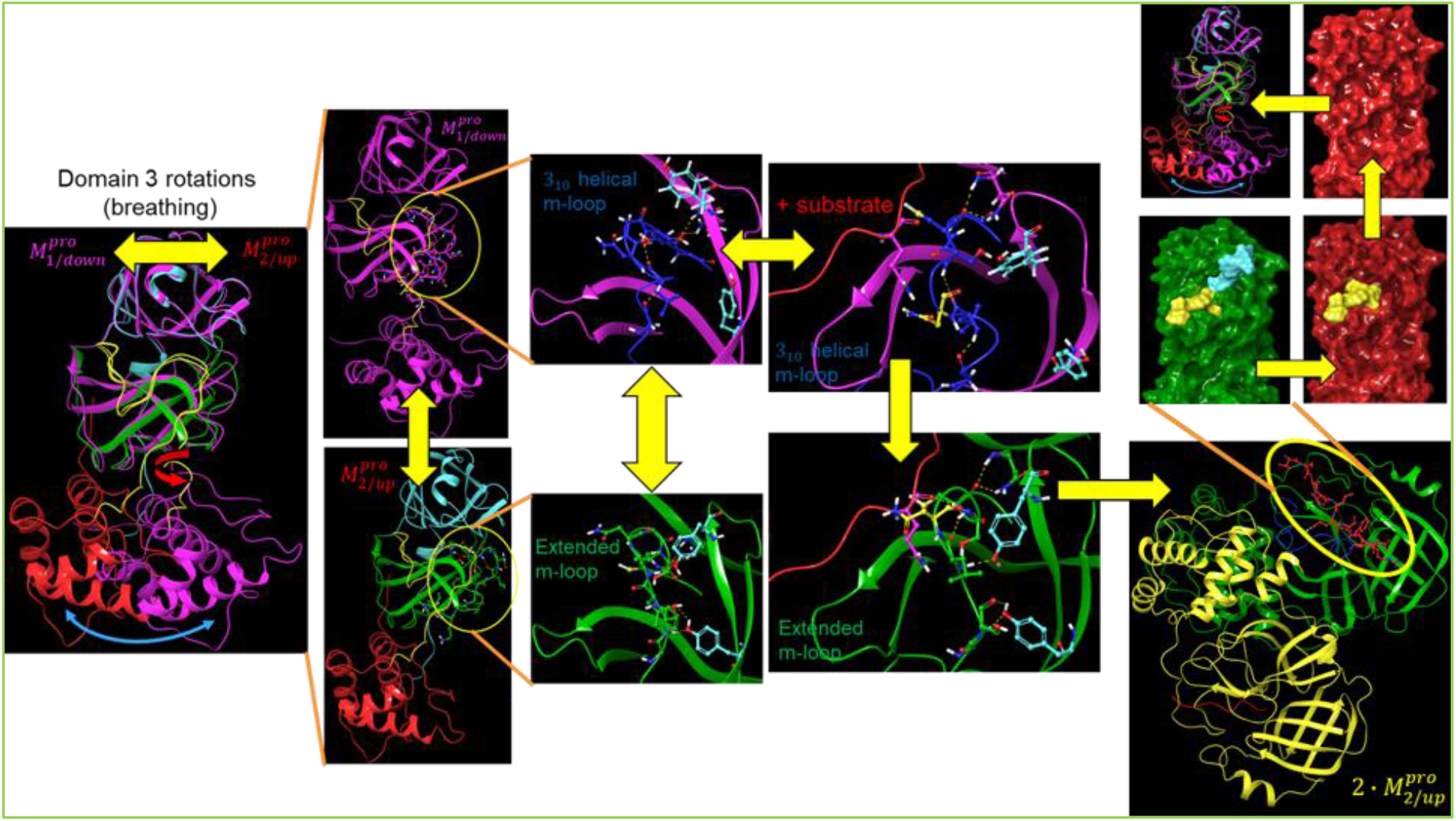
Overview of our proposed dynamic M^pro^ mechanism. Substrate association occurs primarily in the 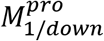 state, in which the S1 pocket is accessible. The substrate-bound 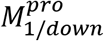 state transitions to the 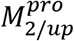 state, which in turn, transitions to the dimeric 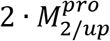-substrate complex. 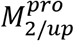-substrate and 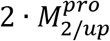-substrate are more stable than the unbound forms of the protein, the t_1/2_ of which is likely on the order of the substrate turnover timescale.

The 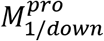 and 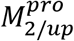 state transitions are guided by specific rearrangements within a large H-bond network spanning across the domain {1-2}-3 interface of the monomeric form, and additionally across the dimer interface of 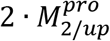. Here, we focus on the configurational rearrangements within this network that switch M^pro^ between the substrate binding, dimerization, and catalytically competent states. The detailed effects of these rearrangements on the domain {1-2}-3 interface, m-shaped loop conformation, and dimer interface are addressed in the following sections. The dilemma for all dynamic intra- and inter-molecular rearrangements relates to the tradeoff between specificity and transience, which according to Biodynamics, is typically achieved via counter-balancing between favorable and unfavorable contributions. The fastest rearrangements prevail, and the balance is tipped transiently toward specific condition-dependent states so as to avoid equilibration. Specificity/recognition is enhanced by high de-solvation costs, which are offset optimally by <u>cognate</u> H-bond partner(s) (noting that electrostatic gains are necessarily balanced against de-solvation costs under unshielded conditions).

### Intra-molecular rearrangements

#### Solvation-powered conformational transitions of domain 3 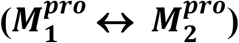

Rigid-body rotations about the domain 2-3 linker chain governing the 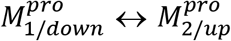 state transition in monomeric CoV M^pro^ can be inferred from the crystal structure of the Arg298Ala mutant (2QCY), together with nearly all of the dimeric apo and ligand-bound CoV and CoV-2 structures (e.g. 6M03) (Figure 5).

**Figure 5.**
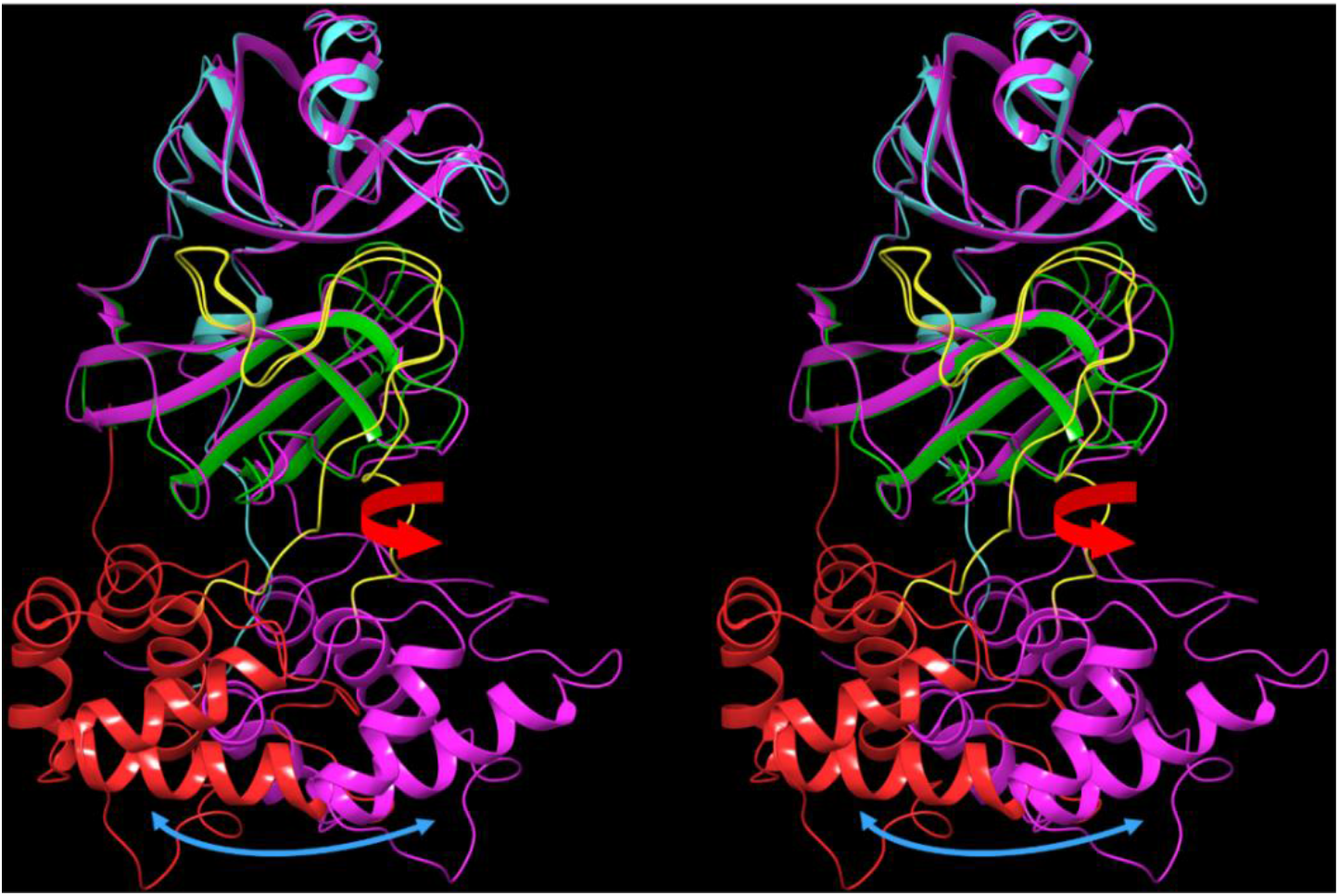
Stereo view of monomeric CoV M^pro^ overlaid on chain A of the apo dimer in 2QCY (domain 3 shown in red) and 6M03 (domain 3 shown in magenta). Spontaneous rigid-body rotation of domain 3 relative to domains {1-2} is denoted by the double-headed blue arrow (noting that the domain 3 structures extracted from the full length protein are approximately superimposable). This rotation is achieved via backbone bond rotations (red arrow) within the domain 2-3 linker region (yellow), which are likely hindered in the substrate-bound state.

We compared the crystal structures of monomeric 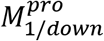 CoV M^pro^ with those of several dimeric 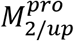 protein structures, focusing on key residues in the H-bond network. The rigid-body domain 3 rotation is guided by transient H-bond switching among these residues, which is best observed by flipping through the Supplementary Information file. The network can be divided into three interacting zones, which undergo concerted signaling into the AS and dimer interface in 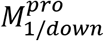 (Figure 6A) and 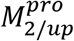 (Figure 6B):

Zone 1: <u>The domain 2-3 linker zone</u>, consisting of an H-bond network centered around Arg131 (Figure 7). This zone is fully disrupted in 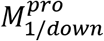.
Zone 2: <u>The m-shaped loop zone</u>, consisting of a ring-like H-bond network comprised of the side chains of Ser139, Glu290, Asp289, and Lys137 (Figure 8). This zone is largely disrupted in 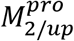.
Zone 3: <u>The CTT/NTL zone</u>, which together with zone 1, governs the rigid-body rotation of domain 3, together with the position of Tyr118, and additionally promotes dimerization (the NTL in particular) (Figure 9).

**Figure 6.**
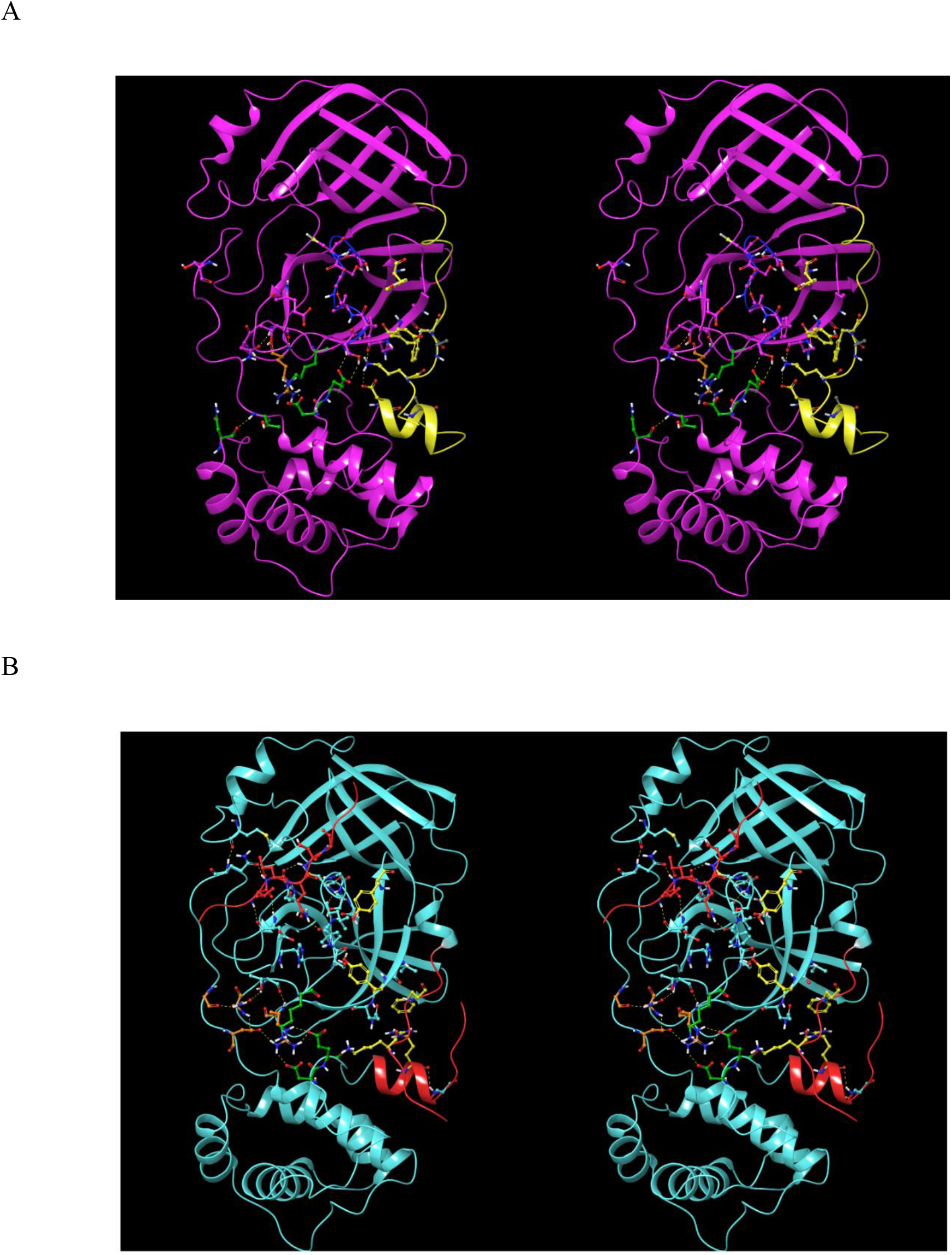
(A) The three zones of the H-bond network in 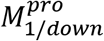 of 2QCY. The network partners switch between this and the 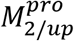 state. Zone 1 (orange side chains), which largely governs the domain 2-3 linker conformation, is disconnected from zone 2 (green side chains) in this state. Zone 2, which bridges between the domain 2-3 linker and the rising stem of the m-shaped loop is well connected in this state (helping to maintain the 3_10_ helical conformation). Zone 3 (yellow side chains), which governs the conformations of Tyr118 and Tyr126, is stabilized by the NTL via Lys5 and the backbone NH of Phe8. (B) Same as A, except for 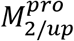 in 2Q6G. Zone 2 merges with zone 1 at the Arg131 nexus in this state, and zone 3 is largely disrupted in this state.

**Figure 7.**
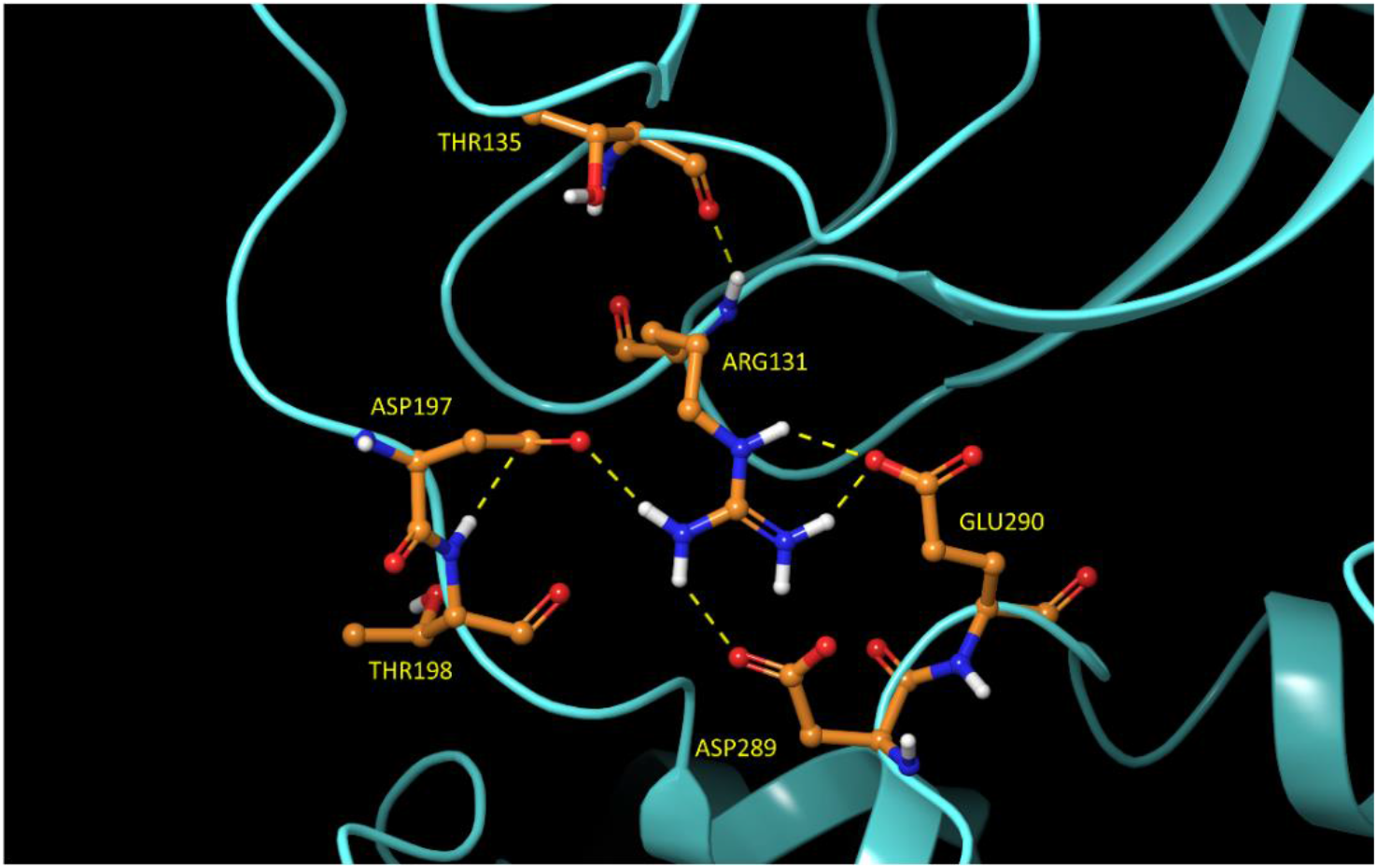
<u>Zone 1</u> of the domain 2-3 H-bond network in 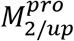 of 2Q6G. The domain 2-3 linker is guided to 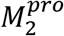 in this network configuration. Glu290 and Asp289 switch to zone 2 in the 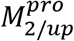 state.

**Figure 8.**
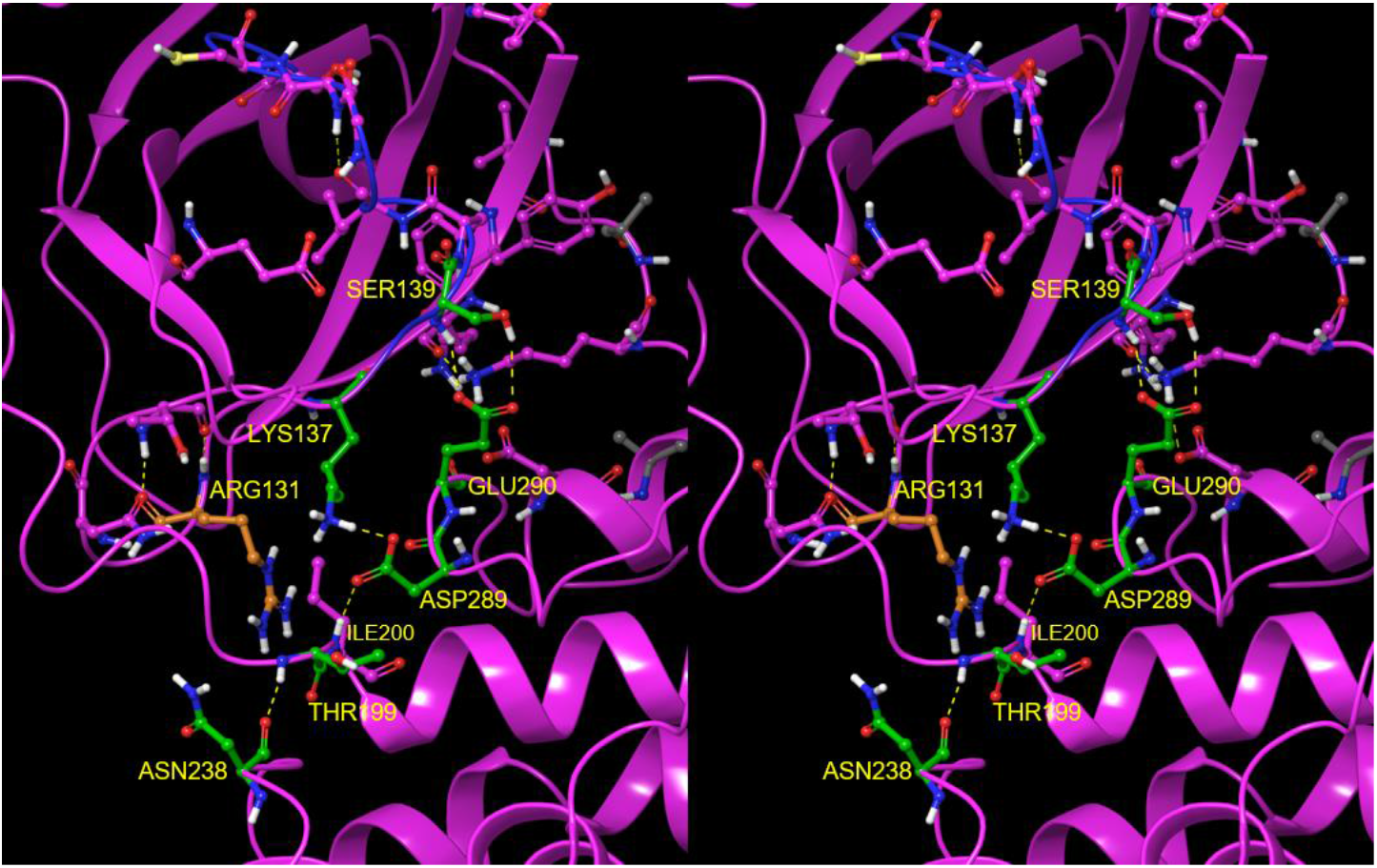
Stereo view of <u>zone 2</u> of the domain 2-3 H-bond network in 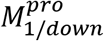 of 2QCY, which forms a circuit (residues highlighted in green) comprised of the side chains of Ser139 (residing just below crest B of the m-shaped loop), Glu290 and Asp289 (both residing on domain 3), and Lys137 (residing at the base of the m-shaped loop). The circuit connects with the backbone NH of Ile200 and backbone oxygen of Asn238 (both of which resided at the base of the domain 2-3 linker). Asp289 and Glu290 switch to zone 1 in the 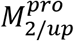 state.

**Figure 9.**
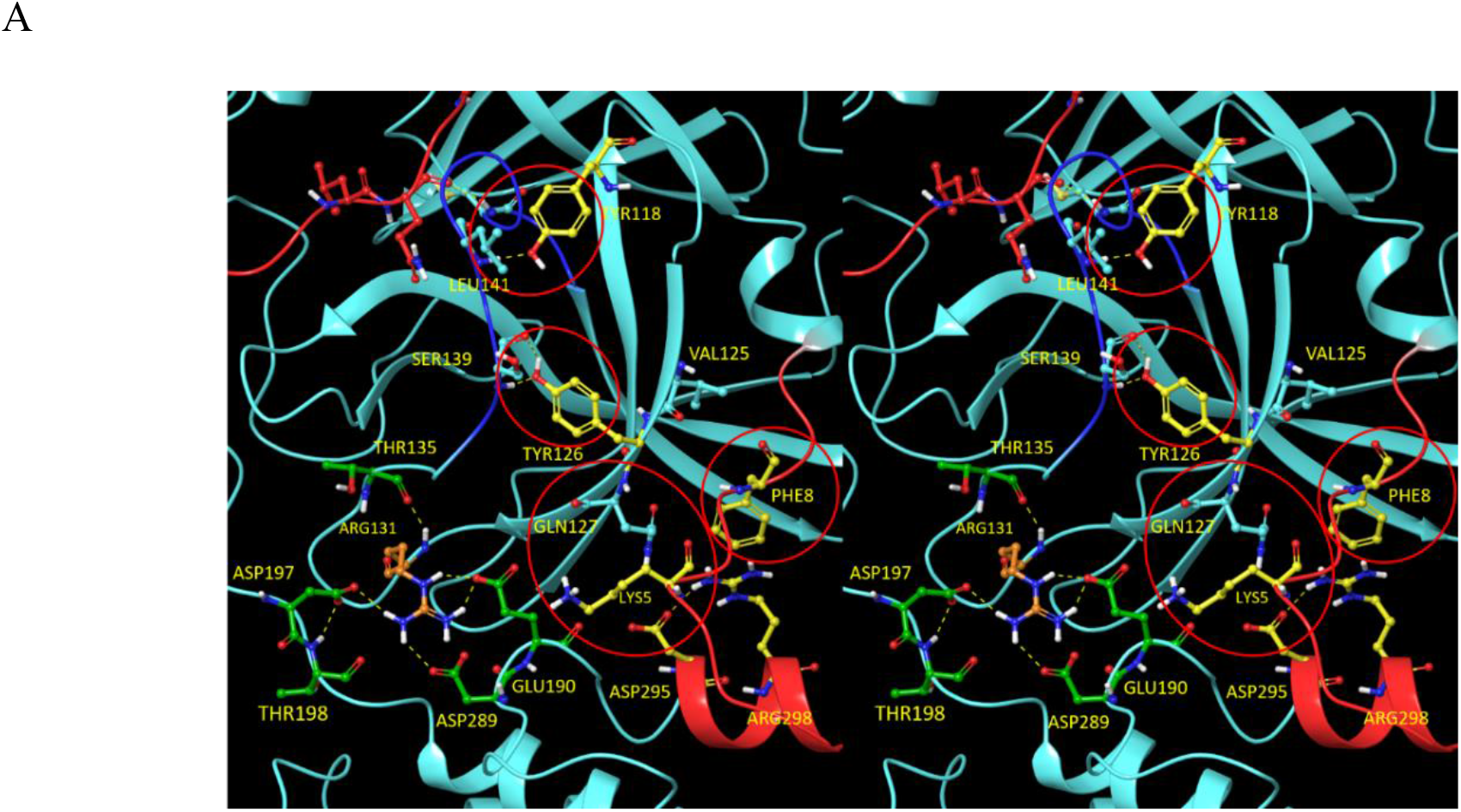
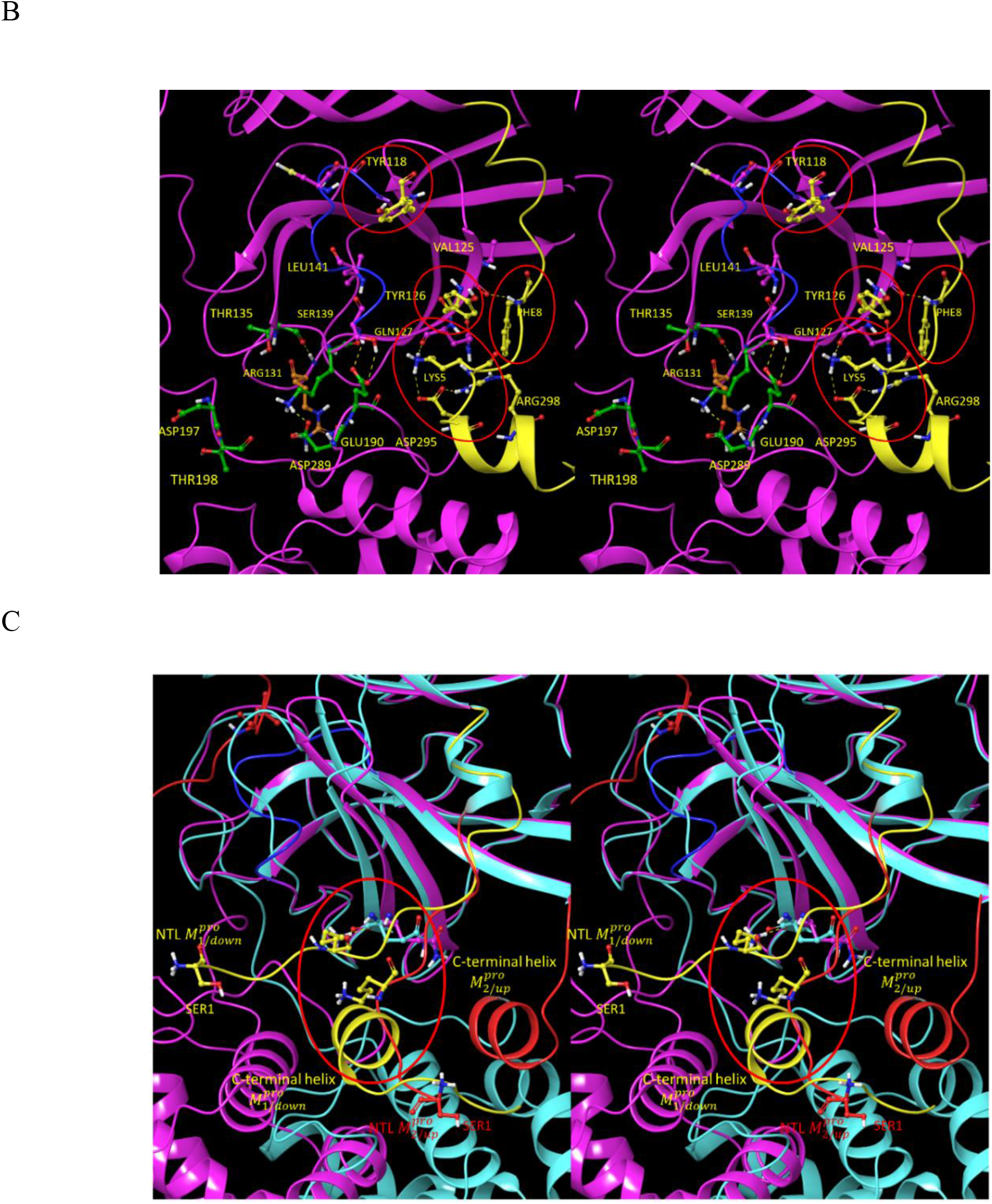
Stereo view of <u>zone 3</u> of the domain 2-3 H-bond network. (A) The 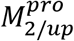 state of 2Q6G. This zone governs the β-hairpin (Gln110-Asn133) conformation on which Tyr118 and Tyr126 reside. The β-hairpin conformation in this state depends on H-bonds between Lys5 of the NTL and backbone oxygen of Gln127 (which is further stabilized by Arg298), together with the backbone NH of Phe8 and backbone oxygen of Val125. H-bonds between Tyr118 and Tyr126 and the backbone NH of Leu141 and backbone oxygen and NH of Ser139, respectively, help promote the <u>extended</u> m-shaped loop conformation in the 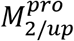 state. The 3_10_ helical conformation in the 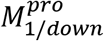 state occurs in the absence of the two Tyr H-bonds, together with additional zone 2 contributions. (B) The 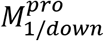 state of 2QCY. The β-hairpin twists in the absence of the Tyr H-bonds in this state, resulting in rotation of Tyr118 and Tyr126 away from the m-shaped loop. (C) The C-terminal helix and NTL in 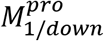 (yellow) and 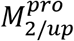 (red). This helix, which is rotated toward the left in 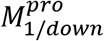 overlaps with the NTL in 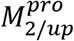 (circled in red), and as such, is pushed away in the 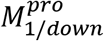 state (blue arrow pointing toward the southwest). The Lys5-Gln127 H-bond is disrupted in this altered NTL trajectory, which signals into Tyr118 and Tyr126 via the β-hairpin. Next, we examined the B-factors in the monomeric 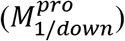 and several dimeric 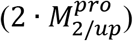 crystal structures as a qualitative metric of the energetic stability of the H-bond network (Figure 10). The data suggests that the H-bond network in the 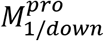 state is stable (B-factors ranging largely between white/light blue/dark blue) (Figure 10A), compared with the significantly less stable network in the dimeric apo 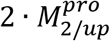 state (B-factors ranging between white/pink/bright red) (Figure 10B). The B-factors of the cognate substrate-bound structure (Figure 10C) are only slightly warmer than those of 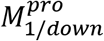, consistent with substrate-mediated stabilization of the 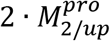 form. The boceprevir-bound 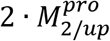 B-factors are comparable to those of the substrate-bound structure (Figure 10D), whereas those of the N3 bound structure are far warmer (nearly comparable to the apo structure) (Figure 10E). The non-native 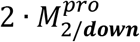 state (PDB code = 2BX3), in which the extended m-shaped loop conformation ordinarily found in this state has been short-circuited, is consistent with the warm B-factors in the rising leg of the loop (Figure 10F). We propose that the rigid-body rotation of domain 3 between the 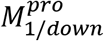 and 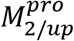 states is subserved by alternating de-solvation and re-solvation of polar and charged side chains and backbone groups (which, according to our WATMD results, are solvated by H-bond depleted water) positioned within the domain 2-3 interface (Figure 11). New H-bond depleted solvation appears in the wake of domain 3 rearrangements, such that the net reduction of unfavorable solvation free energy is never achieved in either configuration (see below).

**Figure 10.**
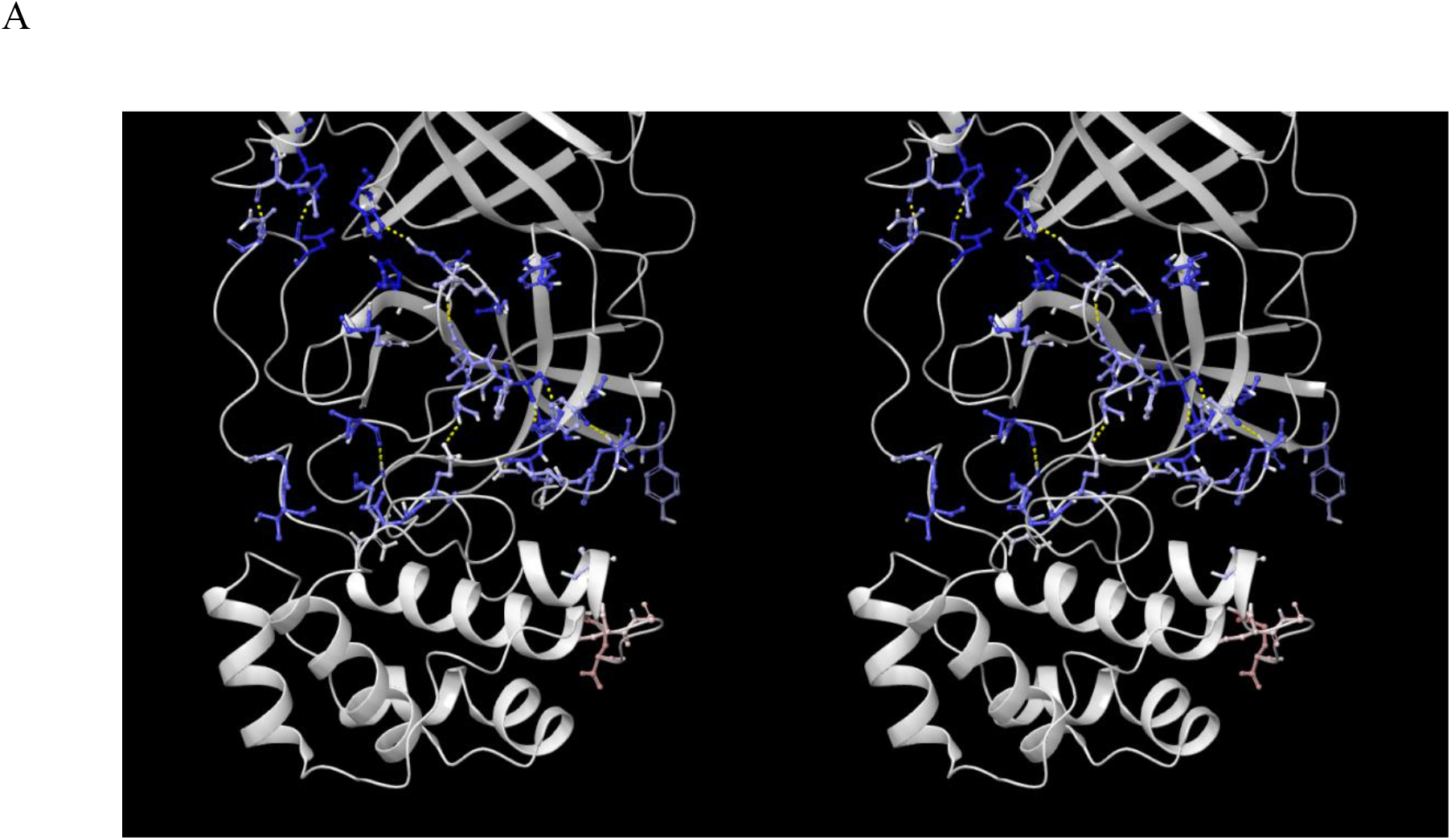
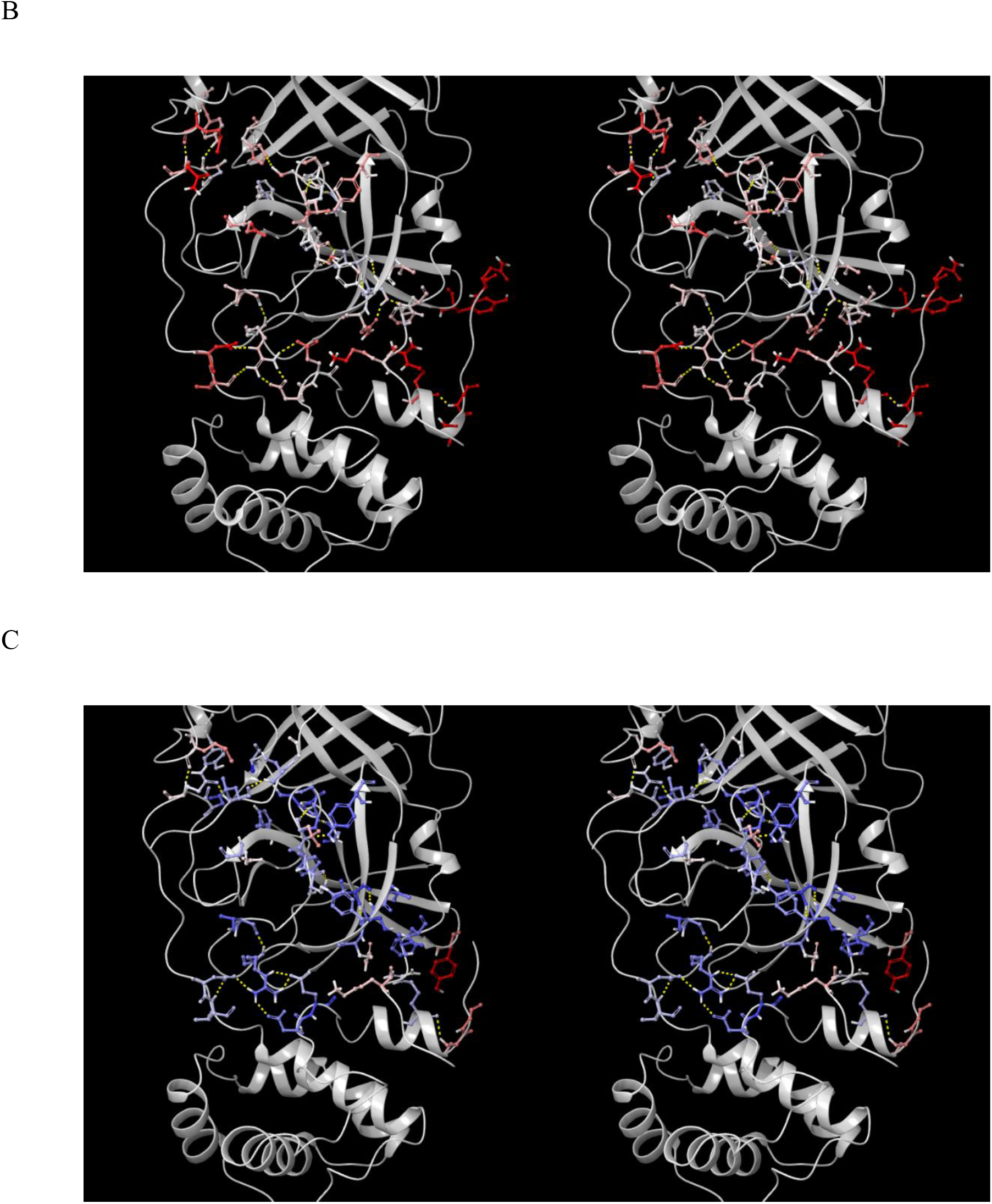
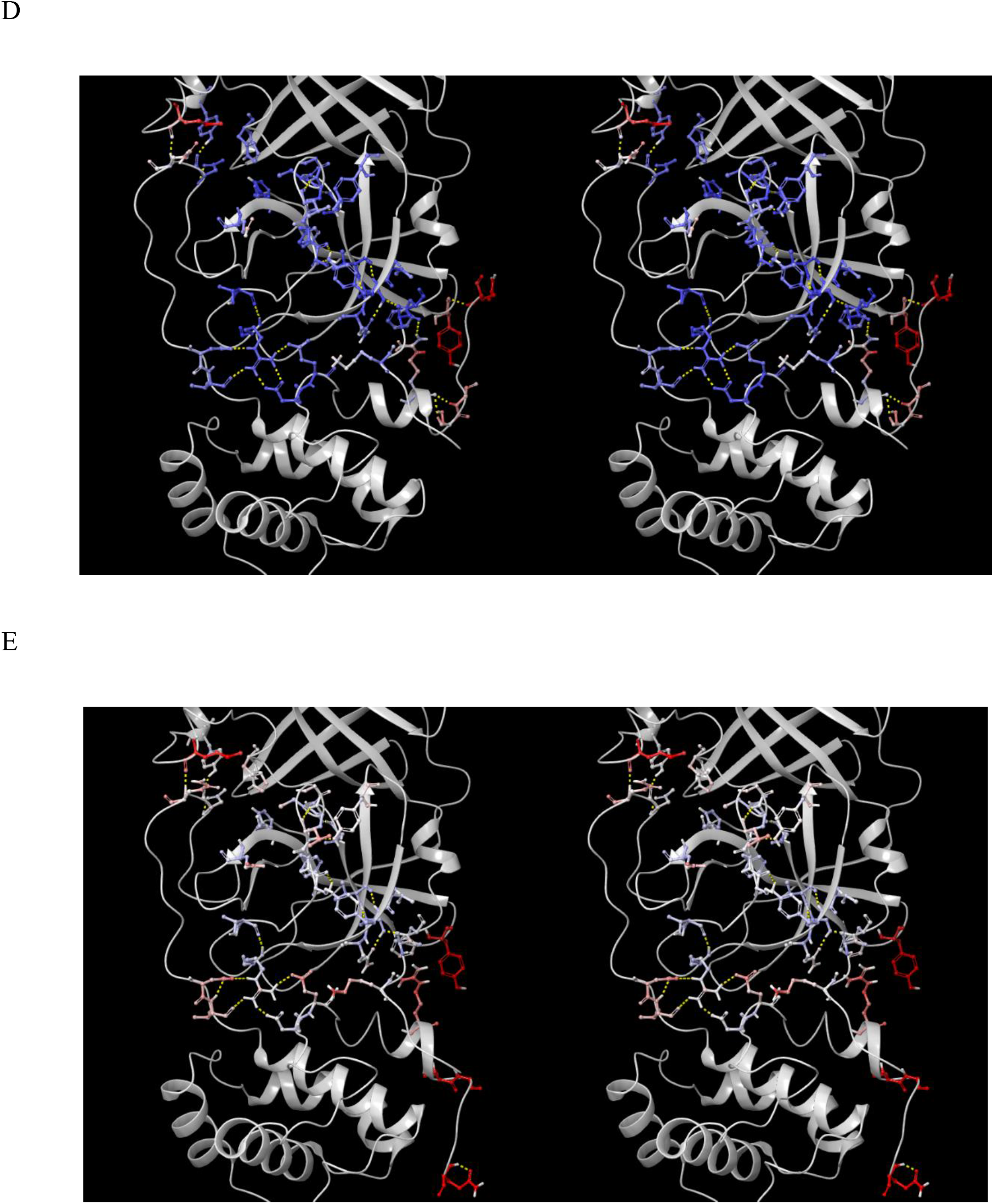
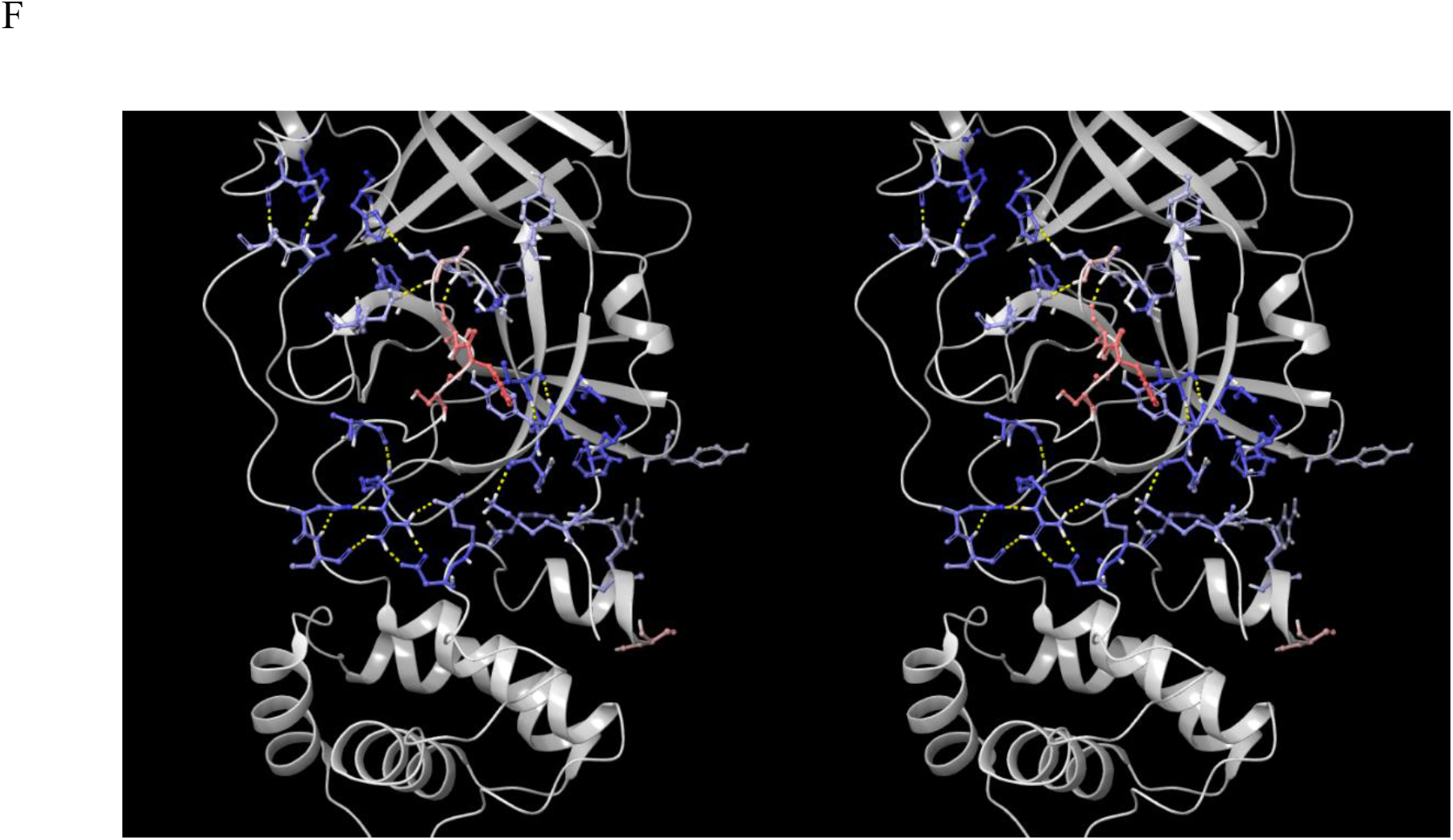
Stereo views of monomeric CoV M^pro^, together with a single chain extracted from selected dimeric structures as noted, showing the gross differences in the H-bond network governing the 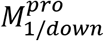 and 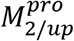 states. (A) The H-bond network in 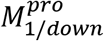 (2QCY) showing key residues color-coded by B-factor (blue ⟶ red color gradient depicting low to high values, respectively). (B) Same as A, except for a single chain of a representative apo 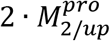 structure (6M03). We note the warmer B-factors are consistent with the higher energy state of the unbound dimer. (C) Same as A, except for a single chain of the substrate-bound 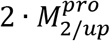 structure in 2Q6G. The cooler B-factors are consistent with the lower energy state of the bound dimer. (D) Same as A, except for a single chain of the inhibited boceprevir-bound 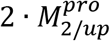 structure (PDB code = 6WNP). We note that the B-factors are somewhat cooler than those in the substrate-bound 2Q6G structure. (E) Same as D, except for the N3 inhibitor-bound 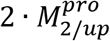 structure (PDB code = 6LU7). The B-factors are only slightly cooler than the apo dimeric structure, consistent with higher energy/lower binding affinity of this inhibitor. (F) Same as A, except for the protein captured in the 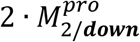 state (revisited in the following section).

**Figure 11.**
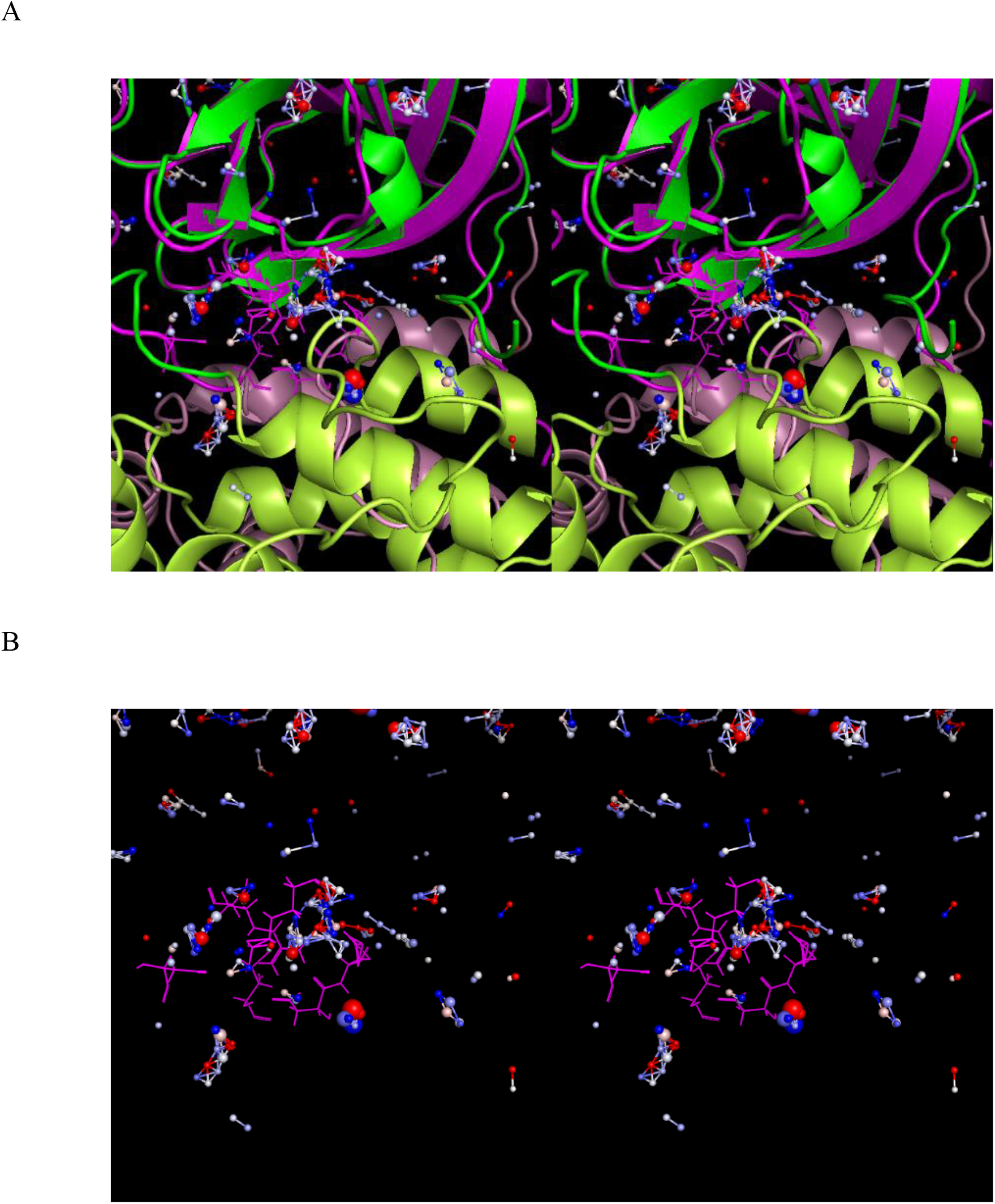
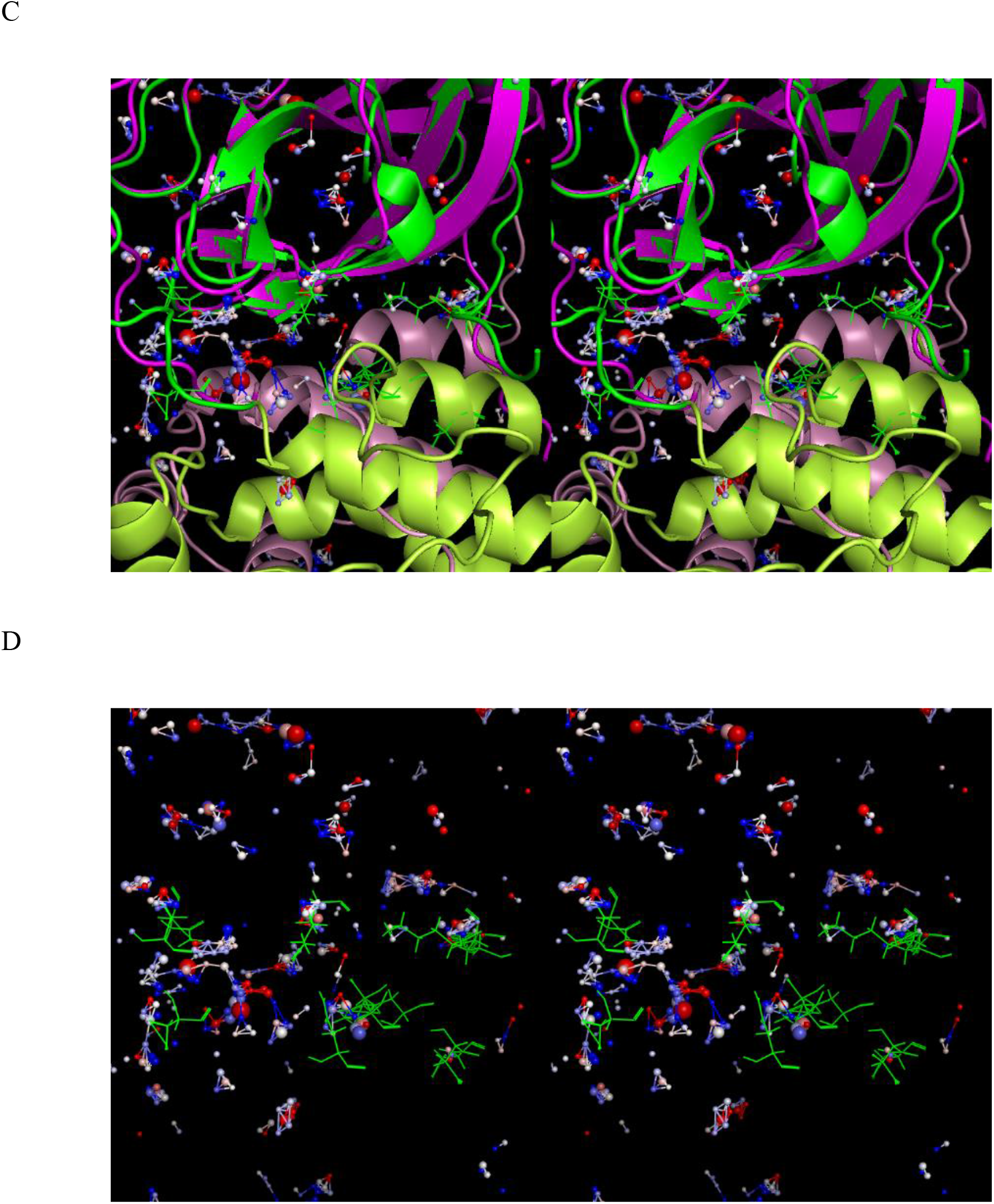
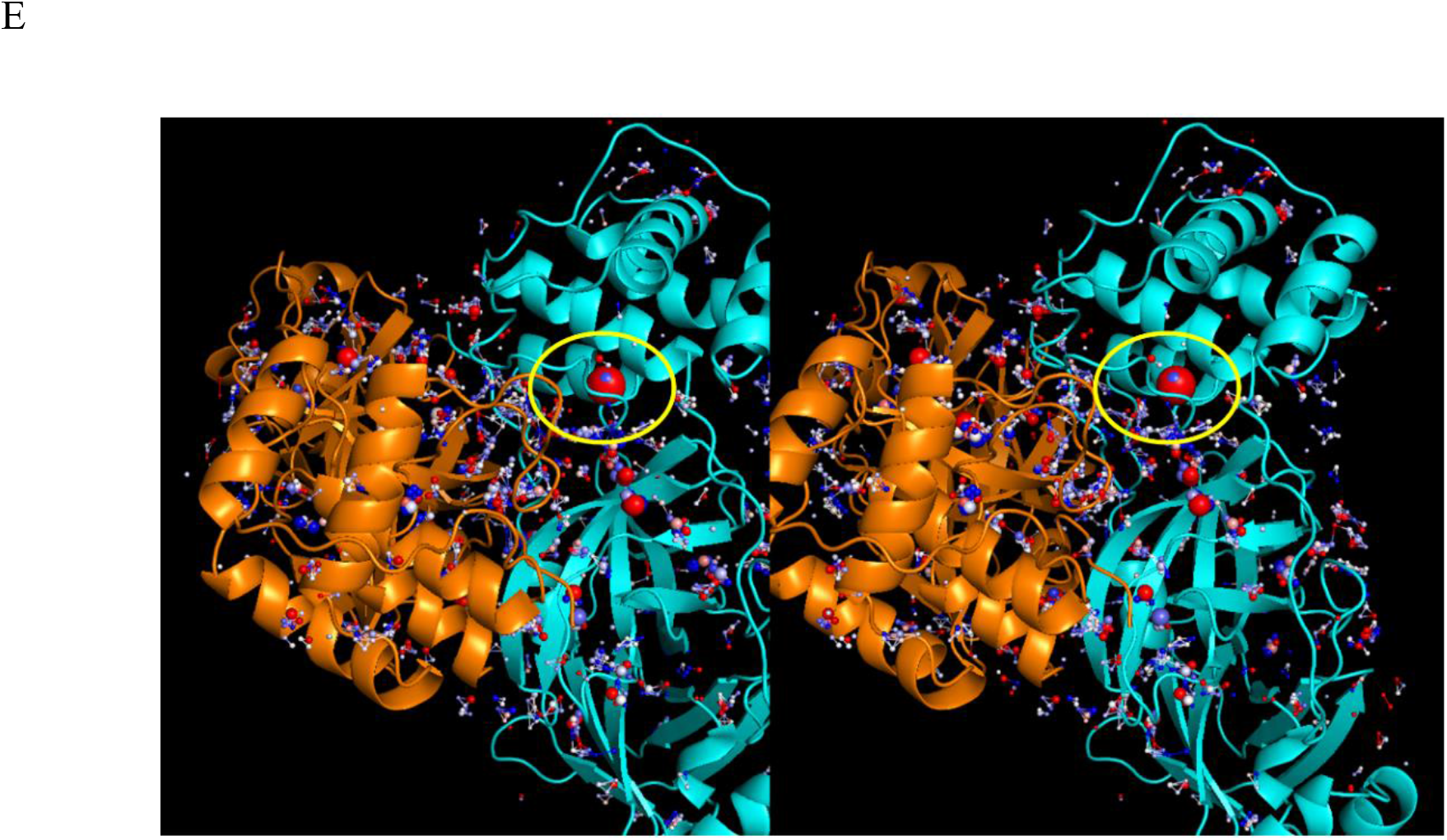
The WATMD-calculated solvation structures within the domain {1-2}-3 interface of <u>monomeric</u> 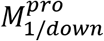 (2QCY) and a single subunit of 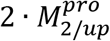 (2Q6G) (in which the time-averaged protein structures were aligned to the α-carbons). Rapid domain 3 rotation is expected in the absence of H-bond <u>enriched</u> solvation in the interface, whereas the t_1/2_ of each state depends on the total free energy magnitude of the H-bond <u>depleted</u> solvation expelled during state entry (which governs the re-solvation cost of exiting from each state), together with any electrostatic gains from charge-charge side chain interactions. (A) The time-averaged 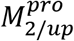 structure (domains {1-2} and 3 shown in magenta and salmon, respectively) overlaid on the solvation structure of 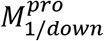 (domains {1-2} and 3 shown in dark green and light green, respectively). Water is expelled from regions of the 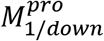 solvation structure that are occupied by side chain or backbone atoms upon entry to the 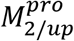 state. (B) Same as A, except showing only the side chain overlaps with the solvation hotspots. (C) The time-averaged 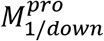 structure (domains {1-2} and 3 shown in dark green and light green, respectively) overlaid on the solvation structure of 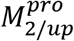 (domains {1-2} and 3 shown in magenta and salmon, respectively). Water is expelled from regions of the solvation structure of 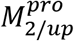 occupied by side chain or backbone atoms upon entry to the 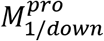 state. (D) Same as C, except showing only the side chain overlaps with the solvation hotspots. (E) The solvation structure within the domain {1-2}-3 interface of the dimeric time-averaged 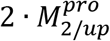 structure (6M03), looking into the space between the two helical domain 3 regions, with the domain {1-2} region of each chain toward the top and bottom of the drawing. The solvation structure in this case corresponds to the <u>residual un-expelled solvation</u>. The interface is solvated largely by H-bond depleted water, much of which is buried within the solvent-accessible surface. We postulate that this solvation is counter-balanced by a high occupancy H-bond enriched solvation hotspot (circled in yellow), in the absence of which, structural stability may be compromised.

#### Solvation-powered conformational transitions of the m-shaped loop

The m-shaped loop, which contains the catalytic Cys (resident on crest A) and oxyanion hole (resident on crests A and B), is common to all members of the chymotrypsin family, although the structural properties of these enzymes vary. Crest B of M^pro^ switches between the down (S1 pocket-accessible) (Figure 12A) and up (S1 pocket-inaccessible) positions (Figure 12B), corresponding to the 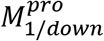 and 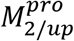 states of the enzyme, respectively. The S1 pocket switches between the open/oxyanion hole misaligned and closed/oxyanion hole aligned states in 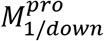 and in 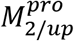, respectively. Although access to the S1 pocket is sterically blocked by Asn142 in the crest B up position, the cavity itself remains intact and occupiable (as such, Asn142 acts as a gatekeeper rather than a plug) (Figure 12C). We postulate that the complex m-shaped loop mechanism of M^pro^ is tailored for lowering the high de-solvation cost of the polar P1 Gln side chain during substrate association to the S1 pocket. The need for this mechanism is obviated in hepatitis C NS3 protease and chymotrypsin due to their preference for Cys/Thr and aromatic P1 side chains, respectively. As such, the m-shaped loops of these proteins are instead rigidified via an extra crest in NS3 (Figures 13A) and a disulfide bond to an adjacent chain in chymotrypsin (Figure 13B), resulting in permanent S1 pocket accessibility (Figures 13C-D).

**Figure 12.**
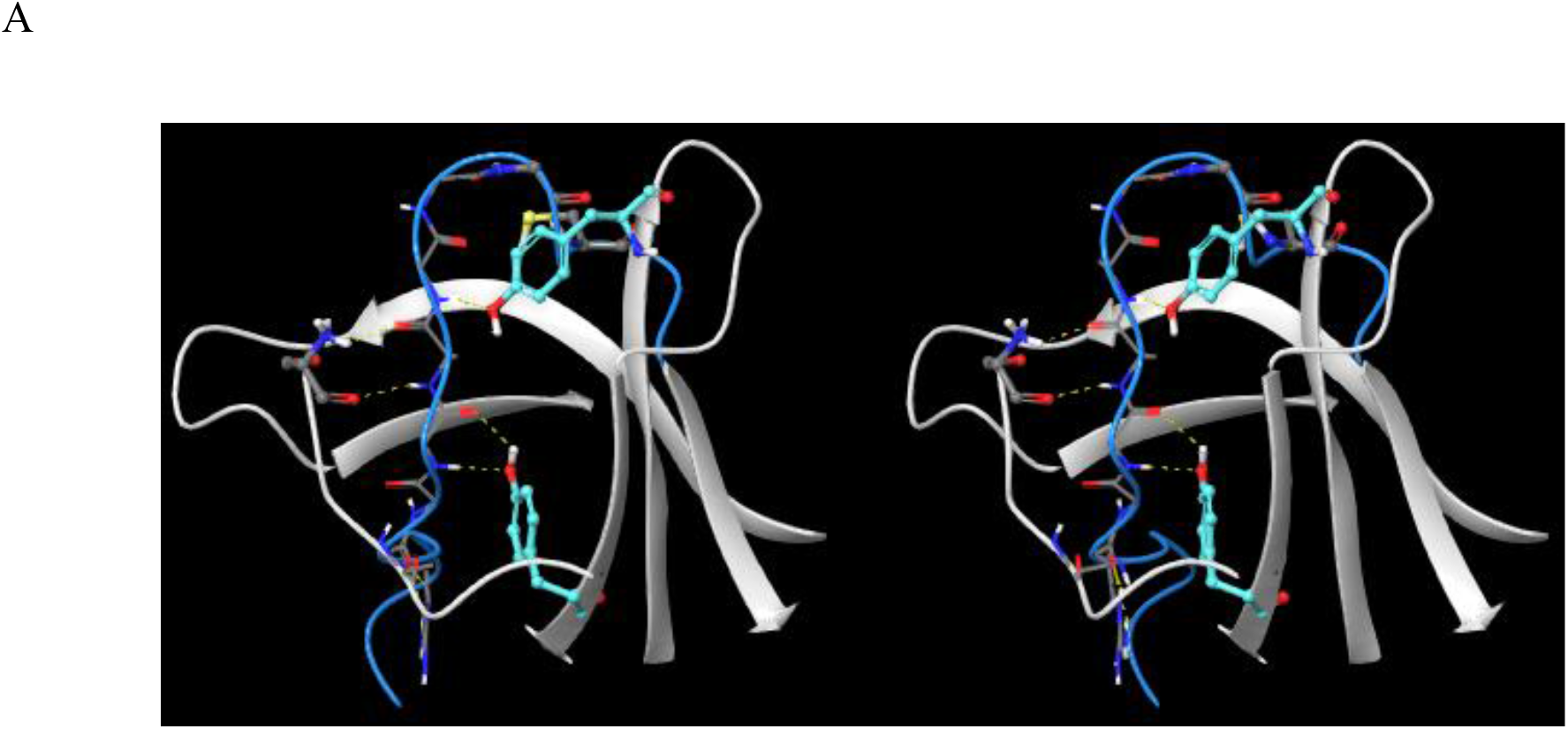
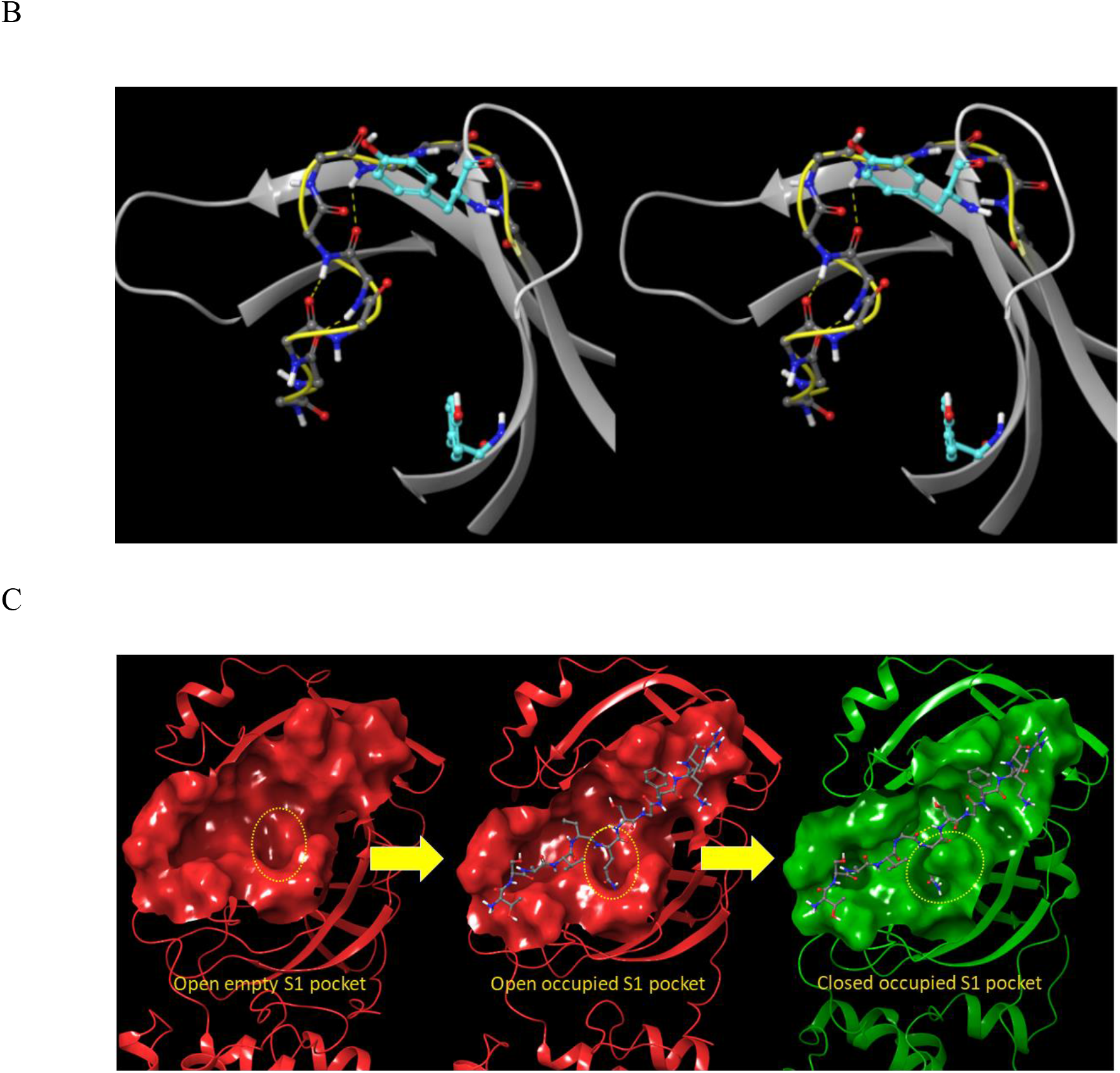
(A) Stereo view of the m-shaped loop of M^pro^ in the up state of crest B (blue). (B) Stereo view of the m-shaped loop of M^pro^ in the down state of crest B (yellow). (C) Left: unbound M^pro^ exists in the open state (corresponding to the down position of crest B), awaiting substrate association. Middle: substrates associate to the AS, projecting their P1 side chain into the open S1 pocket. Right: crest B undergoes substrate-induced rearrangement to the up position, thereby closing the S1 pocket (see text). We postulate that this mechanism facilitates partial de-solvation of the highly polar P1 Gln side chain of cognate M^pro^ substrates.

**Figure 13.**
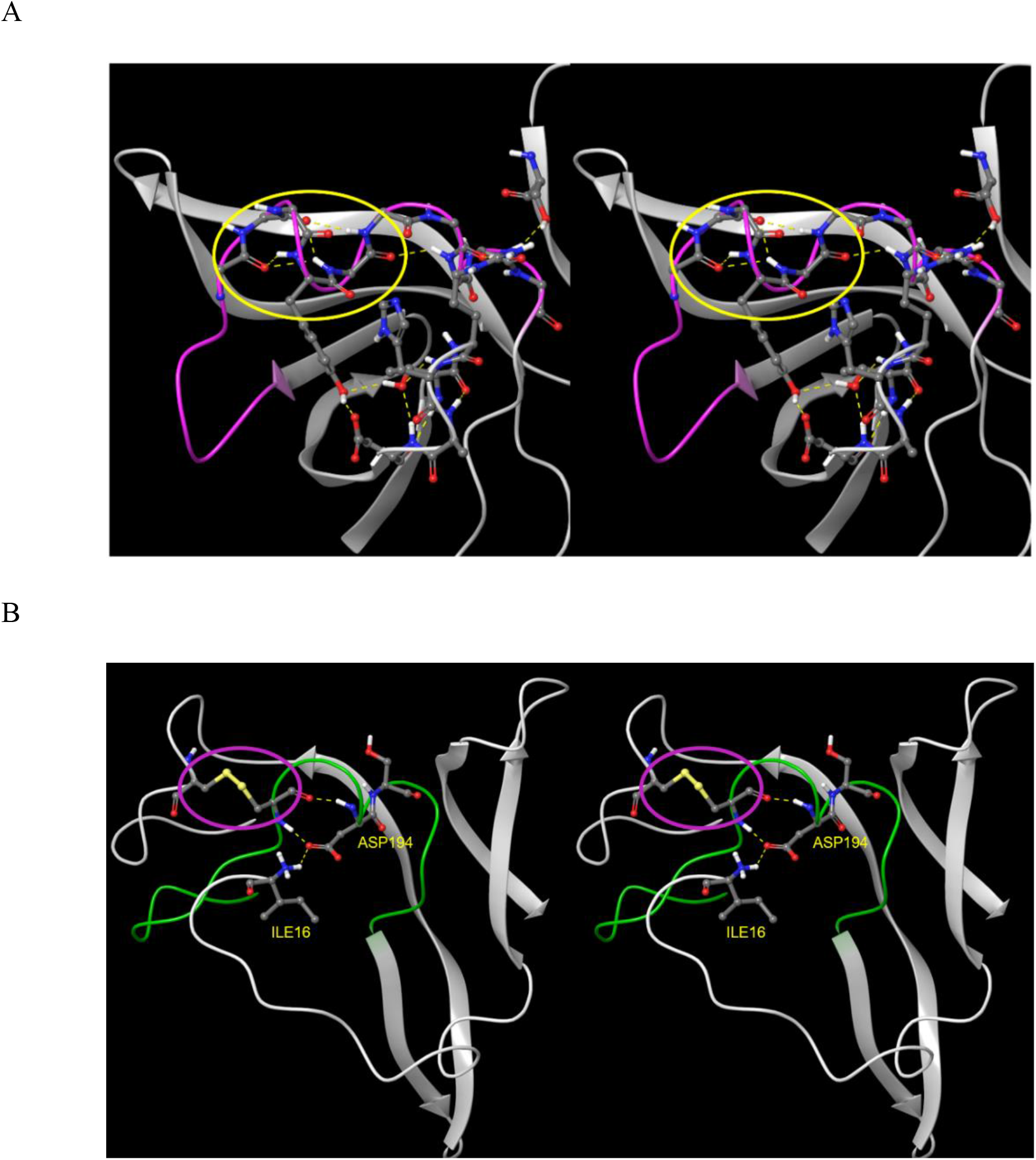
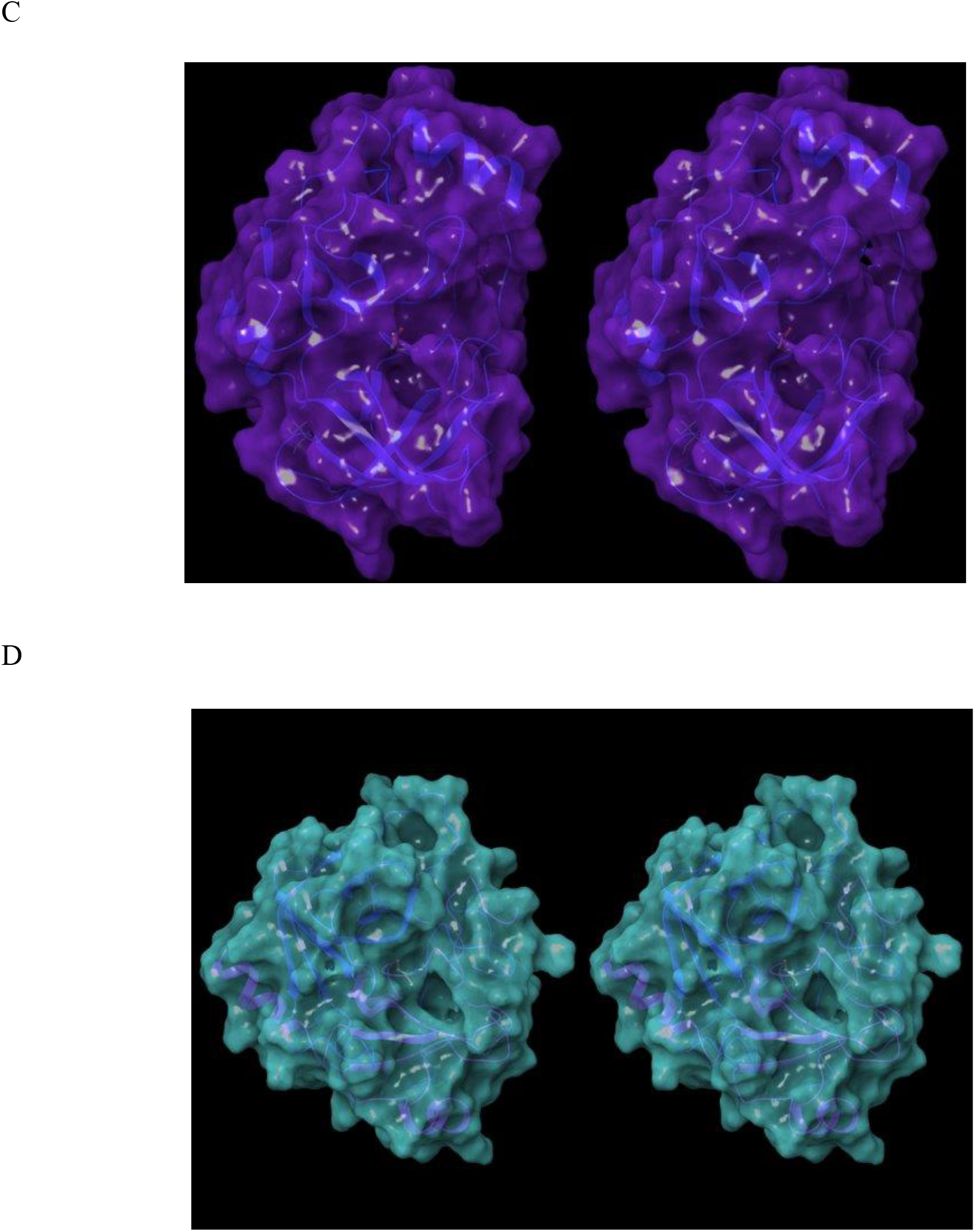
(A) Stereo view of the m-shaped loop of hepatitis C NS3 protease (PDB code = 4KTC). The loop (magenta) is stabilized by a third crest (circled in yellow), together with the H-bond network shown in the figure. (B) Stereo view of the m-shaped loop of chymotrypsin (PDB code = 4CHA). The loop (green) is stabilized by a disulfide bond in the rising leg (circled in magenta), together with H-bonds between backbone groups, and between Asp194 and the protonated N-terminal Ile16. (C) The S1 pocket is permanently accessible in NS3 protease, consistent with the lower de-solvation cost of the <u>Cys/Thr</u> P1 side chains of its cognate substrates. (D) Stereo view of the S1 pocket of chymotrypsin, which is permanently accessible, consistent with lower de-solvation cost of the <u>aromatic</u> P1 side chains of its cognate substrates.

We explored the 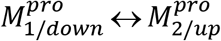 transition mechanism in M^pro^ by comparison of the monomeric and representative dimeric CoV and CoV-2 structures to better understand the functional purpose and detailed structural and energetic basis of the up/down bidirectional state transition of crest B. We now turn to:

1. The conformational properties of the m-shaped loop in the 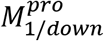 and 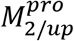 states.
2. The means by which m-shaped loop and domain 3 conformational dynamics are coupled.
3. The role of m-shaped loop conformational dynamics in governing the S1 pocket properties and P1 Gln de-solvation mechanism.

Next, we compared the detailed conformational properties of the rising leg of the m-shaped loop vis-à-vis crest B repositioning in representative crystal structures capturing 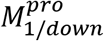 in 2QCY, 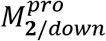 in 2BX3, and 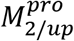 in 6WNP, noting that with the exception of 2QCY and 2BX3, the 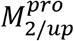conformations are highly similar across all CoV and CoV-2 structures. An overlay of the three structures reveals the existence of a similar 310 helix in 2QCY and 2BX3, despite the differing position of domain 3 in these structures (Figure 14A). The domain 3 position in 2BX3 is similar to that in 6WNP, but the m-shaped loop conformation is extended in the latter structure, and the Lys5-Gln127 H-bond in zone 3 that promotes the 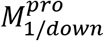 state is also present in the 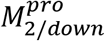 state, suggesting that the m-shaped loop conformation and domain 3 position are anomalously decoupled. These and other differences do not appear to be driven by pH differences, noting that boceprevir crystallized in CoV-2 M^pro^ at pH 6.5 (PDB code = 7BRP) and pH 7.5 in 6WNP exhibit only slight structural differences. A comparison of the m-shaped loops in 2QCY and 6WNP reveals the detailed differences between these two conformations (Figure 14B):

1. Tyr118 and Tyr126 (part of zone 3) in the extended conformation are respectively H-bonded to Leu141 and Ser139 on the rising leg of the m-shaped loop in the 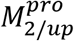 state, but not in the 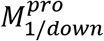 state.
2. The rising leg of the m-shaped loop contributes to the lining of the S1 pocket (addressed in the following section).
3. Glu290 (part of zone 2) is H-bonded to Ser139 on the rising leg of the m-shaped loop in the 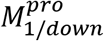 state, but not in the 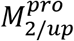 state.
4. The N-terminal basic group of Ser1 binds to the backbone oxygen of Phe140 in some structures, but not in others, suggesting that it plays little or no direct role in substrate binding.

**Figure 14.**
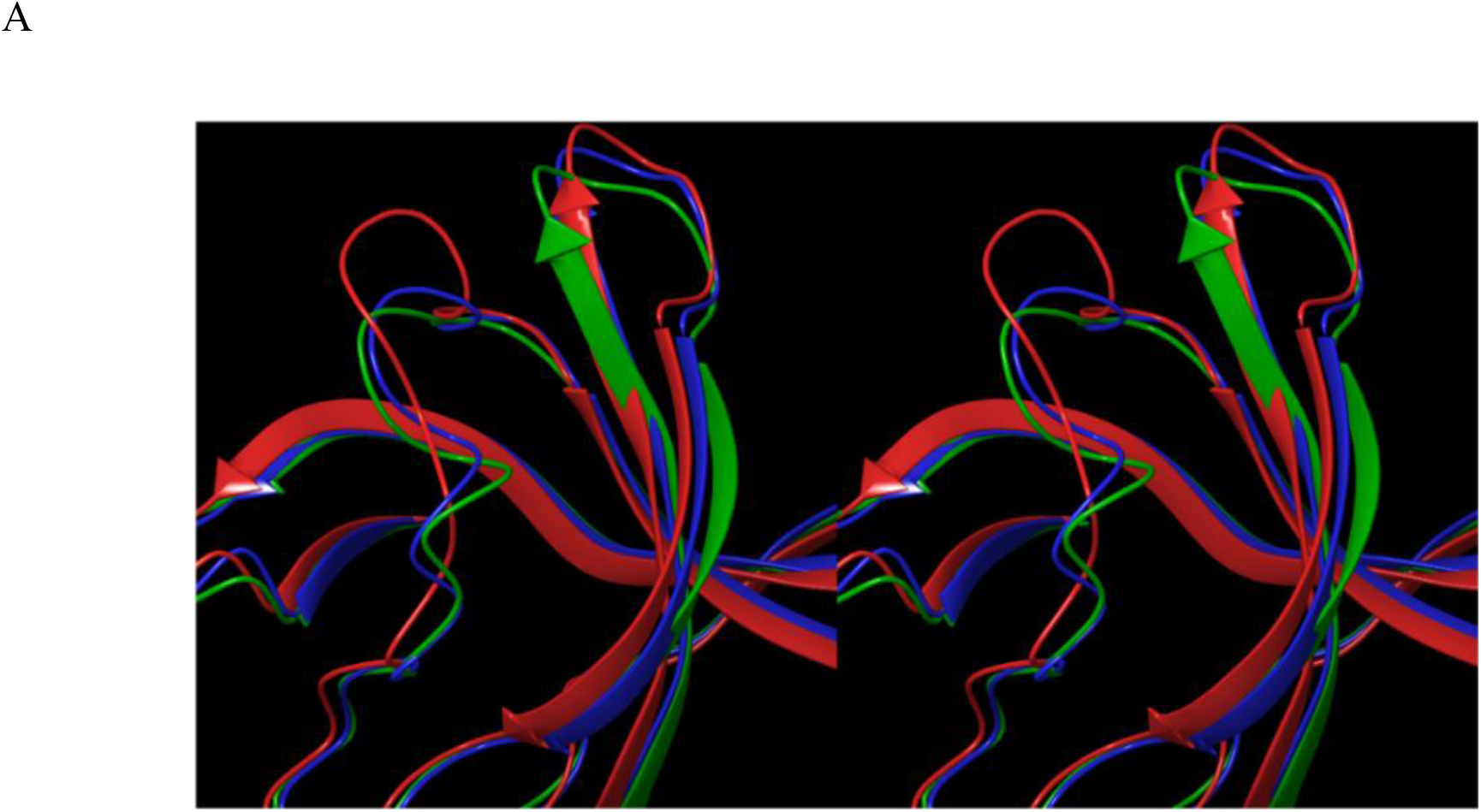
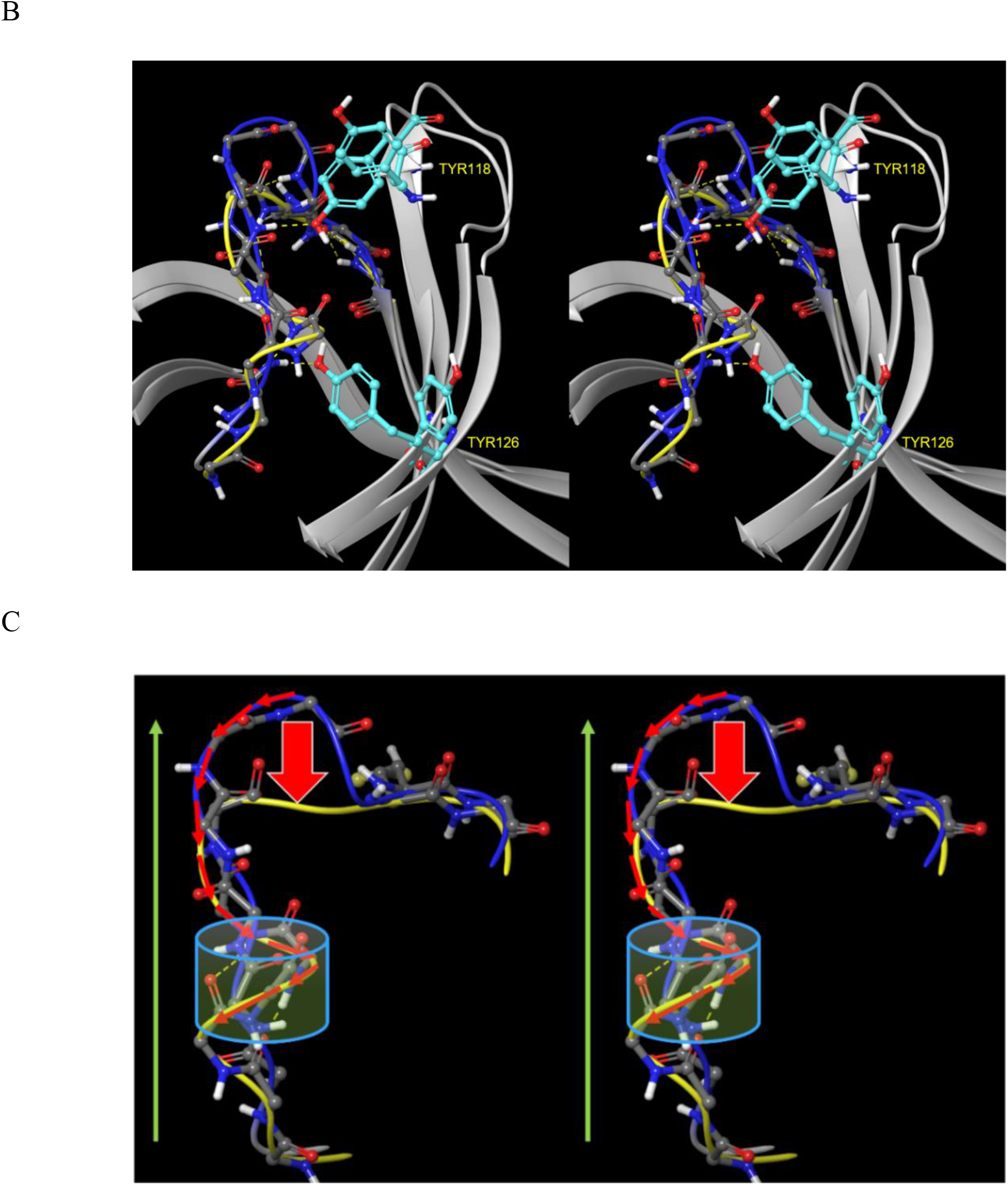
(A) Overlay of the m-shaped loops in the dimeric boceprevir-bound CoV-2 M^pro^ (6WNP, red), monomeric CoV M^pro^ (2QCY, green), and dimeric CoV Mpro (2BX3, blue). (B) Overlay of the m-shaped loop in the up (blue) and down (yellow) states of crest B. (C) The down state of crest B is generated (red block arrow) by reversibly spooling the more steeply sloped extended form (N-to C-terminal direction denoted by the green arrow) to/from the shallower 3_10_ helical turn.

Crest B down/up cycling is coupled directly to domain 3 repositioning, which together form the basis of the 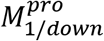 and 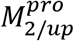 states. Crest B down/up transitions are subserved by 3_10_ helix ⟷ extended conformational transitions in the rising stem of the m-shaped loop, in which the extended chain “spools” in and out of the helical turn, respectively (Figure 14C).

Non-equilibrium energy flows, consisting of dynamic free energy barriers that increase and decrease in a state-dependent fashion (a key tenet of Biodynamics), are apparent in M^pro^. State transitions resulting in increased solvation free energy (i.e. increased barrier height) are slow, whereas those resulting in unchanged (or lowered) solvation free energy (i.e. decreased barrier height) are fast. The affected water molecules may reside anywhere within the solvation structure. The Goldilocks zone of protein stability (too high results in the loss of rearrangeability; too low results in aggregation/loss of structure) is achieved via counter-balancing between H-bond enriched and depleted solvation (a solvation free energy juggling act). We further note that in the absence of shielding, the cost of intra- and inter-protein H-bond rearrangements that depend on the expulsion/de-solvation of H-bond <u>enriched</u> water is typically a zero sum game, in which the fastest rearrangements prevail, and the lifetimes of which are governed by the <u>re-solvation</u> costs of exiting those states (including re-solvation of both H-bond depleted <u>polar</u> and <u>non-polar</u> positions). The specific nature of the aforementioned solvation juggling act is well-explained by the WATMD results for the 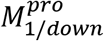 state (2QCPY) and the 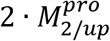 (6M03). The S1 pocket in both structures (denoted as S1_sol_), together with the region in 6M03 near Tyr118 occupied by the 3_10_ helical turn in the 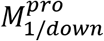 state, are solvated by H-bond depleted water (denoted as S1_sol_ and Tyr118_sol_) (Figure 15). Expulsion of Tyr118_sol_ via 3_10_ helix formation in the 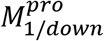 state is energetically preferred in the unbound S1 pocket, which avoids a “double dose” of unfavorable H-bond depleted S1_sol_ and Tyr118_sol_. However, expulsion of S1_sol_ via P1 group binding reduces the free energy cost to that of Tyr118_sol_, thereby facilitating the transition to the extended m-shaped loop conformation in the 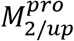 state. We postulate that expulsion of Tyr118_sol_ explains substrate-induced stabilization of the higher energy 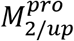 state (consistent with the lower B-factors of 2QCY compared with 6M03). Based on these results, we hypothesize that the 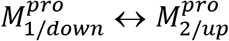 state transitions in the unbound protein are driven by energetic “frustration”, in which H-bond <u>depleted</u> solvation is gained transiently during the 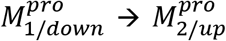 transition, while H-bond depleted solvation (S1_sol_) is expelled transiently from the S1 pocket of 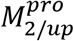 during substrate association and dimerization (see below). The solvation free energy is reset during dissociation of the final cleavage product, at which point the dimer dissociates, and 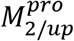 is no longer energetically favored over 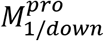.

**Figure 15.**
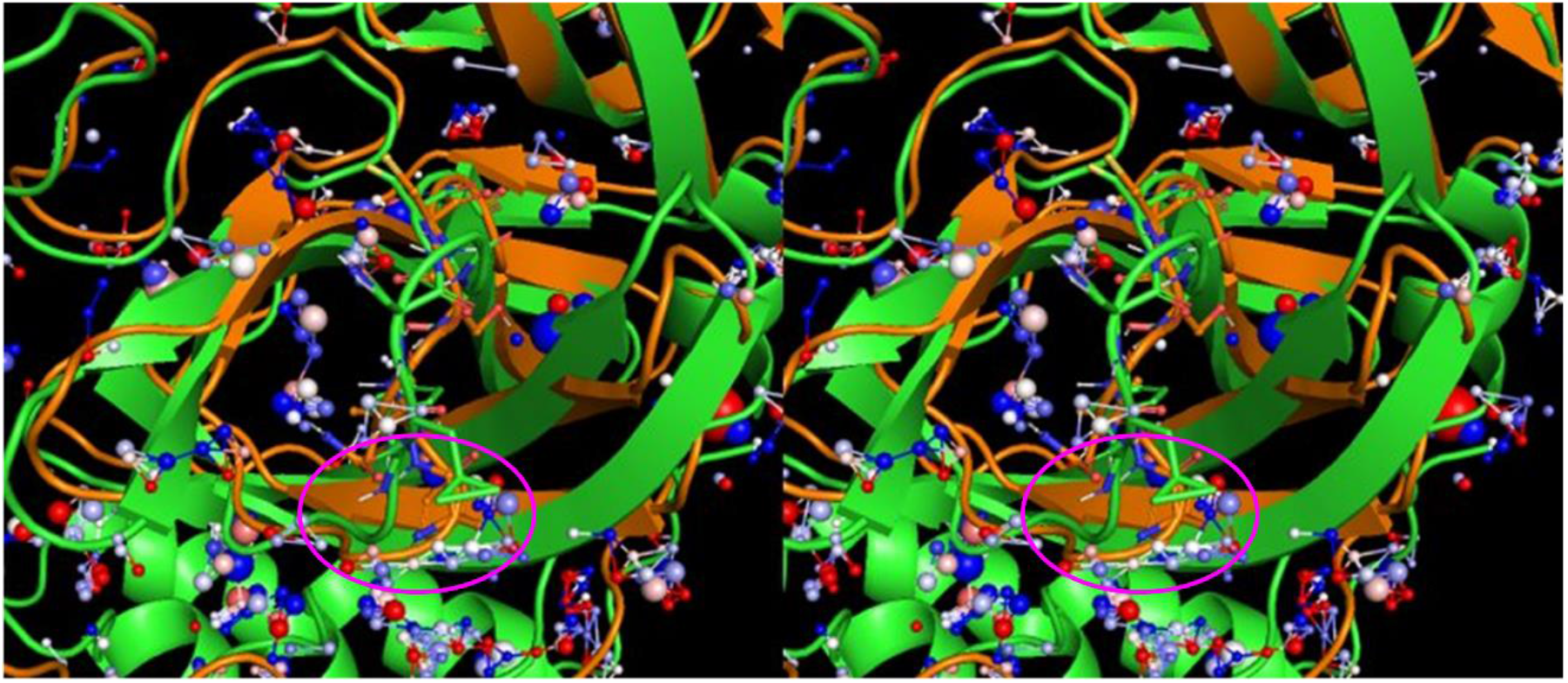
Stereo view of 2QCY (orange) overlaid on the calculated solvation hotspots of 6M03 (green), zoomed in on the rising leg of the extended m-shaped loop in 6M03 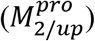 and the 3_10_ helix of 2QCY 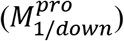. An arc-shaped region of low occupancy/H-bond depleted water (denoted as Tyr118_sol_) is present in the solvation structure of 6M03, which overlaps with the 3_10_ helical turn in 2QCY (circled in magenta). The stability of the 3_10_ helical conformation is enhanced by expulsion of this water during the 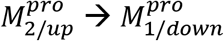 state transition.

### Enzyme dynamics in cis

Dimer-independent catalytic activity of <u>pre-cleaved</u> M^pro^ was observed by Chen et al., who nevertheless proposed the existence of an “intermediate” dimeric form of the enzyme [33]. A more plausible explanation is that pre-cleaved M^pro^ exists exclusively as monomers embedded within the polyprotein, whereas the post-cleaved species necessarily exists as a mixture of monomers and dimers, in which the monomeric form binds substrates that are cleaved by the dimeric form (such that 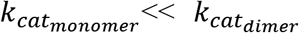. The <u>pre-cleaved</u> monomeric form of M^pro^ cannot be fully represented in 2QCY because the C-terminal peptide is spatially far from the AS (noting that the Gln306 C-terminus serves as the P1 residue of the pre-cleaved protein). We propose the existence of two distinct forms of monomeric M^pro^, consisting of:

1. The <u>post-cleaved</u> species captured in 2QCY.
2. An alternate <u>pre-cleaved</u> polyprotein-embedded form, in which the C-terminal peptide (Gln 276 and Gln306) of domain 3 is unfolded, with:

a. The cleavage peptide projecting into the AS (which likely precludes cleavage of pre-cleaved M^pro^ by post-cleaved M^pro^ in trans).
b. The remainder of the polyprotein exiting from the prime side of the pocket (noting that M^pro^ folding likely occurs after nsp4 cleavage).

We postulate that cis cleavage is facilitated in the 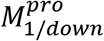 state, in which domain 3 is rotated toward the AS, and the C-terminal region of this domain (including the CTT helix) is partially unfolded (Figure 16). In the absence of this helix, the NTL is free to adopt the active Lys5-Gln127 H-bond-disrupted state that exists in all 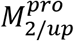 structures (i.e. a hybrid 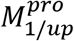 state).

**Figure 16.**
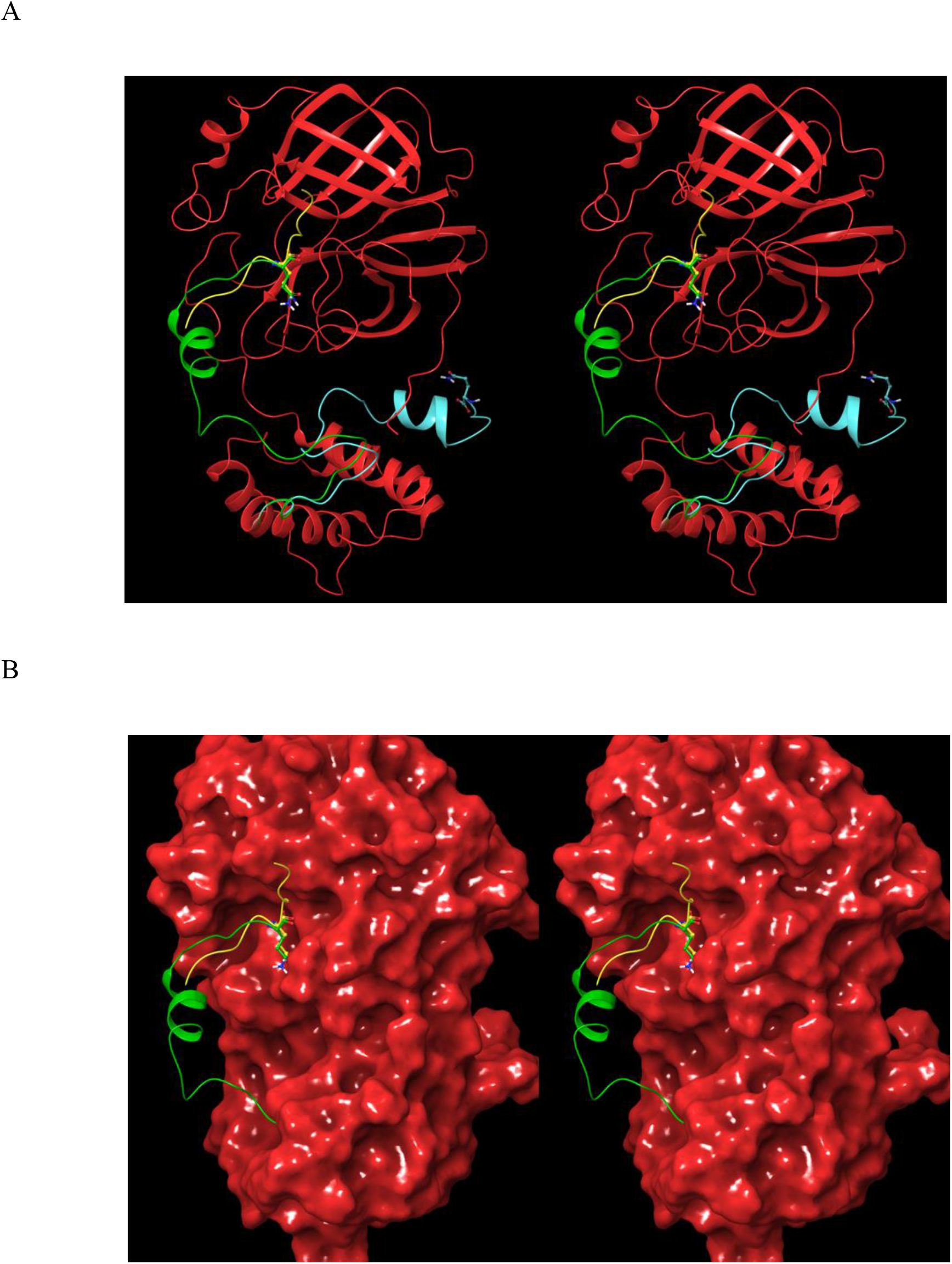
<u>Hypothetical</u> qualitative model of the cis cleavage structure of monomeric M^pro^ subsequent to turnover, in which the partially unfolded domain 3 of 2QCY projects into the AS (the P1 Gln306 side chain is shown for reference). (A) The modeled C-terminal region (green) extends from domain 3 to the AS. The cognate substrate extracted from 2Q6G (yellow) is overlaid on the modeled structure. The original C-terminal chain in 2QCY is shown in cyan. (B) Same as A, except showing the solvent-accessible surface.

### Inter-molecular rearrangements

#### Solvation-powered enzyme dynamics in trans

The M^pro^ catalytic cycle depends integrally on the dynamic intra-molecular rearrangements described above. We propose that substrates bind to monomeric M^pro^ in the 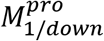 state, which upon transitioning to the 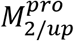 state, is further stabilized by the bound substrate in the catalytically active 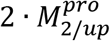 dimeric form. This process is accompanied by additional rearrangements, including switching of:

1. His172 (on the β-hairpin) from a non-H-bonded position (or Glu166-paired position in some structures) to a small H-bond network around the backbone oxygen of Ile136 in the 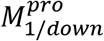 and 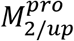 states, respectively.
2. His163 from an H-bonded position with Ser144 to the substrate P1 Gln side chain in the 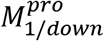 and 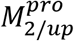 states, respectively.
3. Met165 between two alternate positions, both of which are observed as alternate side chain rotamers in several crystal structures. The S2 pocket is alternately blocked and unblocked in the two rotamers, suggesting that the rate of repositioning may be rate-limiting for cognate substrate binding (i.e. the Met165 side chain is energetically “frustrated”).

The catalytic cycle is energetically self-consistent, beginning with substrate association-induced expulsion of H-bond depleted solvation from the AS (see below) and gain of H-bond depleted solvation at Tyr118_sol_ (see above). Cleavage of the Gln306 peptide bond (Figure 17A) results in two products, consisting of the C- and N-terminal leaving groups (the pre-cleavage form bound to 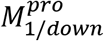 and 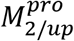 is shown in Figure 17B and 17C, respectively), noting that the chain inserts into the AS in the N-to C-terminal direction. Dissociation of the C-terminal leaving group has no impact on the intra-molecular/dimeric state of M^pro^ (Figure 17D), whereas that of the N-terminal leaving group resets the enzyme to the monomeric 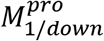 state (Figure 17E).

**Figure 17.**
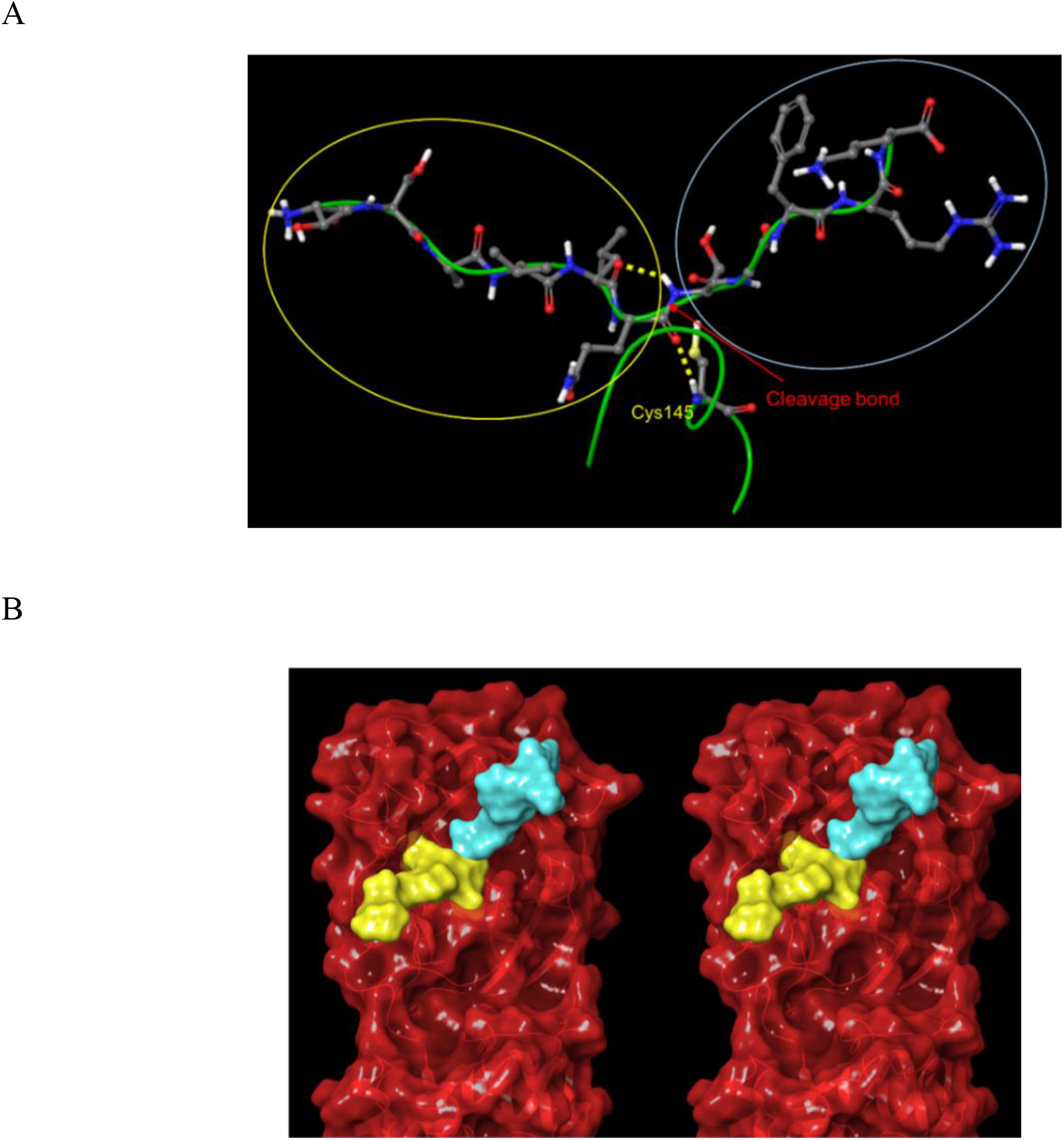
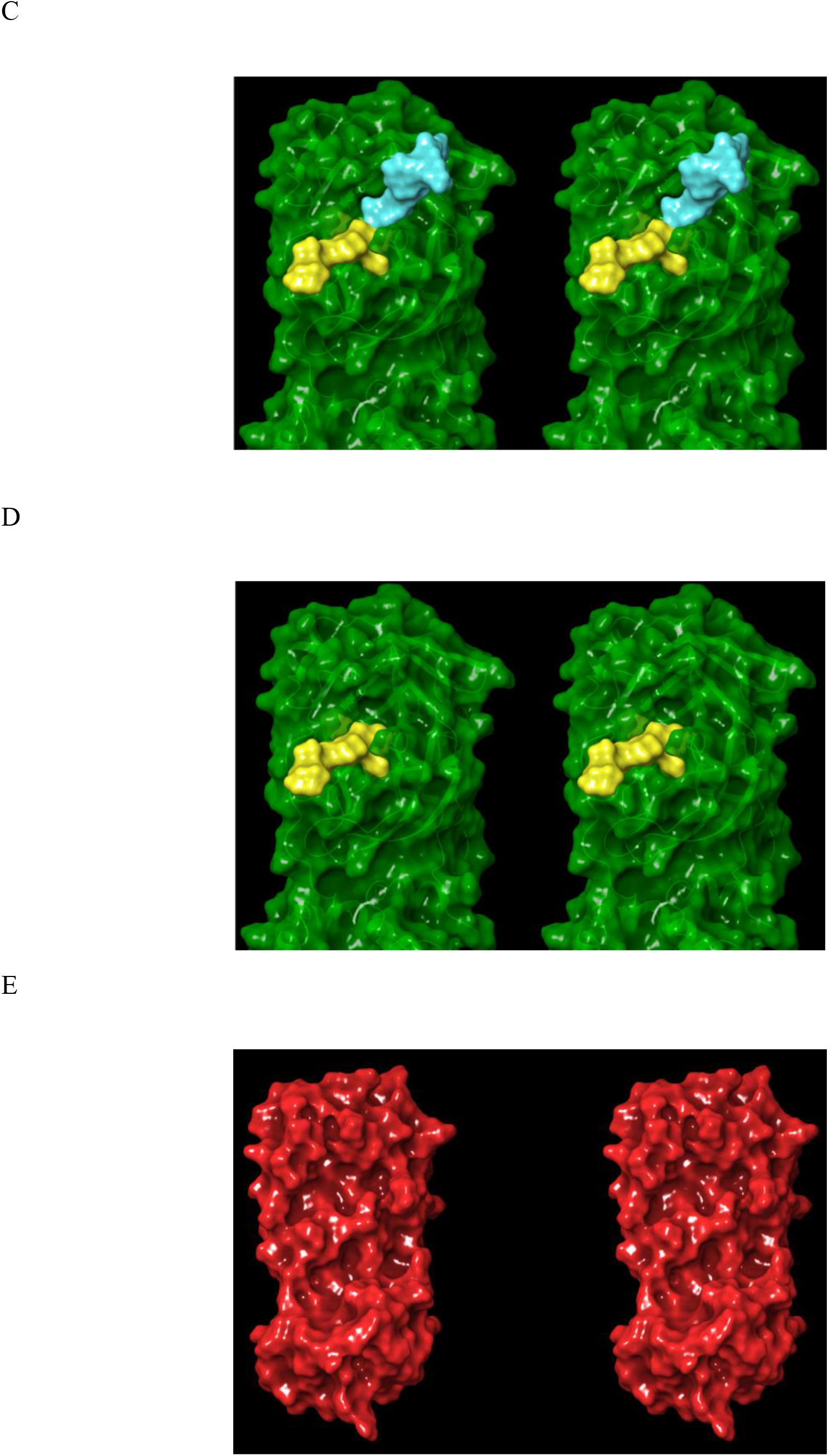
Stereo views of the proposed dynamic enzyme cycle. (A) The cognate CoV M^pro^ substrate from 2Q6G is divided into two zones around the cleavage bond (red arrow). The N- and C-terminal products are circled in yellow and blue, respectively. Cys145 is shown for reference. (B) The modeled substrate-bound structure in the 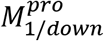 state (overlay of the substrate from 2Q6G on 2QCY) subsequent to association. (C) The substrate-bound structure in the 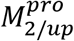 state in 2Q6G (single chain shown for clarity). (D) Same as C, except subsequent to dissociation of the C-terminal product. (E) Same as D, except subsequent to dissociation of the N-terminal product, at which point the dimer dissociates to the oscillating 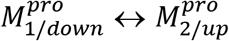 monomeric form. The protein population is unequally distributed among the monomeric and dimeric substrate-bound and unbound forms, each of which is further distributed among the 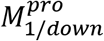 and 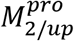 states (with the exception of dimers, which do not exist appreciably in the 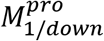 state).

The S1 pocket is comprised of the residues shown in Figures 18 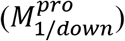 and 19 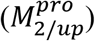, together with the substrate P3 side chain. A subset of these residues plays a dual role in substrate binding (via the backbone of Glu166) and:

1. Coupling the m-shaped loop to zone 3 (the backbone groups of Ser139 and Leu141) and zone 2 (Ser139 and Glu290) of the H-bond network, thereby destabilizing 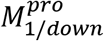 in the dimeric state.
2. Closing the S1 pocket via the crest B down ⟶ crest B up transition, which repositions the Asn142 gatekeeper over the pocket. We postulate that the cost of de-solvating the polar amide group of the P1 Gln side chain is reduced via this mechanism, such that the side chain binds with its solvation partially intact (noting that the S1 pocket is fully open in the 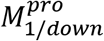 state (Figure 18), whereas the side of the pocket remains open in the 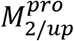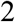 and 2 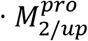 states (Figure 19)).
3. Orienting the scissile bond toward the attacking Cys145 side chain.

**Figure 18.**
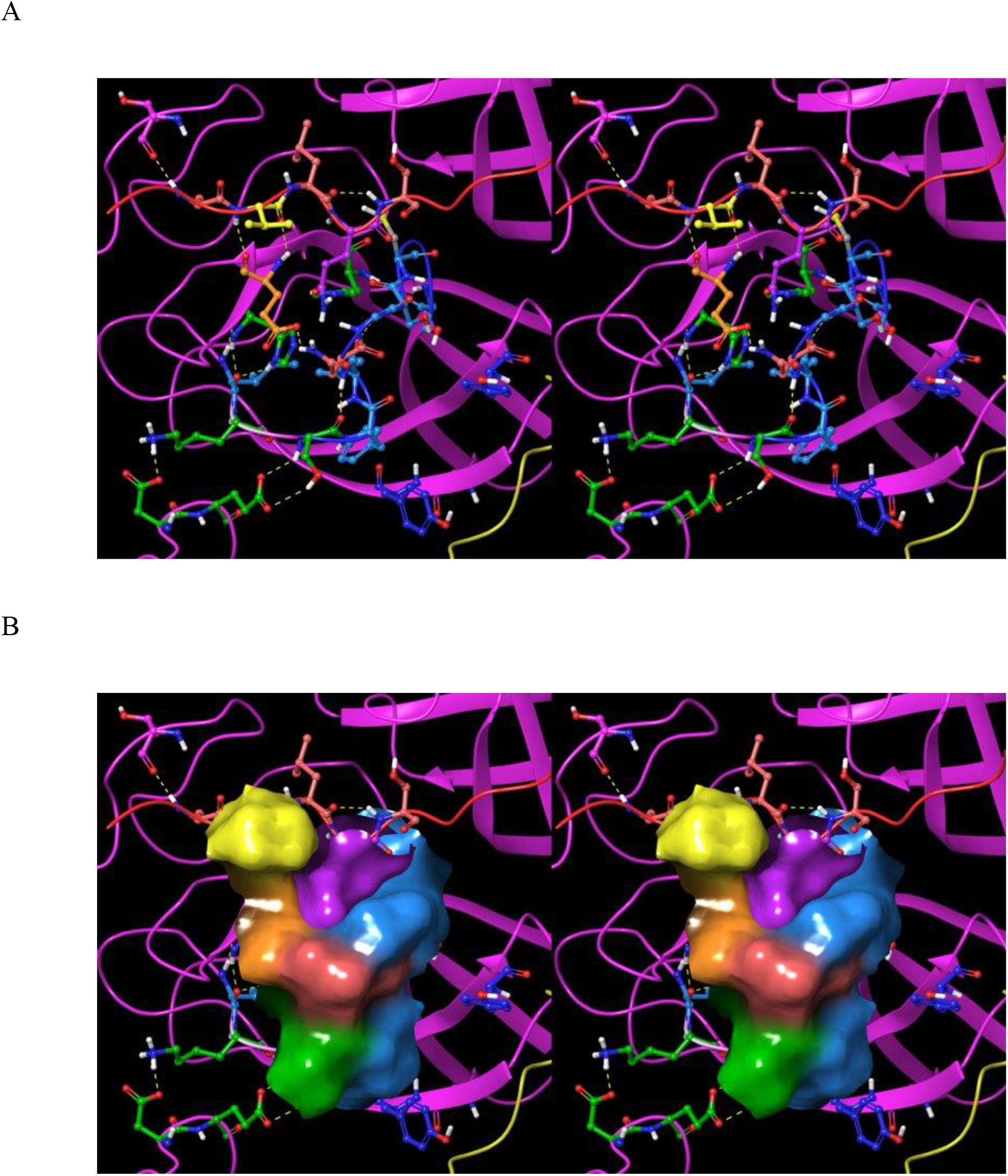
Stereo views of the S1 pocket in the 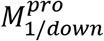 state in 2QCY with the bound substrate P1 group modeled in from 2Q6G. The substrate peptide (red ribbon) is visible at the top of the image. (A) The S1 pocket is lined by Glu166 (orange), His172 (green), His163 (not visible), Ser139 (blue), Phe140 (blue), Leu141 (blue), Asn142 (coral), and the substrate P3 side chain (yellow). The pocket is occupied by the P1 Gln side chain (pink). Many of the residues lining the pocket play dual roles: the backbone of Glu166 H-bonds with the substrate P3 backbone (thereby directly connecting the β-sheet formed by the substrate and β-hairpin to the S1 pocket). (B) Same as A, except showing the solvent-accessible surface (noting that the accessibility of the S1 pocket is underestimated by the solvent-accessible surface).

**Figure 19.**
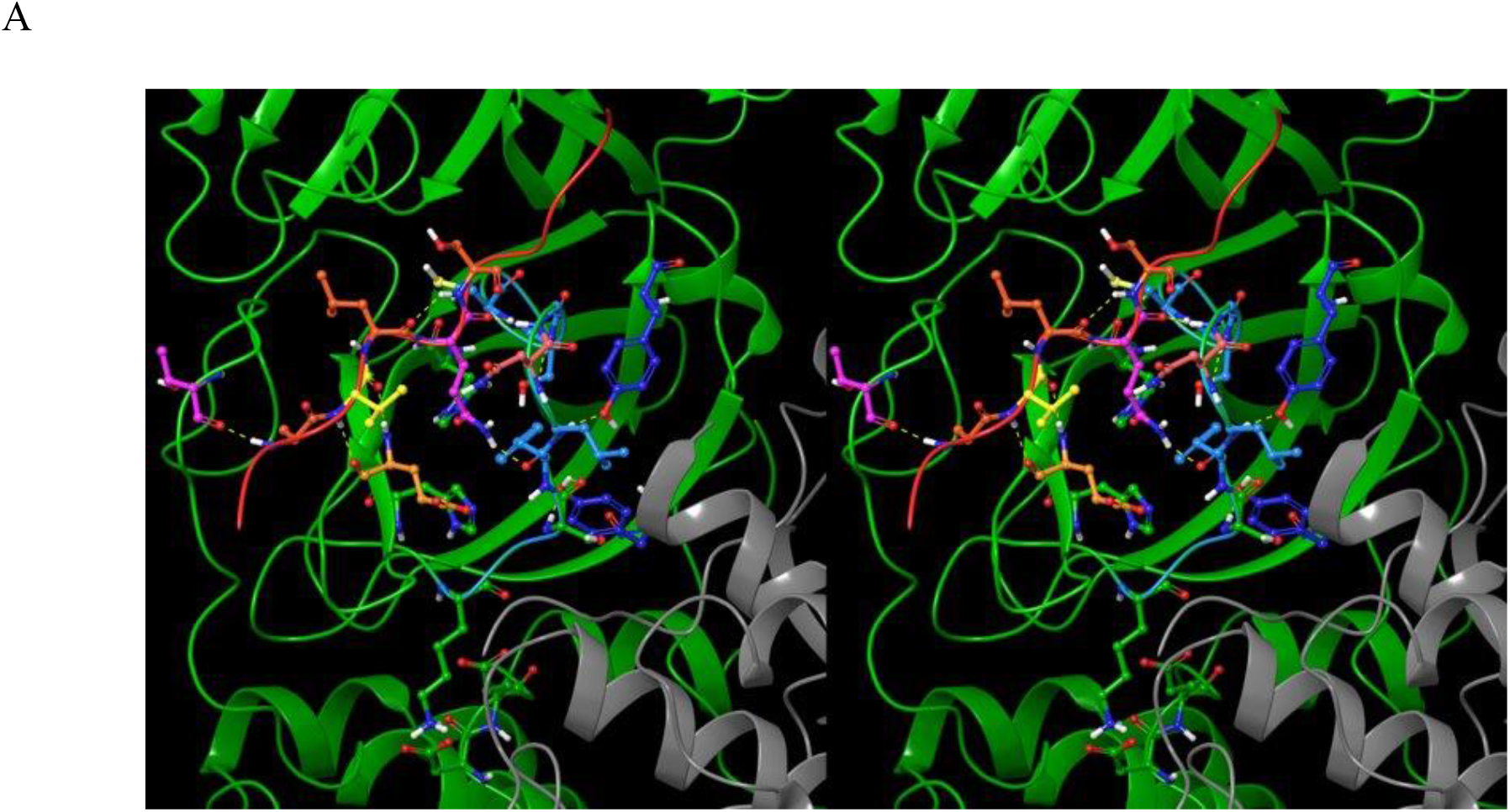
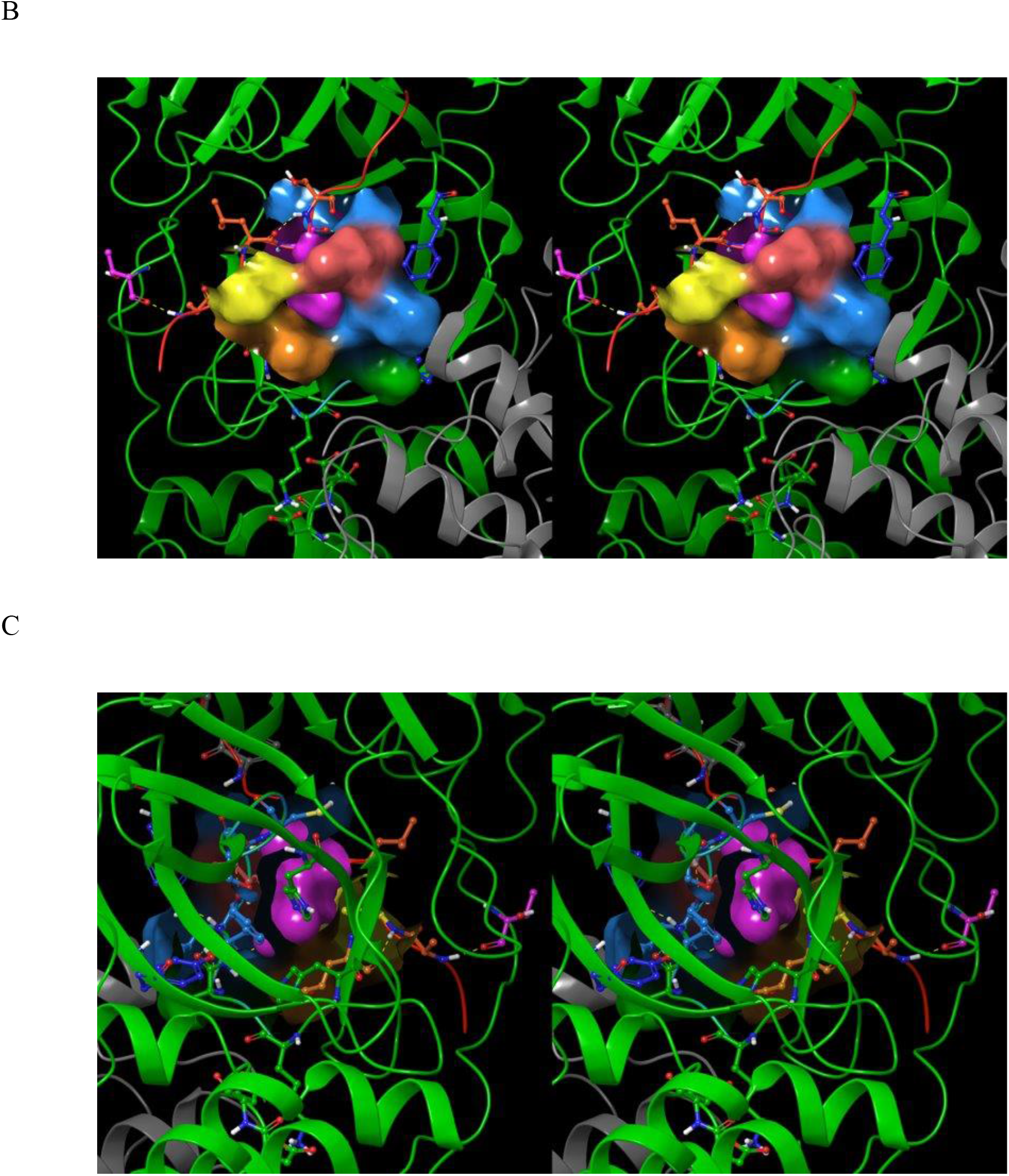
Stereo views of the S1 pocket in the 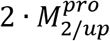 state and the bound substrate P1 group in 2Q6G. The substrate peptide (red ribbon) is visible at the top of the image. (A) The donut-shaped S1 pocket is lined by Glu166 (orange), His172 (green), His163 (not visible), Ser139 (blue), Phe140 (blue), Leu141 (blue), Asn142 (coral), and the substrate P3 side chain (yellow). The P1 Gln side chain (pink) occupies the donut hole, with the open side serving as a solvent-accessible cavity for the Gln amide, thereby reducing the de-solvation cost of this group. Many of the residues lining the pocket play dual roles: the backbone of Glu166 H-bonds with the substrate P3 backbone (thereby directly connecting the β-sheet formed by the substrate and β-hairpin to the S1 pocket) and Asn142 serves as the gatekeeper of the pocket. Tyr118 (zone 3) H-bonds with the backbone NH and oxygen of Ser139, and Tyr126 (zone 3) H-bonds with the backbone oxygen of Phe140. (B) Same as A, except showing the solvent-accessible surface lining the S1 pocket (noting that the pocket entrance is occluded by Asn142 and the substrate P3 group). (C) Same as B, except showing the rear side of the S1 pocket.

Access to the S1 pocket is blocked by Asn142 in the extended 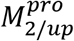 state (Figure 19A), which is pointed away in the 3_10_ helical 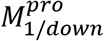 state (Figure 18A). As such, we postulate that substrates bind to the 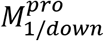 state, which then rotates about the domain 2-3 linker into the substrate-stabilized 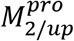 state, followed by dimerization.

#### Solvation-powered dimerization

Dimerization is widely assumed to govern both the activation and substrate complementarity of M^pro^ [34]. The dimer interface bridges the H-bond networks within the individual subunits via their NTL chains (Figure 20). The native dimer interface is well-explained by our WATMD results derived from a single monomeric chain extracted from the dimeric substrate-bound structure in 2Q6G. We explored the complementarity between non-bulk solvation positions and occupancy by atoms on the opposite subunit <u>prior to dimerization</u> via an overlay between the original dimeric structure (representing the inserting subunit) and the opposite subunit plus the solvation structure thereof. Expulsion of solvating water during dimerization is expected in regions where side chain/backbone atoms of the inserting subunit overlap with the solvation structure of the reference subunit. The results clearly demonstrate that the NTL of the inserting subunit (principally backbone atoms augmented by some side chains) is highly complementary to substrate-induced H-bond depleted solvation positions, and is therefore, a key driver of dimerization (noting that the sub-μM and single-digit μM dimerization K_d_ for the substrate-bound and free monomer, respectively [35,36] suggests that the solvation expelled from the interface is only weakly H-bond depleted) (Figure 21). Removal of the NTL results in an alternate tail-tail dimer interface about domain 3 of the member subunits [37]. An additional unfavorable free energy contribution, consisting of entropic loss due to the more restricted motion of domain 3 about the 2-3 linker in the dimeric form, is conceivable.

**Figure 20.**
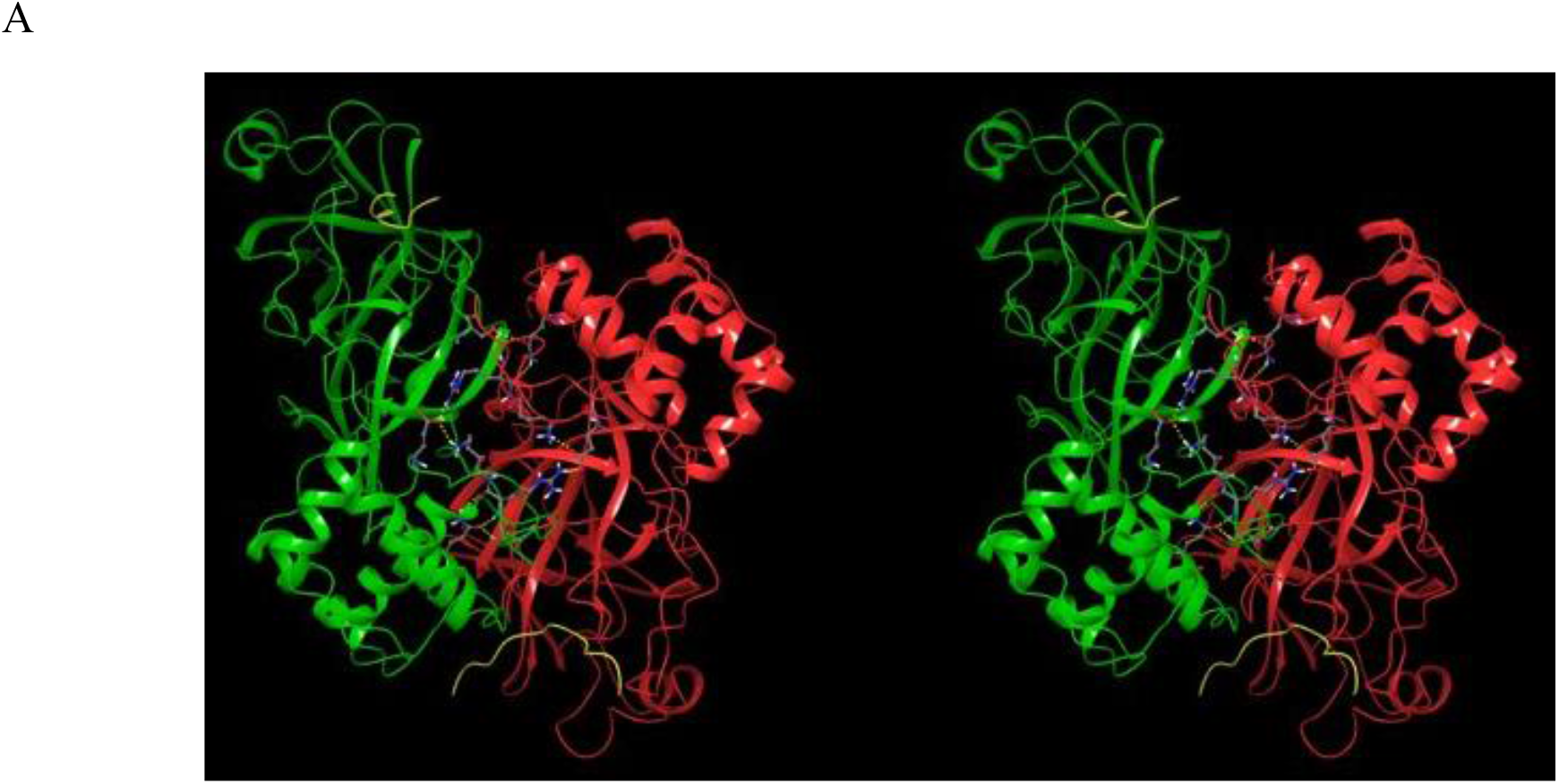
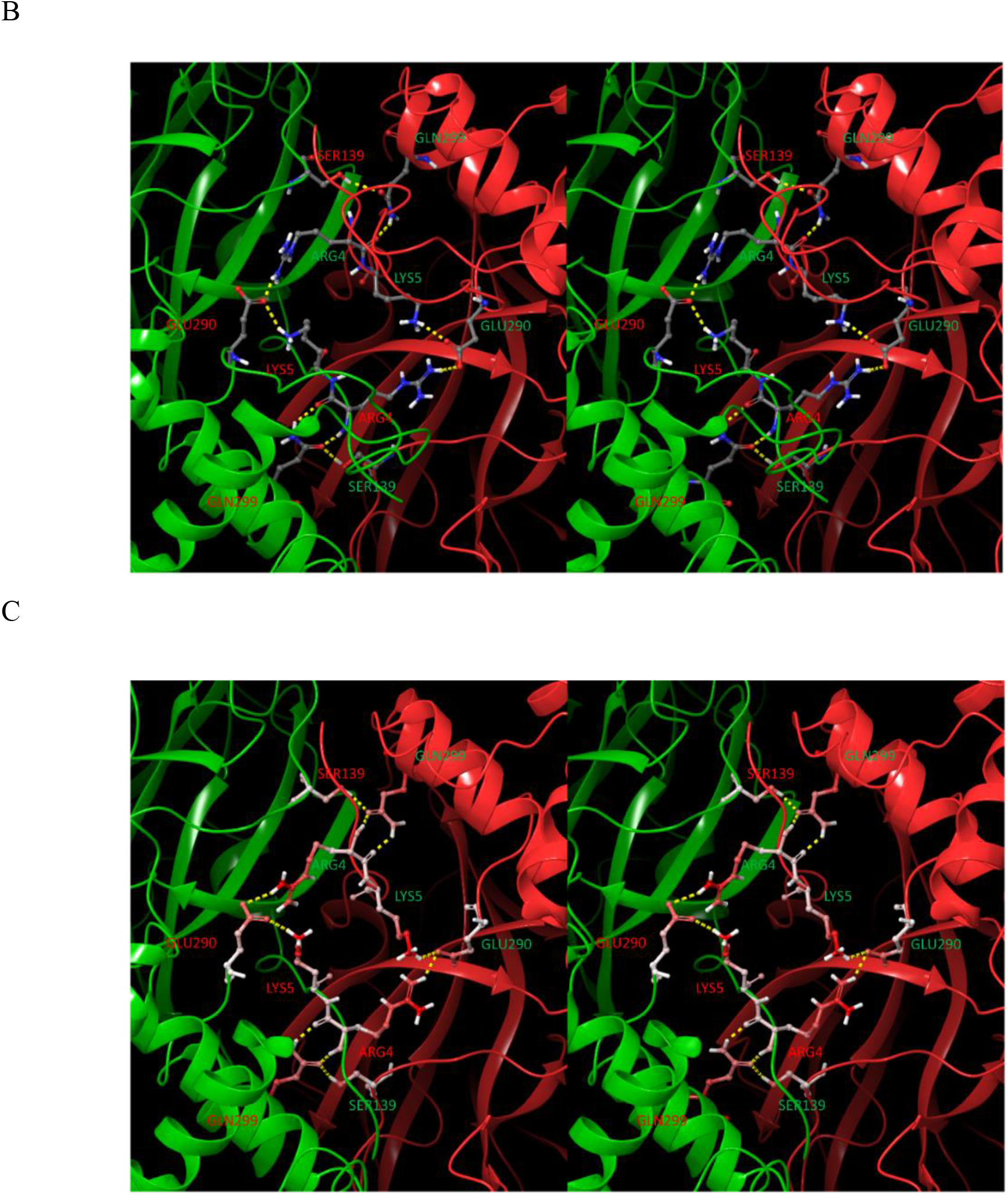
(A) Stereo view of the dimer interface of CoV M^pro^ in 2Q6G, with the individual subunits shown in red and green. Zoomed out view of the circuit-like H-bond network sandwiched between the NTLs of each subunit, and bridging across the networks of the individual subunits. (B) Same as A, except zoomed in to the inter-subunit region, showing the circuit-like H-bond network comprised of Arg4 and Lys5 of the NTL, together with intra-monomer Glu290 and Ser139. The native dimer interface is thus part of a global network of residues that play key roles in the conformational dynamics of the protein. (C) Same as B, except for CoV-2 M^pro^ in 6M03, noting the relatively high B-factors of the residues in this network, which are somewhat higher than those in 2Q6G.

**Figure 21.**
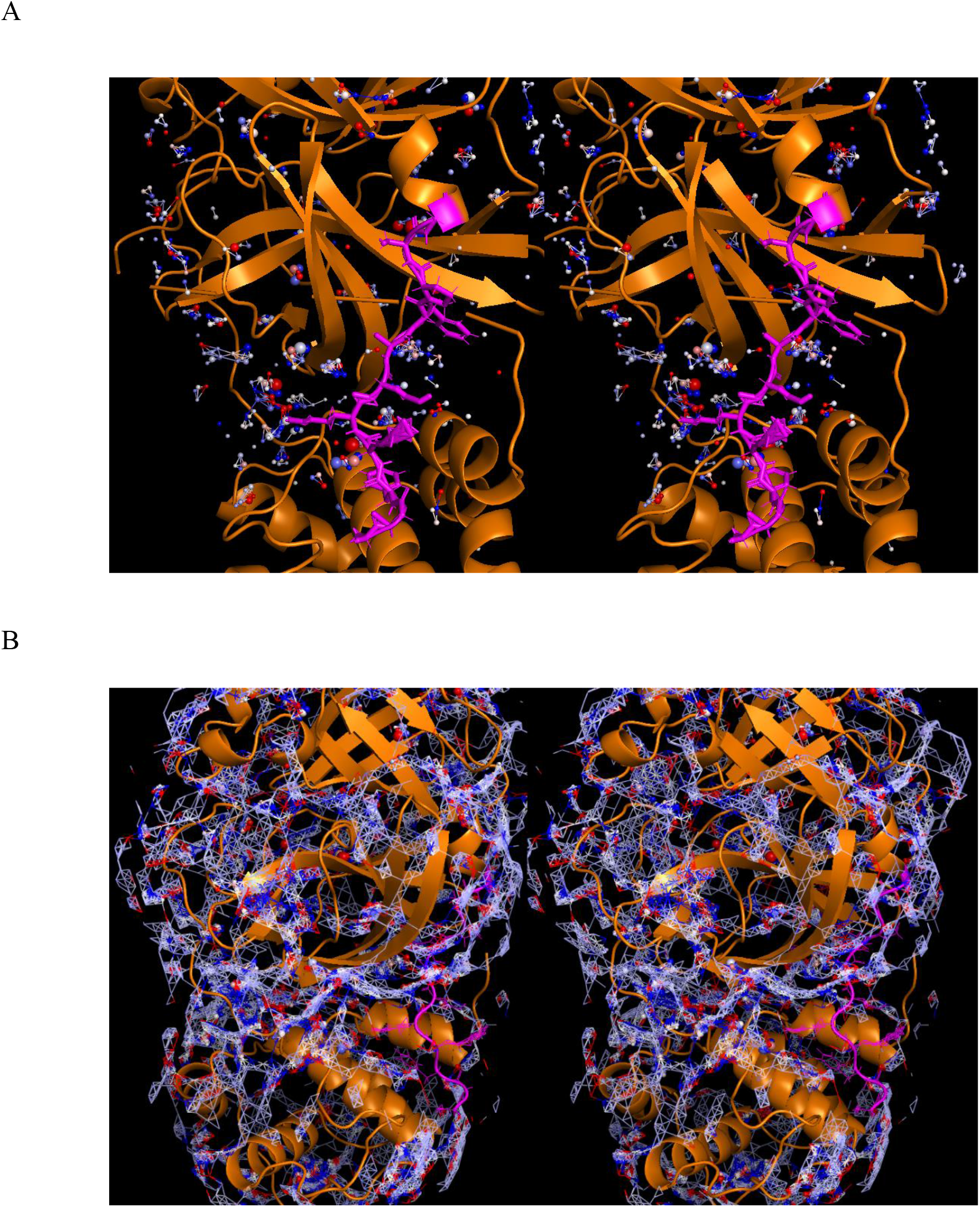
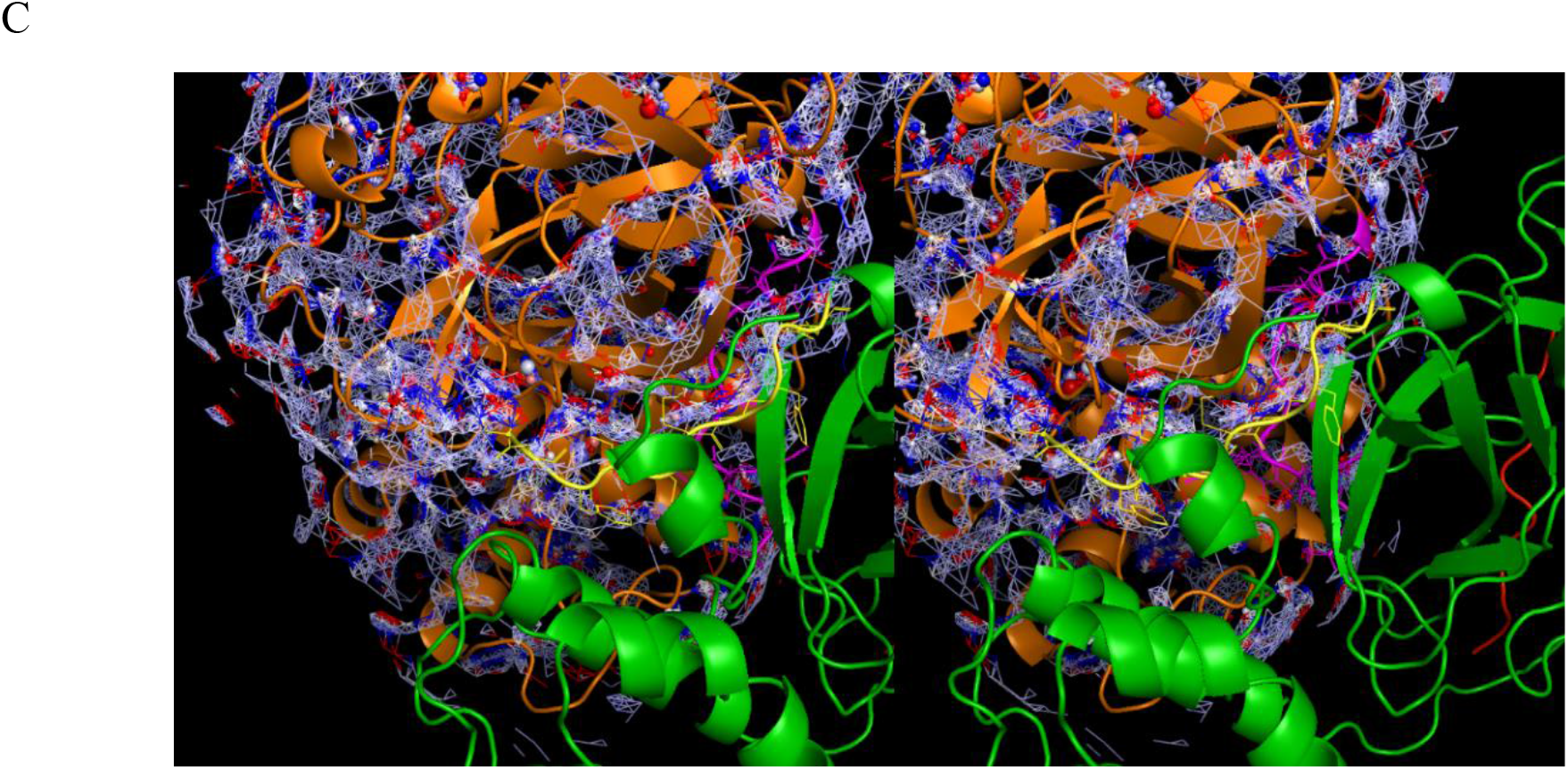
Stereo views of the WATMD-calculated solvation structure within the dimerization interface (same color-coding as in Figure 13). (A) The <u>pre-dimerization</u> interface of the time-averaged 2Q6G structure aligned to the α-carbons, showing only the reference subunit (including the NTL shown in magenta) used in the calculation, together with the H-bond enriched and depleted solvation hotspots. (B) Same as A, except showing the full set of non-bulk-like solvation (blue-white-red mesh surface) in the time-averaged charged voxel (TACV) surface. (C) Same as B, except showing the insertion subunit (green) (including the NTL shown in yellow) overlaid on the solvation structure of the reference subunit. High complementarity between the insertion NTL and a “filament” of the TACV is consistent with expulsion of mildly H-bond depleted water from the dimer interface (consistent with the expected transience of the dimeric form).

#### Solvation-powered substrate and inhibitor binding

We calculated the solvation structures of the <u>apo</u> 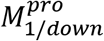 state in 2QCY (Figure 22A-C), the 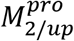 state in 6M03 (Figure 22D-F), and the boceprevir-bound structure in 6WNP (Figure 23A-B) in and around the AS using WATMD, in which the time-averaged protein structure was aligned to Met162-Glu166 (noting that WATMD results on bound structures from which the ligand has been removed are invalid in cases of induced fit). The occupancy of each individual AS pocket by bound substrates or inhibitors is governed by the degree of site-specific <u>mutual</u> complementarity between the protein and ligand solvation structures (the latter of which we did not calculate). As explained above, net H-bond free energy <u>gains</u> are derived primarily from the expulsion of H-bond <u>depleted</u> solvation from either or both of the binding partners (i.e. underlying the re-solvation cost), whereas expulsion of H-bond <u>enriched</u> solvation is typically unfavorable to varying degrees (underlying the de-solvation cost). Remarkably, H-bond enriched solvation is conspicuously absent from the AS, although many of the H-bond depleted positions are nevertheless polar (blue and red color-coding in the figures). We overlaid the native substrate (extracted from 2Q6G), boceprevir (extracted from 6WNP), and N3 (extracted from 6LU7) on the solvation structure of 2QCY (representative of the 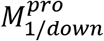 state to which substrates and inhibitors putatively associate), and checked for inhibitor-solvation structure complementarity (Figure 24). We propose that the R-groups of bound ligands wag in and out of their respective pockets at rates governed by pocket-dependent de-solvation and re-solvation costs (i.e. increased wagging at weakly H-bond depleted or bulk-like regions). Such motions may explain the high B-factors observed at the N-and C-termini of N3 in 6LU7 (Figure 25A), compared with the low B-factors in boceprevir in 6WNP (Figure 25B). High conformity between the observed ligand binding modes and the shapes/properties of the solvation structure is apparent in Figures 23–24 consistent with our claim that binding free energy is stored principally within binding site solvation (noting the same for other crystallized ligands that we examined). Nevertheless, room for further optimization of the inhibitors in our dataset is apparent (e.g. the P1 position of boceprevir (Figure 23B)).

**Figure 22.**
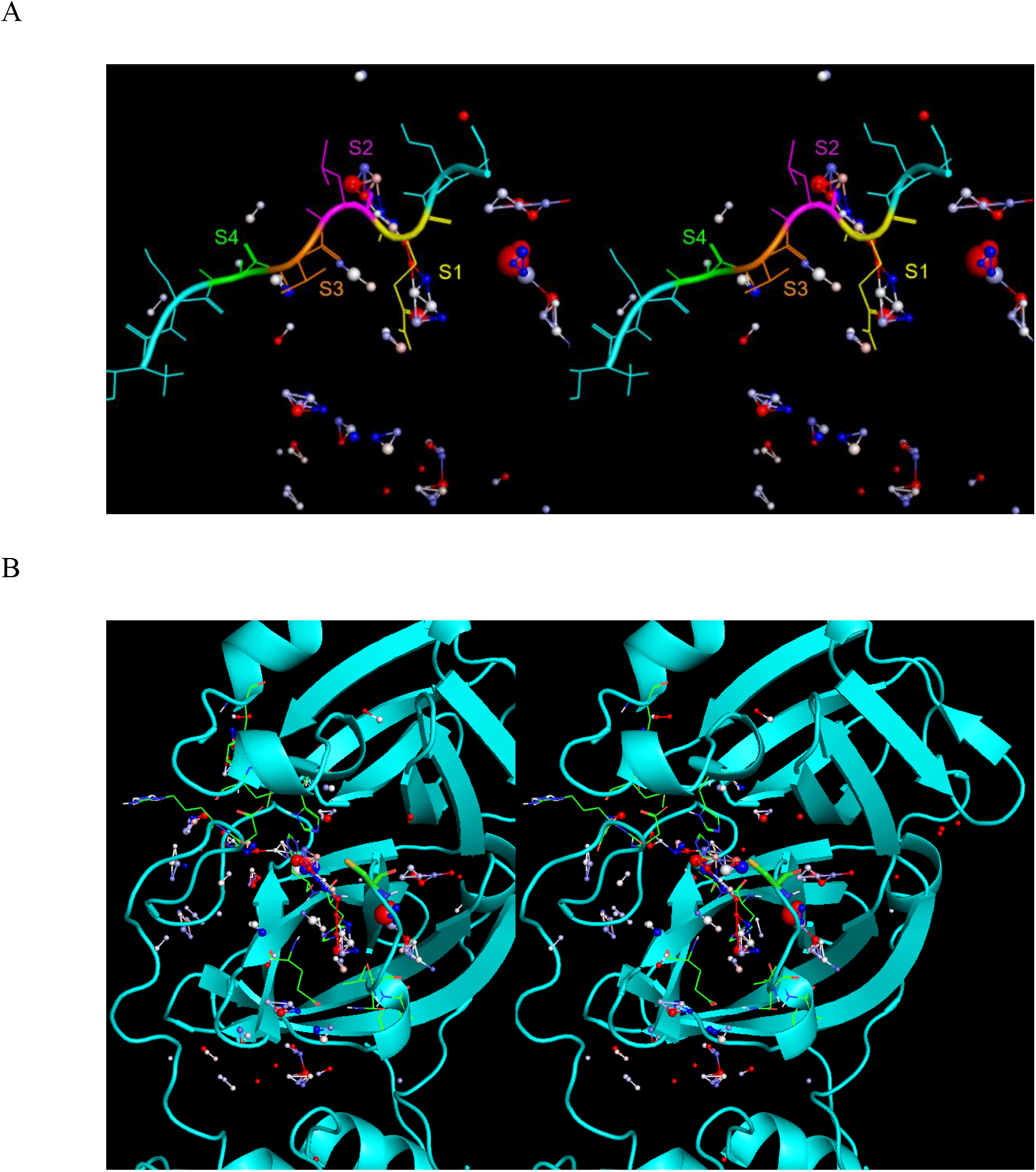
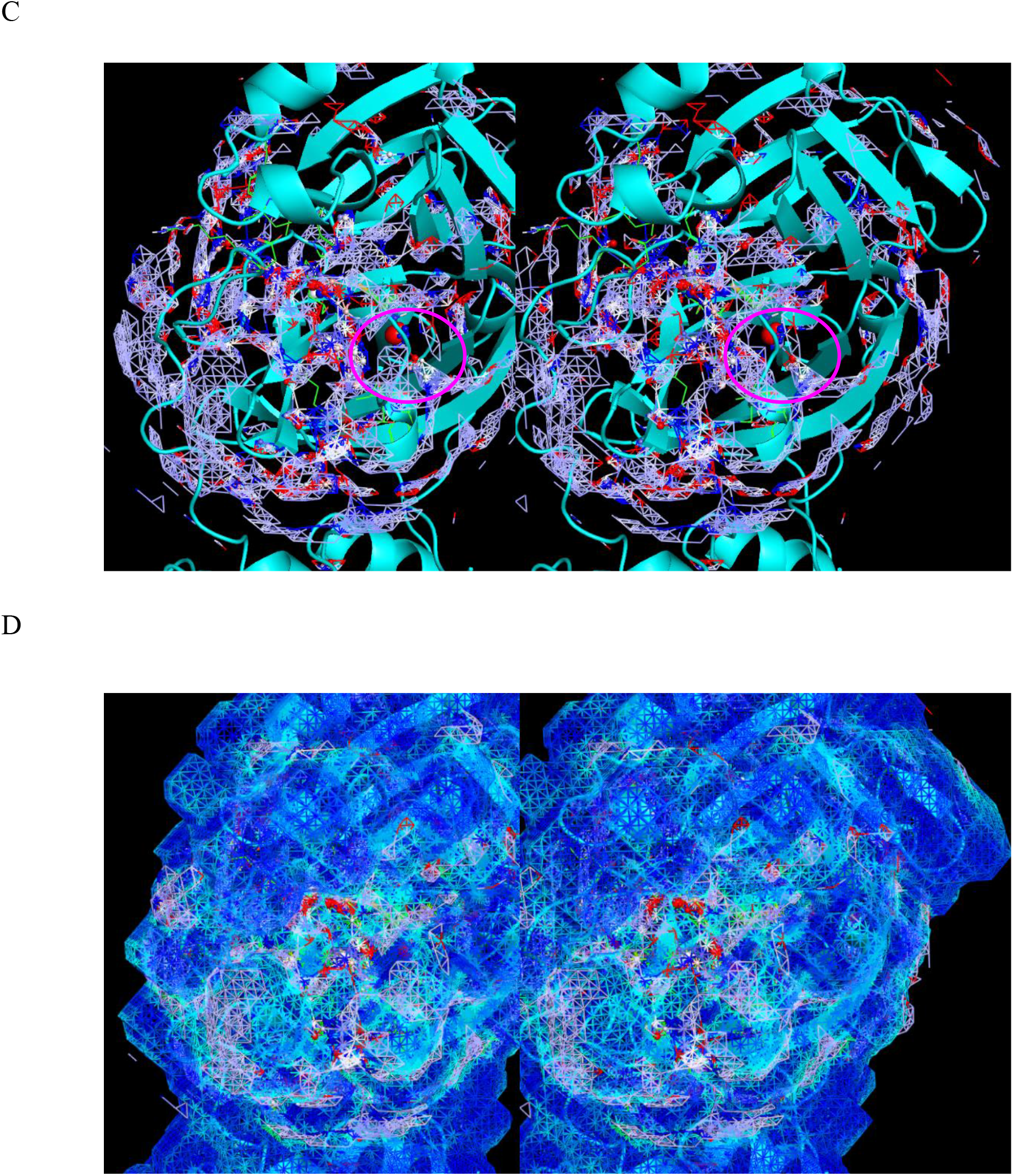
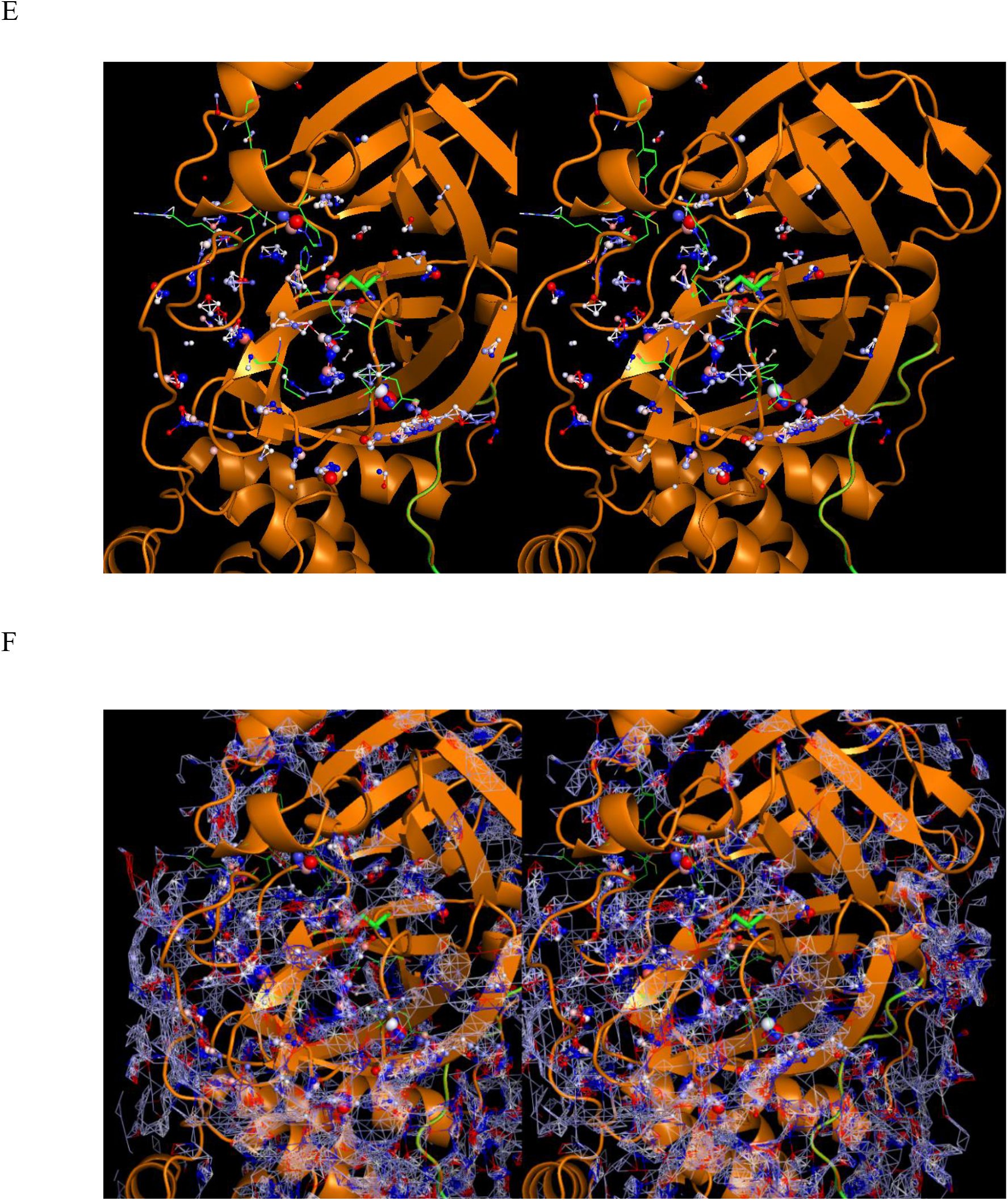
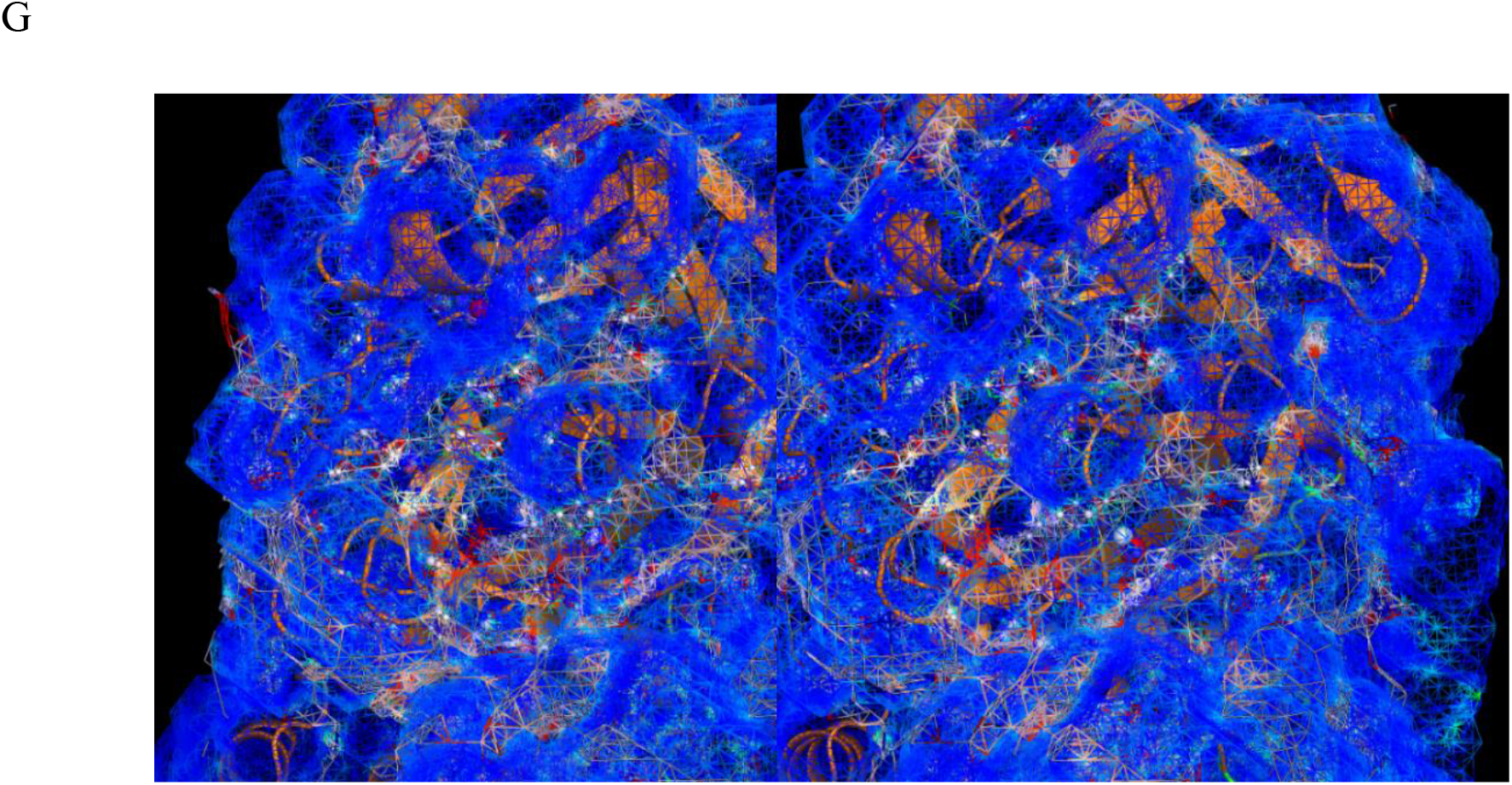
Stereo views of the solvation structures within the AS of <u>apo</u> CoV and CoV-2 M^pro^. (A) Mapping between the solvation hotspots in 2QCY and the P1-P4 substrate regions (shown for reference). (B) Apo 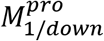 (2QCY). An H-bond depleted “water chain” spans between the S1 and S2 pockets, the expulsion of which is expected to slow substrate k_-1_ and inhibitor k_off_. (C) Same as A, except including the TACV. A high occupancy H-bond enriched hydration site is present within the peri-m-shaped loop region (circled in magenta), which is absent in the 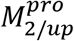 state (noting the presence of a high occupancy H-bond enriched solvation hotspot in the domain {1-2}-3 interface in the 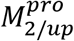 state (Figure 11E)). We postulate that the H-bond depleted solvation in the 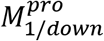 state is counter-balanced by this water. (D) Same as C, except including the time-averaged solvent-accessible surface (TASAS). (E) Apo 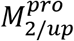 in 6M03. The S1 ⟶ S2 water chain is partially disrupted in this state, further suggesting that the AS in 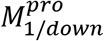 is optimized for substrate association. (F) Same as E, except including the TACV. (G) Same as E, except including the TASAS.

**Figure 23.**
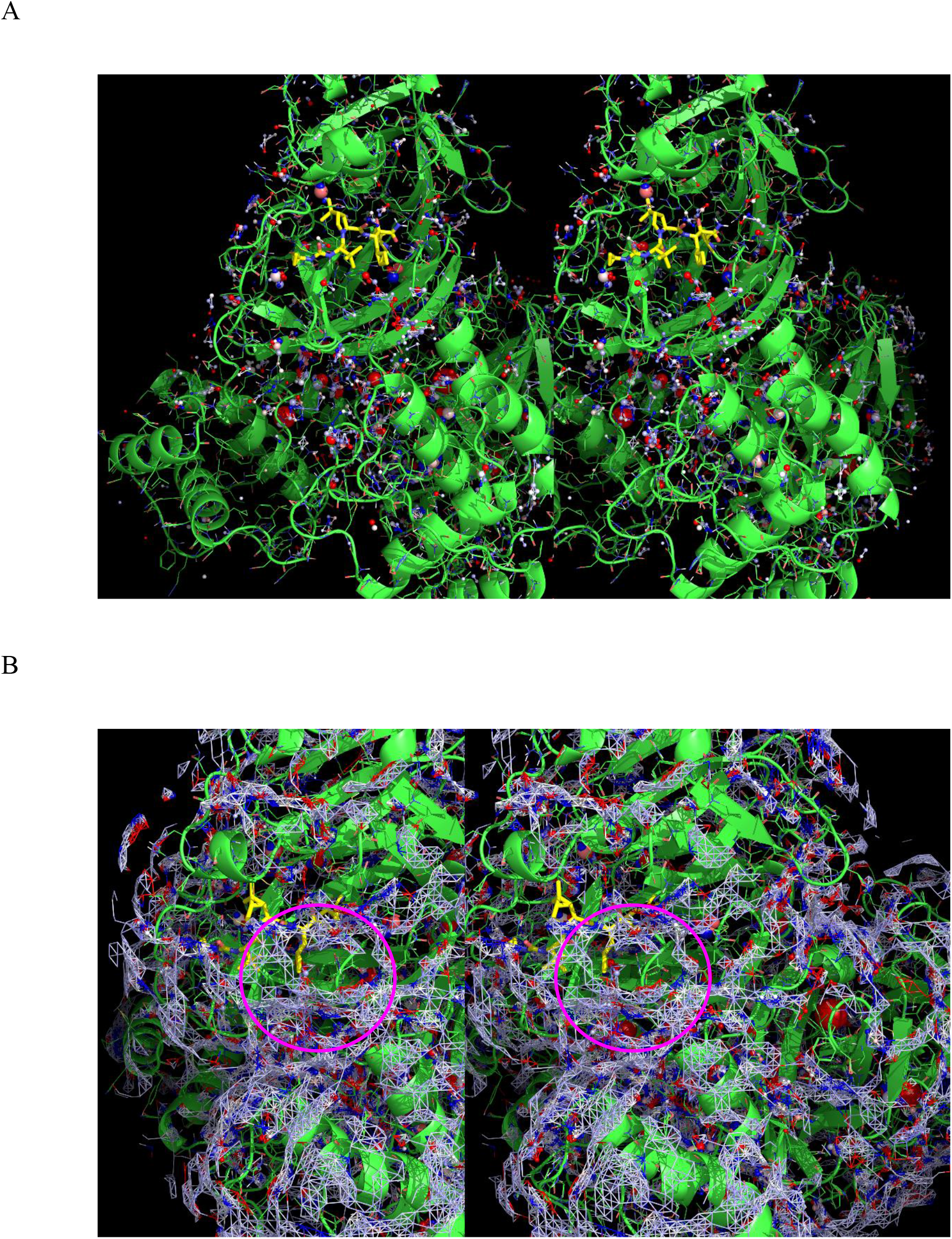
(A) Stereo view of the solvation structure of <u>boceprevir-bound</u> CoV-2 M^pro^ in 6WNP (with the inhibitor shown in yellow). (B) Same as A, except including the TACV. These results suggest that further expulsion of H-bond depleted water (circled in magenta) from the AS is conceivable via a macrocycle linking the P1 and P3 groups of boceprevir.

**Figure 24.**
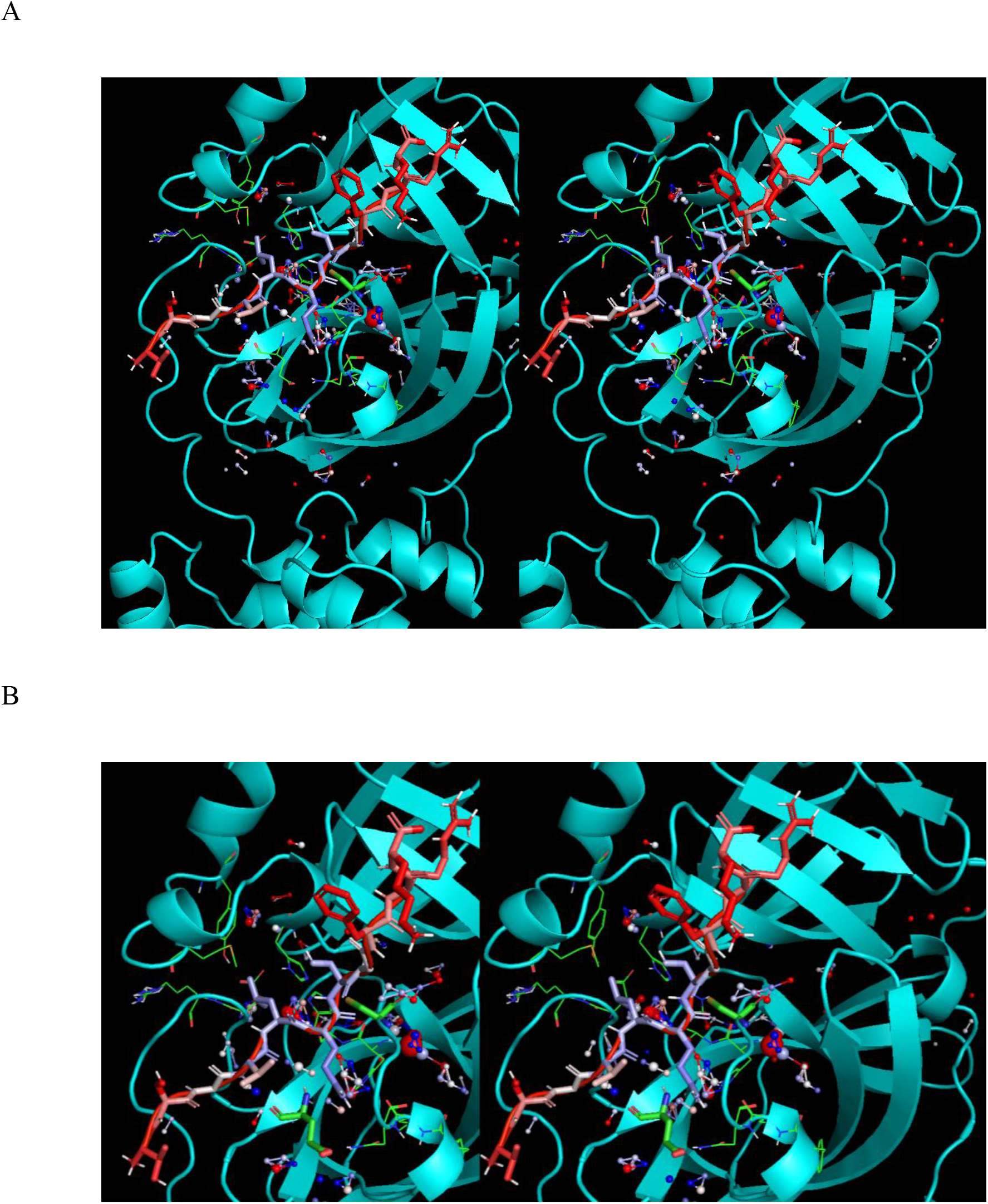
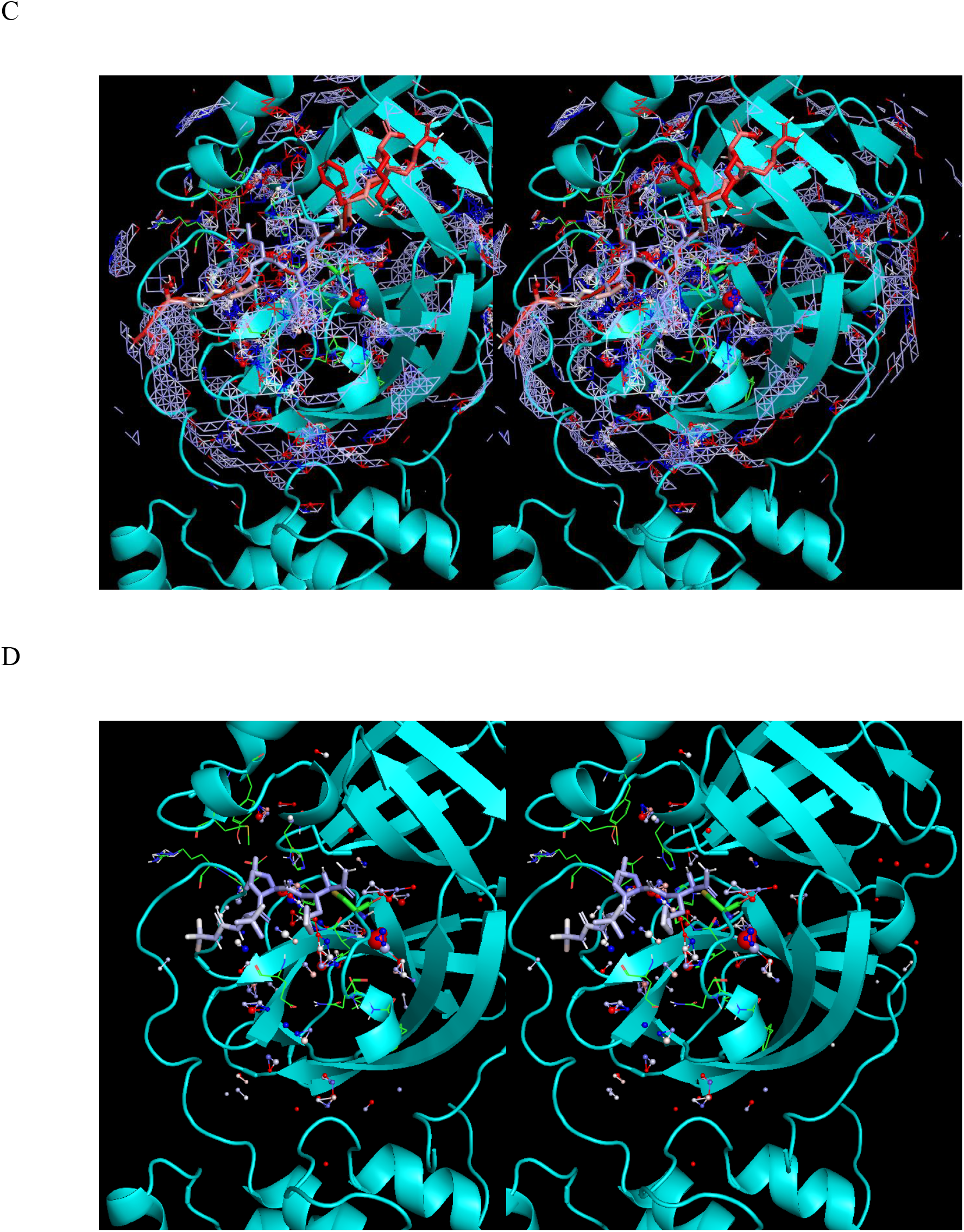
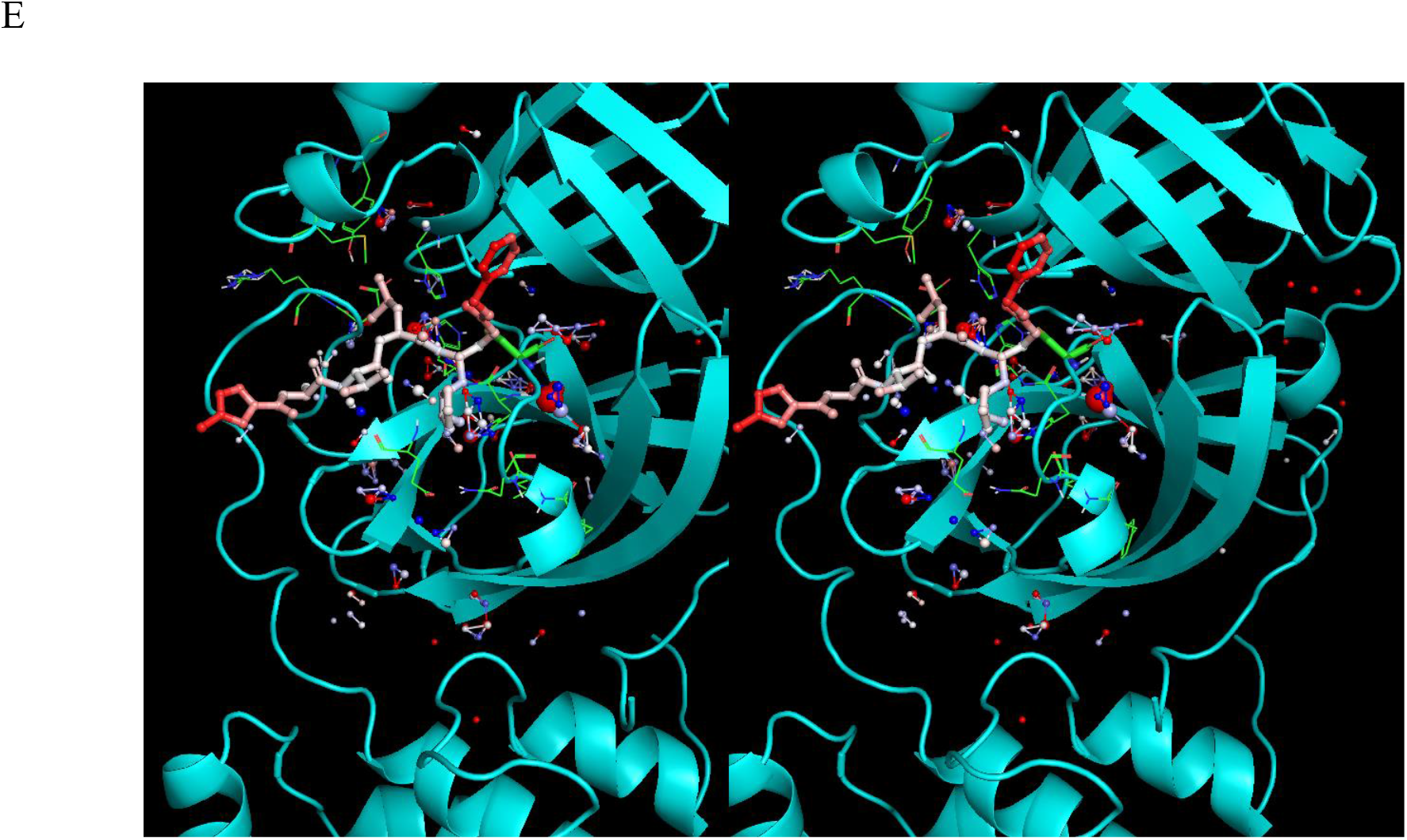
Same as Figure 22, except with crystallized reference ligands (colored by B-factors) overlaid on the solvation hotspots of 2QCY (ligands were extracted from the structures denoted by ‡ in Table 1). (A) The cognate substrate extracted from 2Q6G, noting high overlap between the ligand moieties and the solvation hotspots. (B) Same as A, except zoomed in around the S1 and Glu166 backbone regions. (C) Same as A, except including the TACV. (D) Boceprevir extracted from 6WNP. (E) N3 extracted from 6LU7, noting the high B-factors of the N- and C-terminal regions, which reside in bulk solvent.

**Figure 25.**
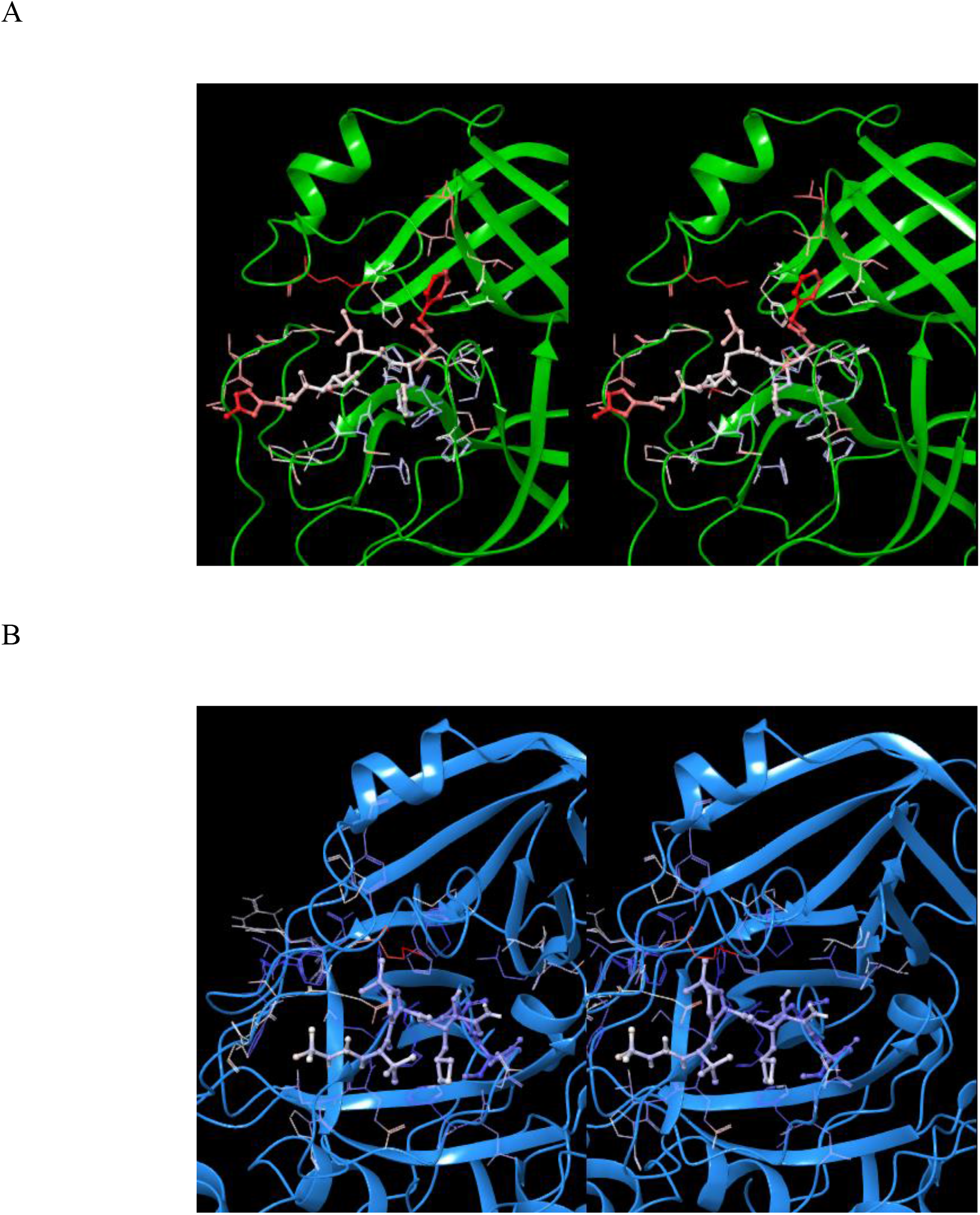
(A) Stereo view of the crystal structure of the dimeric inhibitor N3-CoV M^pro^ complex color-coded by B-factors (6LU7). (B) Same as A, except for boceprevir-CoV-2 M^pro^ in 6WNP, noting the cooler B-factors.

**Figure 26.**
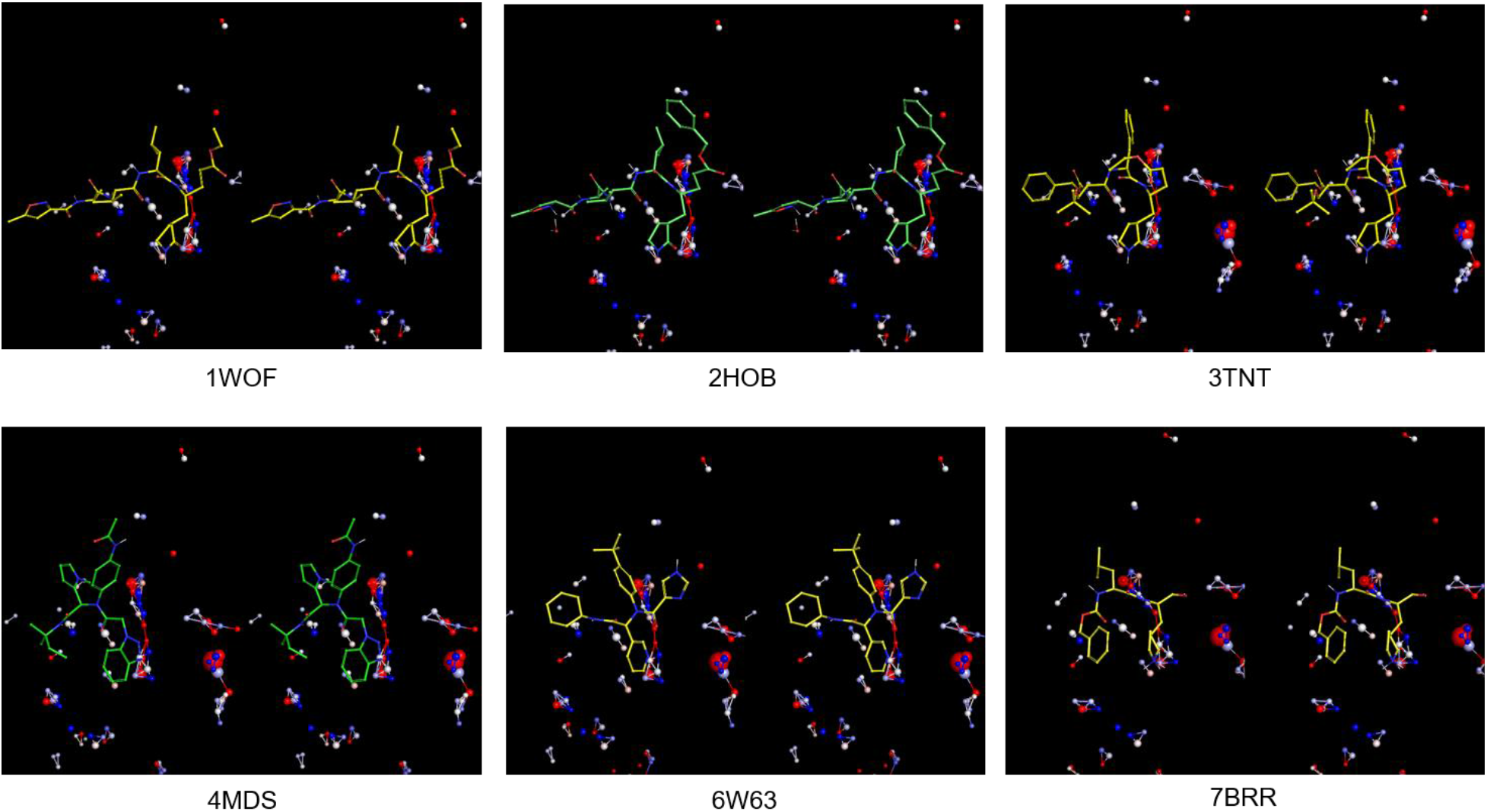
Stereo view of a set of representative crystallized inhibitors (listed in Table 2) overlaid on the time-averaged 2QCY structure generated by WATMD (all α-carbon atoms), showing inhibitor-solvation hotspot overlaps corresponding to expelled H-bond depleted water in the inhibitor-bound state. The results suggest that the k_off_ may be slowed by further optimization of the inhibitor-solvation structure relationship.

#### A non-equilibrium perspective on M^pro^ catalysis and inhibition

Enzyme kinetics are typically measured and analyzed under the assumption that the rate of enzyme-substrate complex formation and turnover are equivalent (the steady state assumption). However, this assumption need not apply under native cellular conditions, in which the enzyme and substrate concentrations vary over time, and the rate of enzyme-substrate complex formation is necessarily described using ordinary differential equations (ODEs) of the form:

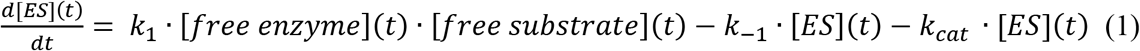

where ES denotes the enzyme-substrate complex, and k_1_, k_-1_, and k_cat_ denote the association, dissociation, and turnover rates, respectively. At constant free enzyme and substrate concentrations, equation 3 reduces to 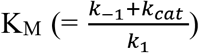 and the Michaelis-Menten equation. The rate of M^pro^ catalysis depends on several contributions governing the enzyme and substrate concentrations (polyprotein expression, possible M^pro^ degradation, 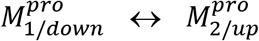 transitioning, substrate binding, and dimerization), which is described by the following set of coupled ODEs corresponding to the reaction scheme summarized in Figure 26:

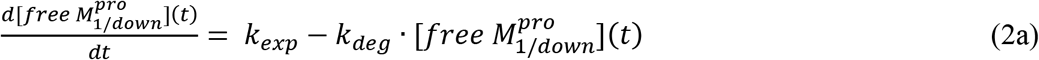

where k_exp_ and k_deg_ are the rates of monomer synthesis and monomer degradation, respectively (assuming the possible existence of one or more protein degradation pathways).

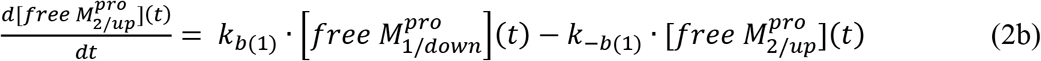

where k_b(1)_ and k_-b(1)_ are the rates of rocking between the two domain 3 positions in the free M^pro^ monomer.

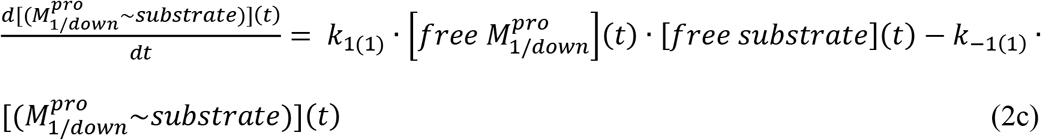

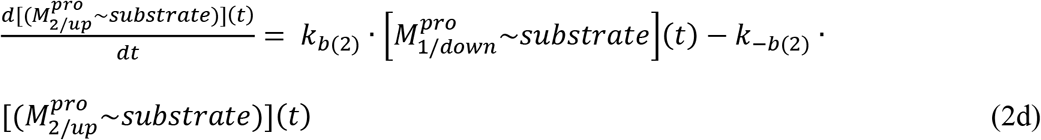

where k_1(1)_ and k_-1(1)_ are the rates of M^pro^-substrate association and dissociation, respectively, and k_b(2)_ and k_b(−2)_ are the rocking rates between the two domain 3 positions in the substrate-bound M^pro^ monomer.

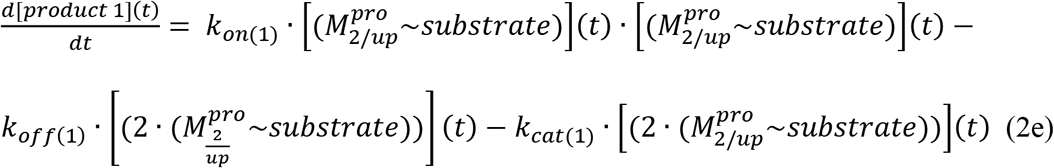

where product 1 is the hydrolyzed C-terminal product, k_on(1)_, k_off(1)_, and k_cat(1)_ are the rates of dimerization, dimer-substrate dissociation, and turnover, respectively

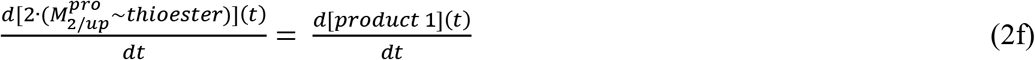

where 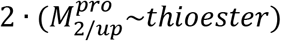 is the thioester adduct, which is equal to the rate of product 1 generation.

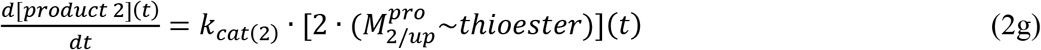

where product 2 is the hydrolyzed C-terminal product, and k_cat(2)_ is the turnover rate constant for thioester adduct decay (where the functional unit is dimeric).

**Figure 26.**
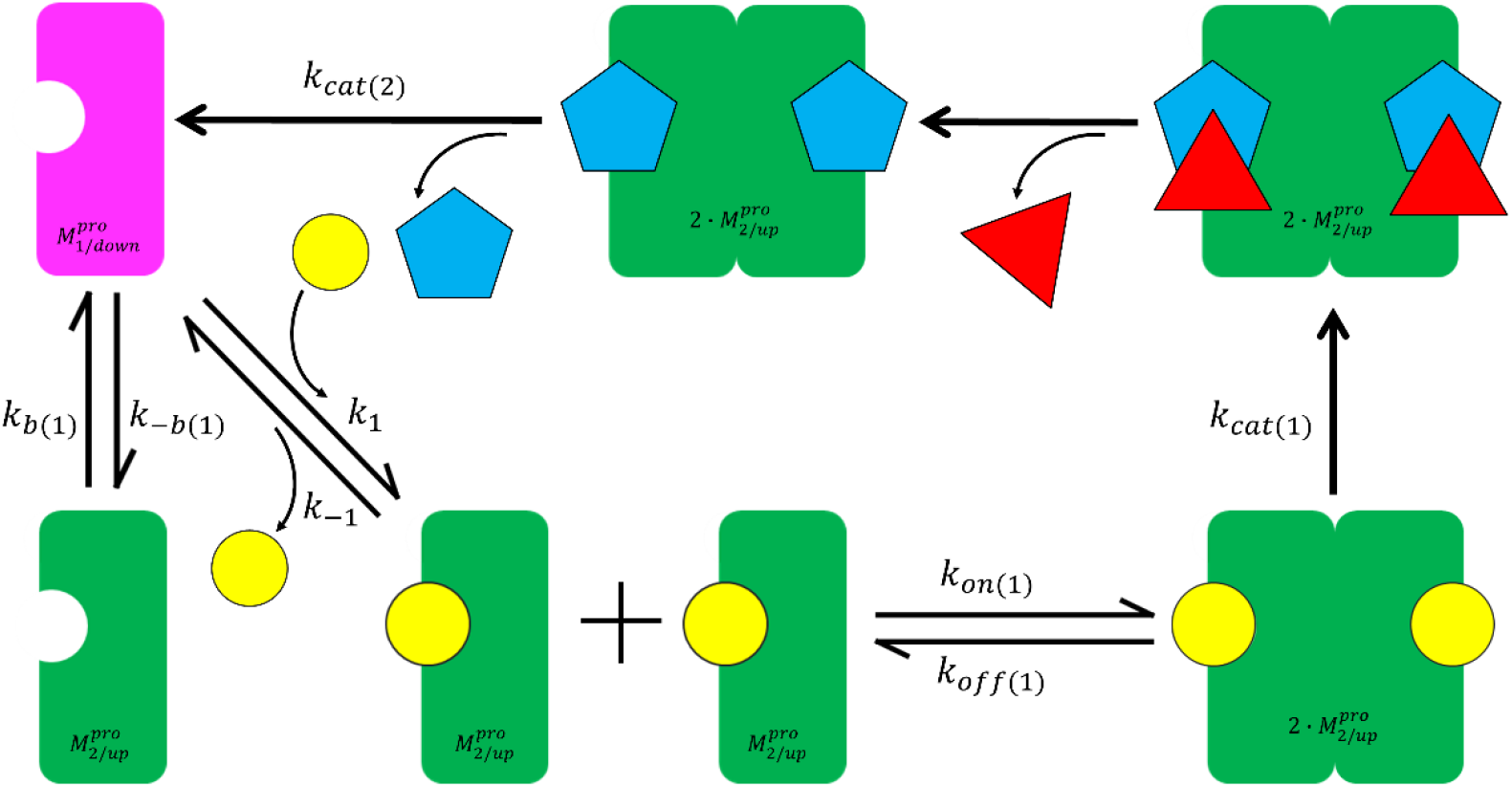
The proposed M^pro^ reaction scheme, including substrate binding, domain 3/m-shaped loop rearrangement, dimerization, turnover, and leaving group dissociation (the rate constants are defined in the text).

Under non-equilibrium conditions, the catalytic efficiency of M^pro^ depends on synchronous dimerization, substrate binding, and turnover, equating to:

1. A substrate-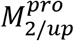 association rate approaching the turnover rate (k_1(1)_ ≳ k_cat_). The slowest binding step is otherwise rate-determining.
2. A dimer 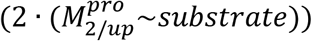 lifetime on the order of the reaction time constant (1/k_off(1)_ < 1/k_cat_). Turnover is disrupted when the dimer and/or bound substrate dissociate prior to product formation (noting that K_d_ is agnostic to binding partner exchanges, whereas enzyme-mediated turnover is not [5]).

For non-covalent inhibitors:

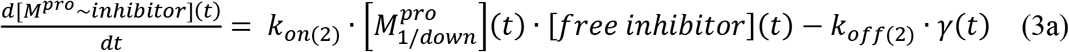

where k_on(2)_ and k_off(2)_ are the inhibitor association and dissociation constants, respectively. We assume that inhibitors bind to the 3_10_ helical state of the m-shaped loop.

For <u>reversible</u> covalent thioester inhibitors:

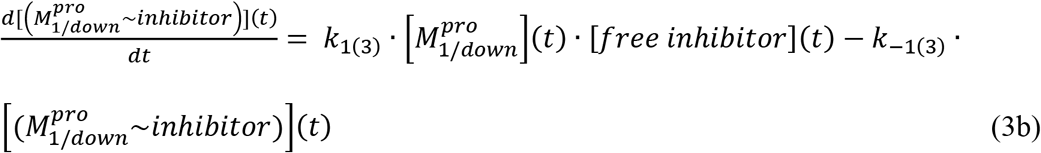

where k_1(3)_ and k_-1(3)_ are the unreacted inhibitor-M^pro^ association and dissociation constants, respectively.

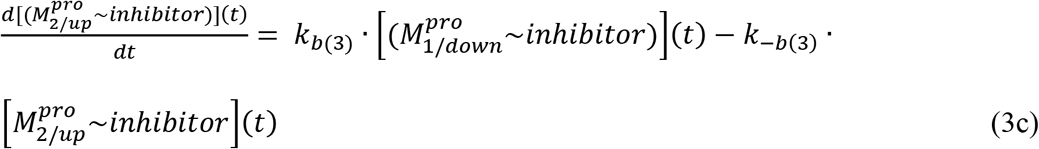

where k_b(3)_ and k_-b(3)_ are the rates of rocking between the two domain 3 positions in the inhibitor-bound M^pro^ monomer.

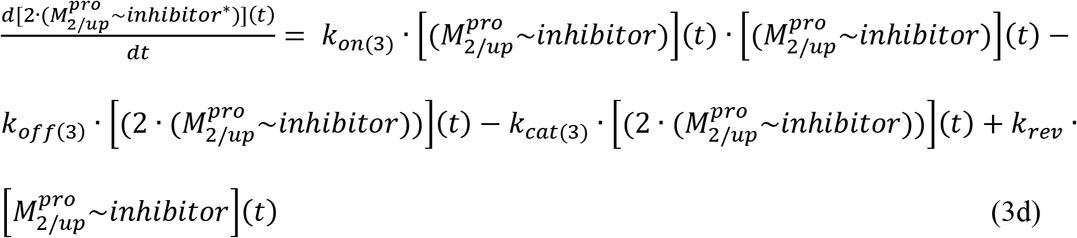

where k_on(3)_, k_off(3)_, and k_cat(3)_ are the rates of 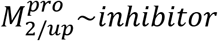 association, dissociation, and adduct formation (denoted as 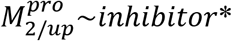, respectively, and k_rev_ is the rate of adduct hydrolysis (noting that dimer dissociation is expected upon adduct hydrolysis).

For <u>irreversible</u> covalent thioester inhibitors:

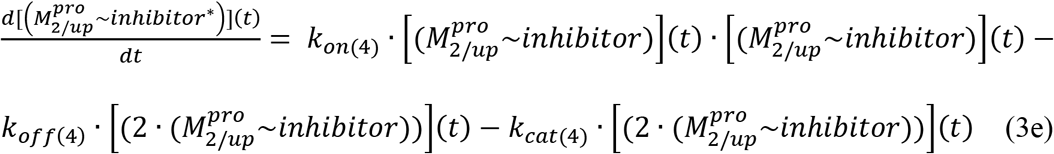

where k_1(4)_, k_-1(4)_, and k_cat(4)_ are the rates of inhibitor-M^pro^ association, dissociation, and adduct formation, respectively, and k_on(4)_ and k_off(4)_ are the dimerization and dimer dissociation rates, respectively (noting that slow dimer dissociation may result in the presence of irreversible adduct formation).

The solution to the above set of coupled ODEs consists of an exponential function, commensurate with rapid growth in polyprotein processing and virion production over time. However, implementation of this model leads to a catch-22, in which experimental parameter measurement depends on the model and vice versa. The enzyme kinetics data reported for SARS-CoV-2 M^pro^ is out of line with respect to that of other characterized enzymes [38], as follows:

1. K_M_ ranging between 189.5 and 228.4 μM for three model substrates [39] (consistent with other reported values [34,40,41]), compared with the median K_M_ of 130 μM reported for 5,194 enzymes.
2. k_cat_ ranging between 0.05 and 0.178 s^−1^, compared with the median k_cat_ of 13.7 s^−1^ reported for 1,942 enzymes. Slow turnover by CoV 3CL^pro^ has been attributed to slow hydrolysis of the acyl adduct (reaction step 2), rather than slow proton abstraction or TI formation (reaction step 1) [20].
3. k_cat_/K_M_ ranging between 219 and 859 M^−1^ s^−1^, compared with the median k_cat_/K_M_ of 125E3 reported for 1882 enzymes. The k_cat_/K_M_ equates to unrealistically slow processing throughput (e.g. ~1 mM of substrate is needed to achieve an overall processing rate of 1 s^− 1^, compared with 8 μM at the median k_cat_/K_M_).

The above discrepancies may result from lack of consideration of substrate and dimerization contributions to M^pro^ activation, in which case, data analysis cannot be based simply on the total enzyme and fixed substrate concentrations. Furthermore, the dimerization K_d_ necessarily differs between the substrate-bound and unbound states, as confirmed by Cheng et al. for CoV M^pro^, who reported substrate-bound and unbound K_d_ values of 0.8 μM and 2 μM, respectively [36]. Graziano et al. reported a somewhat higher dimerization K_d_ for the unbound form based on three orthogonal measurement techniques (ranging between ~5 and 7 μM) [35]. In practice, dimer buildup is a non-equilibrium process due to the time-dependence of the total M^pro^ and polyprotein concentrations resulting from first order auto-cleavage, and furthermore, the monomer-dimer-substrate distribution is highly non-linear due to the 3-way relationship among the species. We calculated the equilibrium dimer concentration as a function of substrate-independent free monomer concentration in multiples of K_d_ = 5 and 0.8 μM (Table 3). It is apparent from the table that the fractional dimer concentration increases slowly as a function of total M^pro^ concentration (i.e. dimer + monomer). A significant fraction of monomer is present at the optimal <u>total</u> M^pro^ concentration, which is tipped toward the dimer in the presence of substrates.

**Table 3.** The equilibrium dimer fraction and concentration as a function of substrate-independent and substrate-dependent monomer concentrations in multiples of K_d_ (based on the Hill equation).

| $K_d$<br>($\mu\text{M}$) | [Monomer]<br>($\mu\text{M}$) | Dimer<br>fraction | [Dimer]<br>($\mu\text{M}$) | Monomer<br>fraction | [Monomer]<br>($\mu\text{M}$) |
| --- | --- | --- | --- | --- | --- |
| Substrate-unbound |  |  |  |  |  |
| 5.0 | $1 \cdot K_d = 5.0$ | 0.5 | 2.5 | 0.5 | 2.5 |
| 5.0 | $2 \cdot K_d = 10.0$ | 0.67 | 6.7 | 0.33 | 3.3 |
| 5.0 | $3 \cdot K_d = 15.0$ | 0.75 | 11.25 | 0.25 | 3.75 |
| 5.0 | $10 \cdot K_d = 50.0$ | 0.91 | 45.5 | 0.09 | 4.5 |
| Substrate-bound |  |  |  |  |  |
| 0.8 | $1 \cdot K_d = 0.8$ | 0.5 | 0.4 | 0.5 | 0.4 |
| 0.8 | $2 \cdot K_d = 1.6$ | 0.67 | 1.07 | 0.33 | 0.53 |
| 0.8 | $3 \cdot K_d = 2.4$ | 0.75 | 1.8 | 0.25 | 0.6 |
| 0.8 | $10 \cdot K_d = 8.0$ | 0.91 | 7.28 | 0.09 | 0.72 |

The M^pro^ dynamics proposed in this work is remarkably similar to that proposed by Datta et al. for caspase-1 [42], noting that the substrate-bound caspase-1 activated dimer/monomer ratio = 20-fold (corresponding to a 10-fold increase in k_cat_/K_M_) versus ~2.5 to 9-fold for M^pro^, depending on the dimerization K_d_ of the unbound subunits (the corresponding increase in k_cat_/K_M_ is unknown). However, whereas suboptimal inhibition was found to promote caspase-1 dimerization and activation, this effect is unlikely with M^pro^ due to the decreased accessibility of the S1 pocket in the dimeric 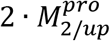 form of the protein. The experimental protocol developed for substrate-induced dimerization/activation of caspase-1 (described in the Supplementary Information of [42]) suggests that a similar protocol may be necessary for the correct measurement of M^pro^ enzyme kinetics, which may explain the unrealistic M^pro^ k_cat_/K_M_ determined from the conventional Michaelis-Menten enzyme kinetics model.

#### Implications of our findings for inhibitor design

The time-dependence of all processes in which M^pro^ participates under native *in vivo* conditions, including monomer expression and degradation, rearrangement, and substrate binding are key elements of inhibitor design (except in the case of irreversible covalent inhibitors, the occupancy of which accumulates as a function of concentration). In our previous work, we demonstrated the high sensitivity of non-covalent drug occupancy to the rates of binding site buildup and decay (in order of precedence: k_on_, concentration(t), and k_off_) [1]. Two non-mutually exclusive M^pro^ inhibition approaches are conceivable:

1. Inhibition of M^pro^ auto-cleavage in cis. Under this approach, the inhibitor k_on_ must necessarily keep pace with the rate of polyprotein synthesis, and remain bound throughout the protein lifetime. However, this approach is likely nonviable if the cleavage substrate folds within the AS.
2. Inhibition of M^pro^-mediated polyprotein cleavage in trans. We assume that most covalent inhibitors containing substrate-like P1 groups bind to the monomeric 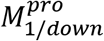 (S1 pocket-accessible) form of post-cleaved M^pro^.

It is reasonable to assume that efficacious inhibition <u>at the lowest possible concentration</u> depends on high occupancy of the AS over time (especially in the absence of M^pro^ degradation or down-regulation), which in turn, depends on kinetically tuned inhibitor binding (such that k_on_ ≈ k_i_ and k_off_ approaches the protein lifetime), as follows:

1. High complementarity between the inhibitor solvation structure and that of the <u>monomeric</u> M^pro^ AS (including the Glu166 backbone β-hairpin positions and S1 pocket), resulting in maximal expulsion/re-solvation cost of H-bond depleted solvation (noting that tolerance for H-bond enriched <u>inhibitor</u> solvation may be limited largely to the P1 position).
2. Reversible covalent reactivity needed to further slow k_off_ beyond the re-solvation free energy contribution, the latter of which may result in an inadequate t_1/2_ of the bound state relative to the pharmacokinetic timescale (given the possibility that the M^pro^ concentration does not decay over time in the absence of a degradation pathway). Rapid adduct formation is conceivable based on the general cysteine protease mechanism reported by other workers, where the rate-determining step consists of hydrolysis (step 2) rather than thioester formation (step 1) [22].
3. Irreversible covalent inhibition is driven largely by covalent adduct <u>accumulation</u>, although efficacy may nevertheless depend on the accumulation <u>rate</u> (noting that leakage of M^pro^ and its downstream products may result from slow accumulation due to slow k_on_ and/or k_cat_).
4. Inhibitor-mediated stabilization of the 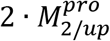 state needed for enzyme-mediated covalent bond formation (just as for endogenous substrates).
5. Fast k_cat_ governing covalent adduct formation.

Surprisingly, our WATMD results suggest the AS is solvated exclusively by bulk-like and mildly to strongly H-bond depleted solvation, although the protein environments around the solvation sites contain H-bond acceptor and donor groups on a case-by-case basis (the water H-bonds are simply fewer or weaker than in bulk solvent). As such, the de-solvation cost (i.e. the free energy lost in expelling solvating water to bulk solvent) underlying the association barrier is relegated largely to the ligand, whereas the re-solvation cost underlying the dissociation barrier (i.e. the free energy lost in transferring bulk solvent to H-bond depleted solvation sites on the dissociated partners) is relegated to the AS (and possibly the ligand as well). Although, the numerical magnitudes of these costs are beyond the scope of our calculations, it is reasonable to assume that, for substrates, they reside within a Goldilocks zone giving rise to k_1_ ≈ k_i_ and k_-1_ ≲ k_cat_ (noting that product inhibition would occur in the presence of excessive H-bond depleted solvation within the AS).

Remarkable complementarity between inhibitor and substrate H-bond groups is apparent from overlays of bound M^pro^ crystal structures on our predicted solvation structures (Figures 24 and 26), consistent with the key de-solvation contribution to the binding free energy. Mutual mismatches between the bound inhibitors and solvation structures, together with intra-ligand de-solvation costs and entropic losses, may be reflected qualitatively in higher B-factors (Figure 25A). Occupancy detractors include:

1. The high entropic cost of binding flexible peptide-like substrates (reflecting the cost of ordering) also factor into the association free energy barrier.
2. The lack of an optimal P1 group, which is expected to slow k_on_ (and possibly k_cat_) and speed k_off_ due to higher inhibitor desolvation cost and indirect loss of substrate-induced enzyme activation (corresponding to the lack of alignment of the oxyanion hole in the 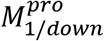 state). Furthermore, the lack of ligand-protein solvation complementarity in the S2, S3, and S4 pockets can result in independent binding/re-binding behavior (“wagging”) of the occupying P2, P3, and P4 groups (reflected in the high B-factors in 6LU7) due to local solvation free energy losses in each pocket. Such losses can result in low fractional occupancy of the closed P1 pocket across the M^pro^ population.

From a systems perspective, efficacious inhibition/occupancy depends on reducing the active M^pro^ population below a critical threshold (noting that all uninhibited copies can drive the downstream processes), <u>and maintaining this inhibition level over time</u> (keeping in mind that M^pro^ inhibition during the virion production phase may have little impact on the disease outcome given that the ship has already sailed by then).

## Discussion

The primary objective of preclinical drug discovery consists of <u>predicting</u> chemical entities that can cure or alleviate human disease via a combination of experimental and *in silico* approaches. In the absence of a theoretical foundation to formulate such predictions, drug discovery is relegated to a step-wise trial-and-error process centered on empirical approaches and data fitting techniques (i.e. inductive reasoning-based). Whereas drug discovery is predicated on equilibrium drug-target/off-target structure-free energy relationships (expressed as n · K_d_ or n · IC_50_, where n is a scaling factor between the drug concentration at 50% occupancy versus that at the efficacious occupancy), cellular function and pharmacodynamics are predicated on non-linear dynamic structure-kinetics relationships, in which targets/off-targets, their endogenous cognate partner(s) and drug concentrations vary over time. The equilibrium and non-equilibrium worlds rarely converge due in no small measure to the fact that free energy, occupancy, and concentration/exposure <u>under</u> *in vivo* <u>conditions</u> are not related by empirical principles (e.g. ΔG = −RT · ln(K_d_), Hill equation, Michaelis-Menten equation), but instead by <u>coupled</u> <u>time-dependent</u> systemic relationships, such as those outlined for M^pro^ in the previous section. Recognizing that <u>cellular</u> function/dysfunction is generated from coupled sets of dynamic solvation free energy-powered <u>molecular</u> rearrangements (that we refer to as “physical math”) is the first step toward transforming drug discovery (i.e. a predominantly data modeling-based undertaking) into drug design (a first principles theory-based undertaking). The first step is to improve the predictiveness of *in vitro* data via *in vivo*-relevant non-equilibrium dynamic measurements and models of drug-target/off-target binding. Cellular function is inextricably linked to dynamic biomolecular concentrations and dynamic state occupancy, which are the primary currency by which information is conveyed and processed. It is evident that each molecular state is optimized by nature according to the following criteria:

1. The timescale of the processes in which the species participate (e.g. 1/_kcat_), such that no entry or exit step to/from a given state becomes excessively rate-determining. As such, H-bond depleted solvation is expected to be sparingly present in native binding sites (although H-bond depleted solvation plays a key role in both inter- and intra-molecular rearrangements, as well as protein folding).
2. Tuning between the transition rate to/from each molecular state and the rates of buildup and decay of the molecular species or binding site (if the train is missed, it matters not how long the trip). We showed in our previous work that optimal dynamic drug-target occupancy depends first and foremost on the drug-target association rate constant (k_on_, k_1_), and that the k_on_ of many marketed drugs is fast, even when the k_off_ is slow [1].

In this work, we have attempted to identify and characterize the dynamic states of M^pro^, and relate the rates of buildup and decay of those states to structure-solvation free energy relationships (i.e. de-solvation and re-solvation costs/barriers) needed to inform drug design. The general idea is to reverse-engineer sets of high potential energy solvation (relative to bulk solvent) into complementary ligand solvation structures capable of achieving efficacious non-equilibrium occupancy at their intended targets via:

1. Expulsion of H-bond depleted solvation from the binding interface.
2. Mutual H-bond replacement of H-bond enriched “gatekeeper” solvation (which is absent in the AS of M^pro^) that is necessary for access to H-bond depleted solvation.

The critical question for M^pro^-based anti-viral therapies concerns the time-dependent <u>uninhibited</u> fraction of the enzyme that is incapable of sustaining viral replication, which constitutes the minimum dynamic occupancy level that efficacious inhibitors must achieve and maintain. As for cognate substrates, covalent inhibitors must associate to, and stabilize the active dimeric form of the enzyme (noting that occupancy by irreversible covalent inhibitors accumulates over time). Accurate qualitative determination of M^pro^ enzyme kinetics and time-dependent concentration buildup/decay of the monomeric and dimeric species (e.g. possibly measureable via replicon assays) in the context of the true substrate binding/catalytic mechanism is essential for successful drug design targeting k_on_, k_off_, and pharmacokinetic exposure. Furthermore, enzyme kinetics models must account for the dependencies between product generation and dynamic substrate, monomer, and dimer concentrations, and substrate-induced dimerization (possibly adapted from the putatively analogous caspase-1 mechanism [42]). Furthermore, inhibitor binding dependencies that are insensitive to resistance mutations are essential. We have offered a starting point for the development of such models, stopping short of the actual design of M^pro^ inhibitors, which is beyond the scope of this work.

## Conclusion

We showed that the dynamic non-covalent intra- and inter-molecular rearrangements underlying M^pro^ structure-function, consisting of intra-molecular 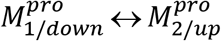 state transitions, substrate binding, and dimerization, are powered by inter-dependent multi-correlated solvation free energy barriers that subserve <u>transient and specific</u> structural responses (a Goldilocks zone of behaviors), including:

1. Domain 3/position 1-dependent 3_10_ helical m-shaped loop conformation (corresponding to 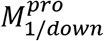.
2. Domain 3/position 2-dependent extended m-shaped loop state (corresponding to 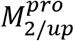.
3. 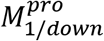-dependent substrate association to the open S1 pocket.
4. Substrate-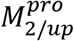-dependent dimerization, in which the monomer is stabilized by bound substrate in the dimer compatible conformation.
5. Substrate-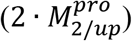-dependent catalysis, in which the oxyanion hole is aligned in the crest B up position.

We further showed that solvation free energy is ideally suited for powering the aforementioned rearrangements via counter-balanced, location-/state-specific H-bond enriched and depleted solvation, the de-solvation and re-solvation of which govern the rates of entry and exit of molecular populations to/from the available enzyme states (including substrate and inhibitor-bound states).

Lastly, we challenged the reported enzyme kinetics data for M^pro^, in which the enzyme efficiency and inhibitory requirements may be grossly underestimated by the classical Michaelis-Menten approach used in those studies.

## Acknowledgements

The authors thank Dr. Andrei Golosov for his important contributions to the WATMD parts of this work, and helpful suggestions during the manuscript preparation.

## Notes

### Competing Interest Statement

The authors have declared no competing interest.

