## Supplemental information 1 for "Probing the dynamic structure-function and structure-free energy relationships of the corona virus main protease with Biodynamics theory"

### Supplementary Information

Wan et al.

Rearrangements of the H-bond network within the domain {1-2}-3 interface (same as Figure 10)

2QCY

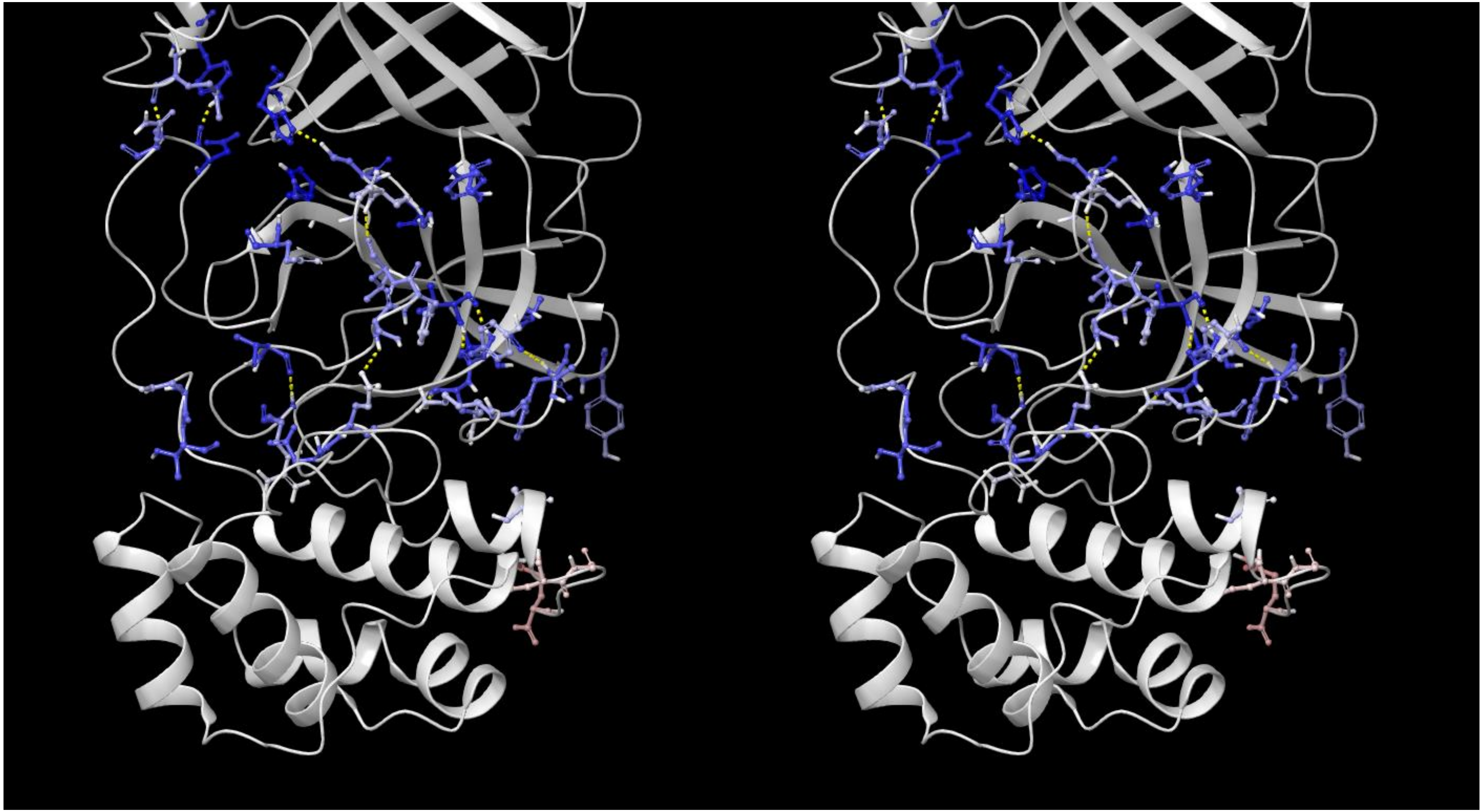

6WNP

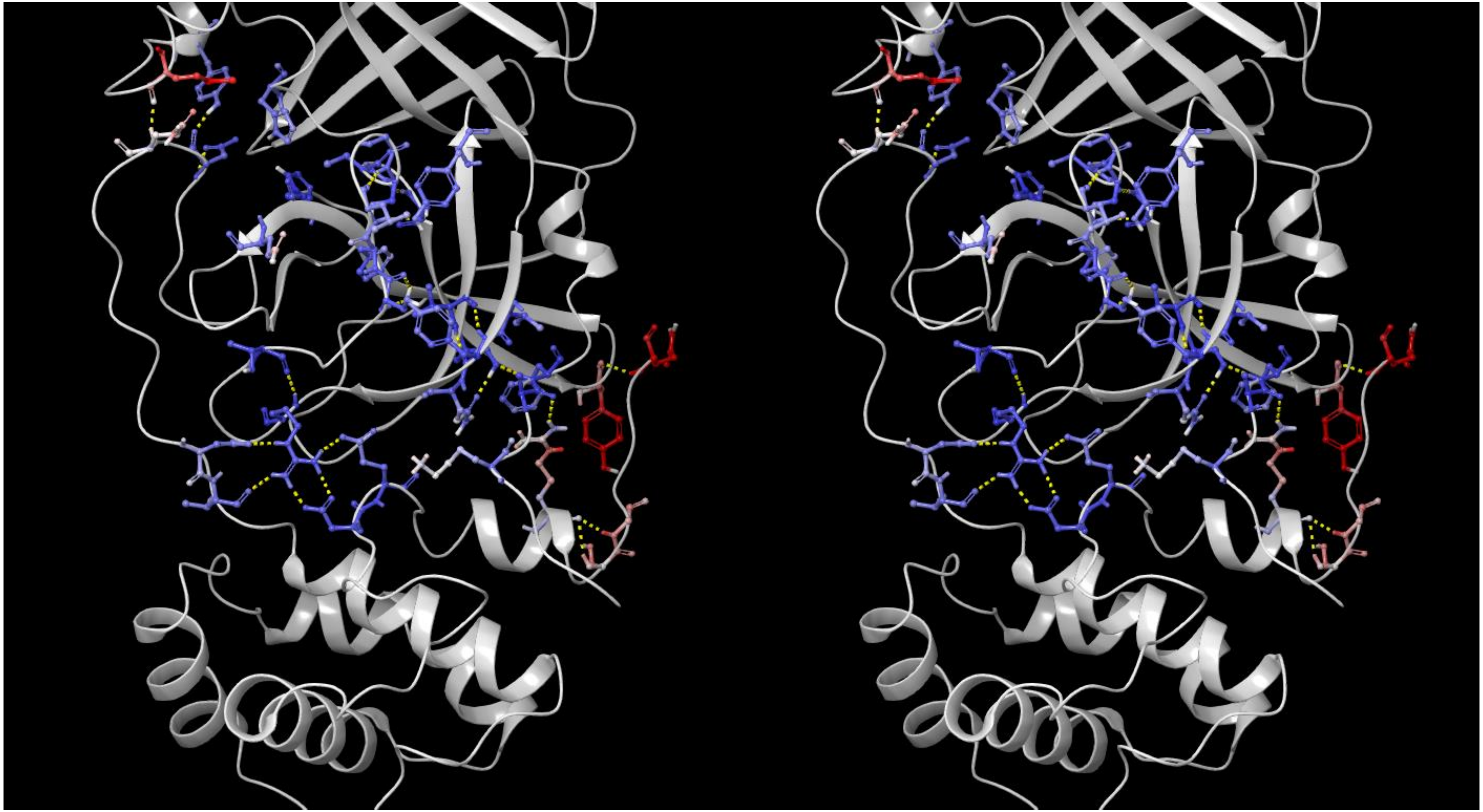

6M03

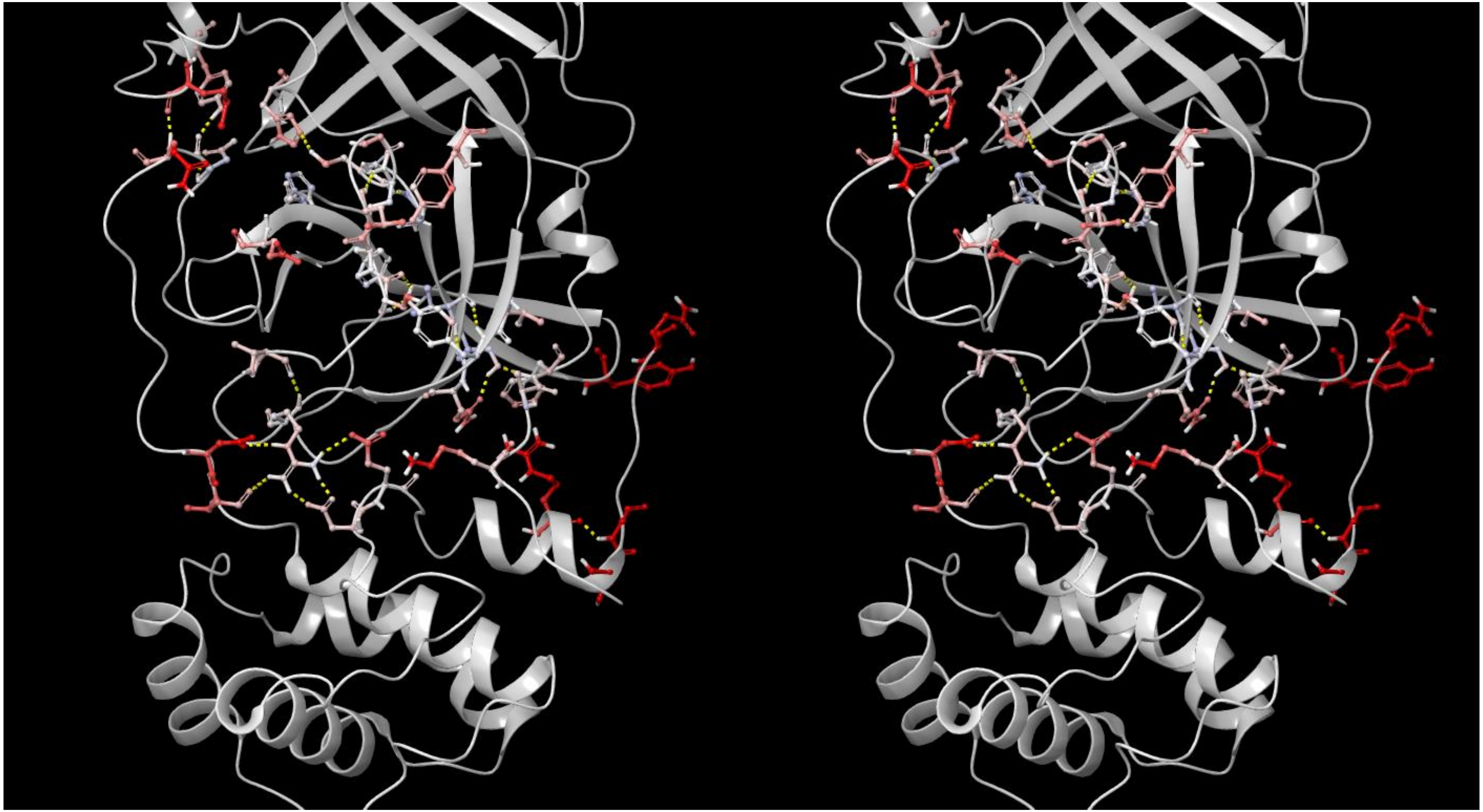

2QCY

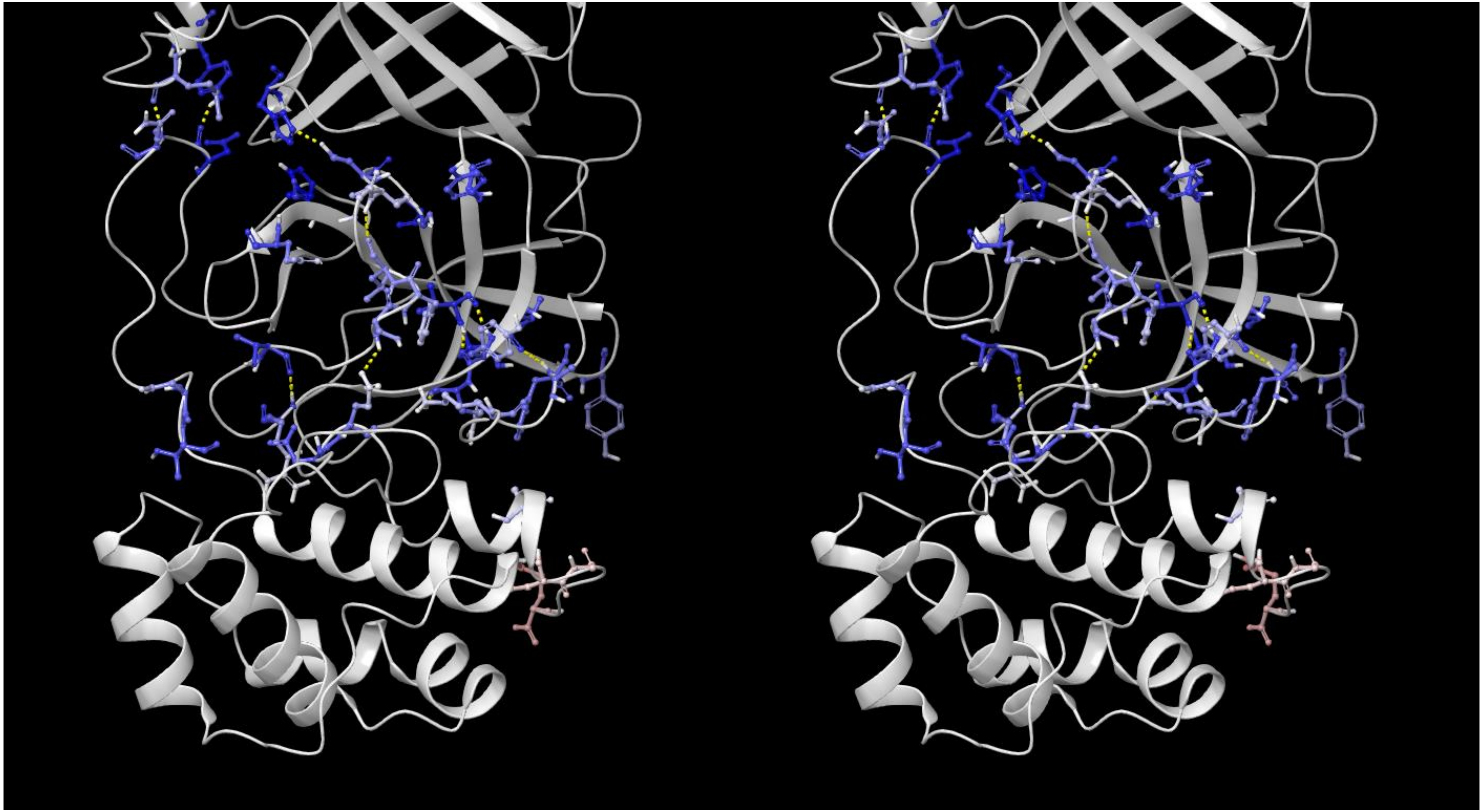

2BX3

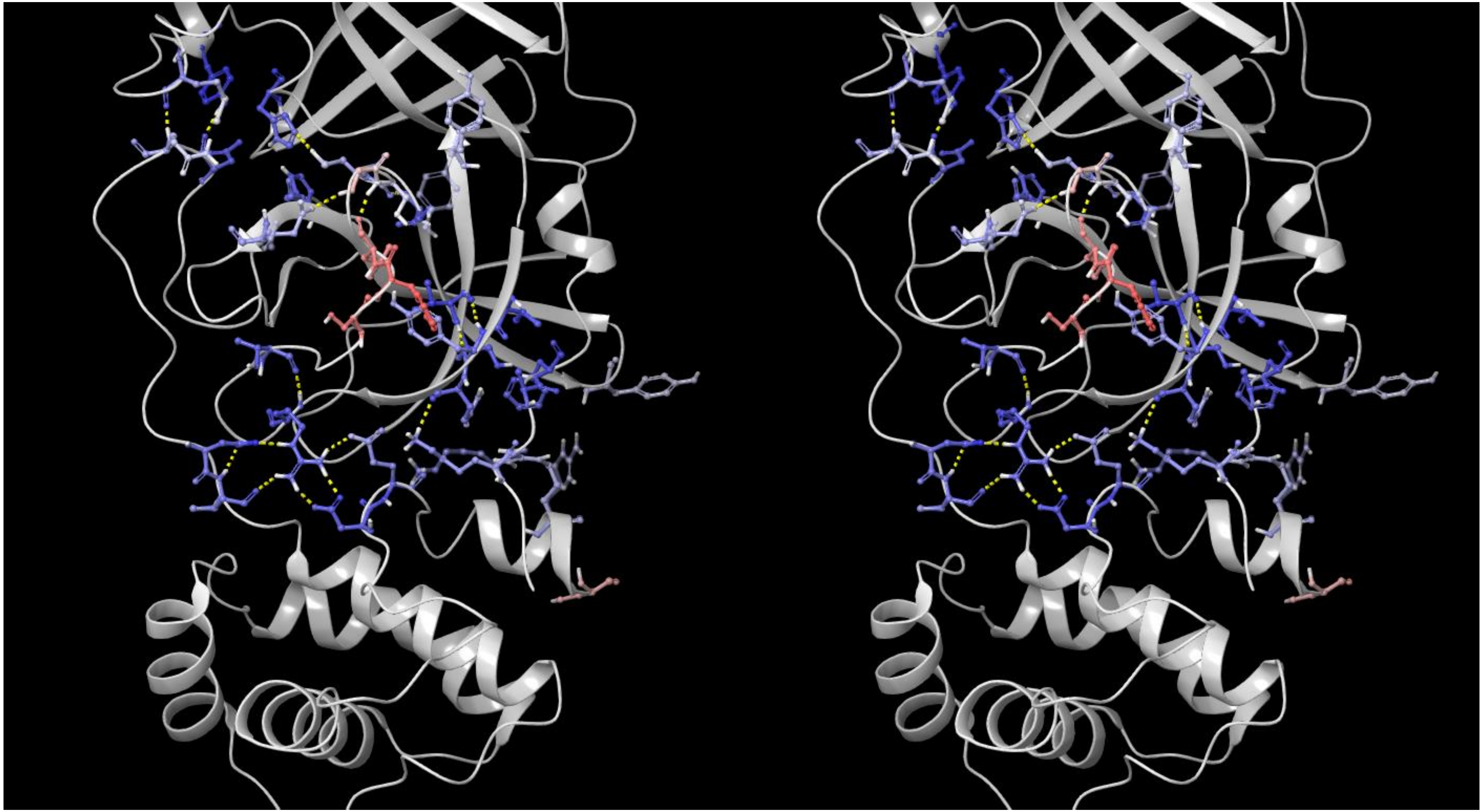

# 2G6G

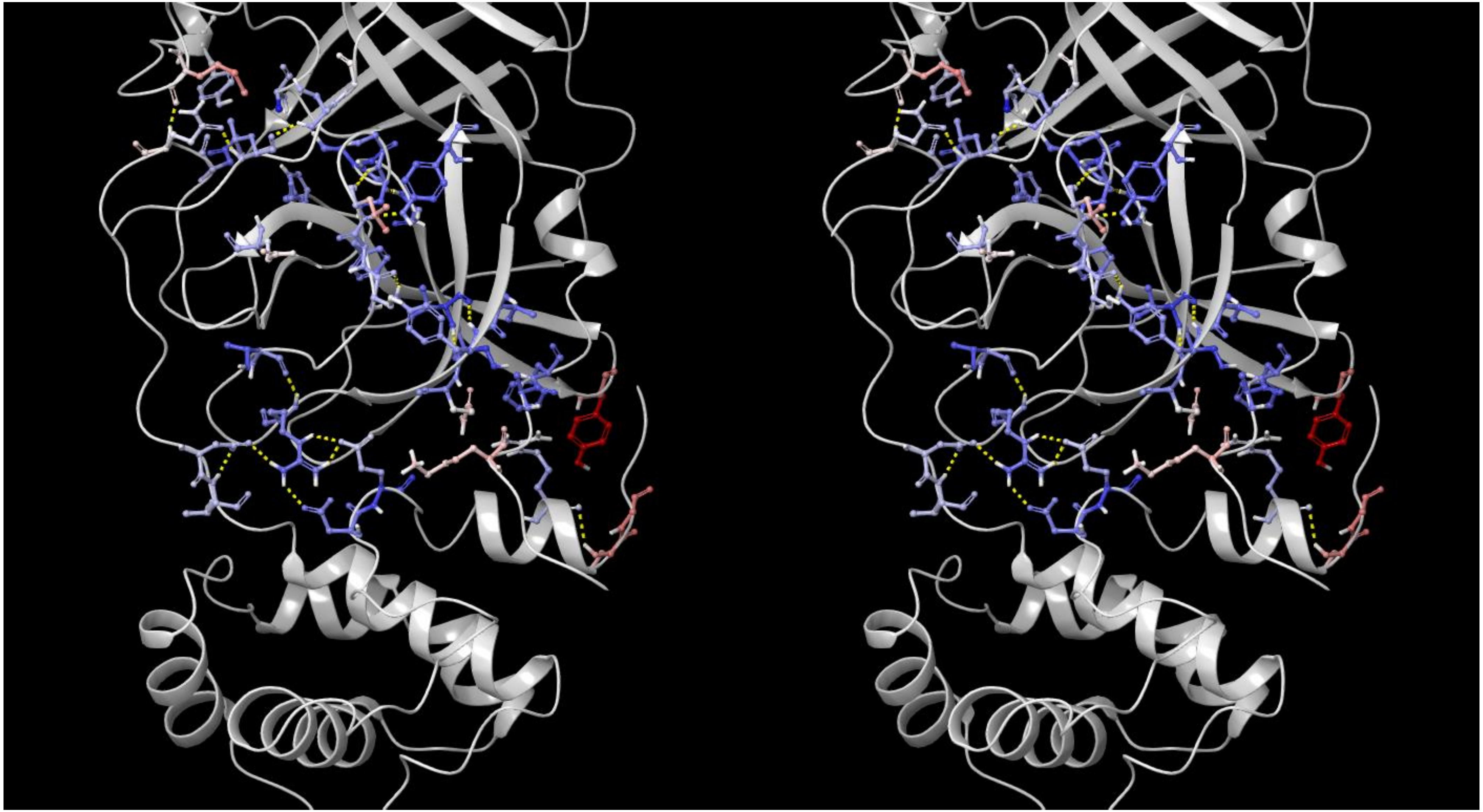

# 6LU7

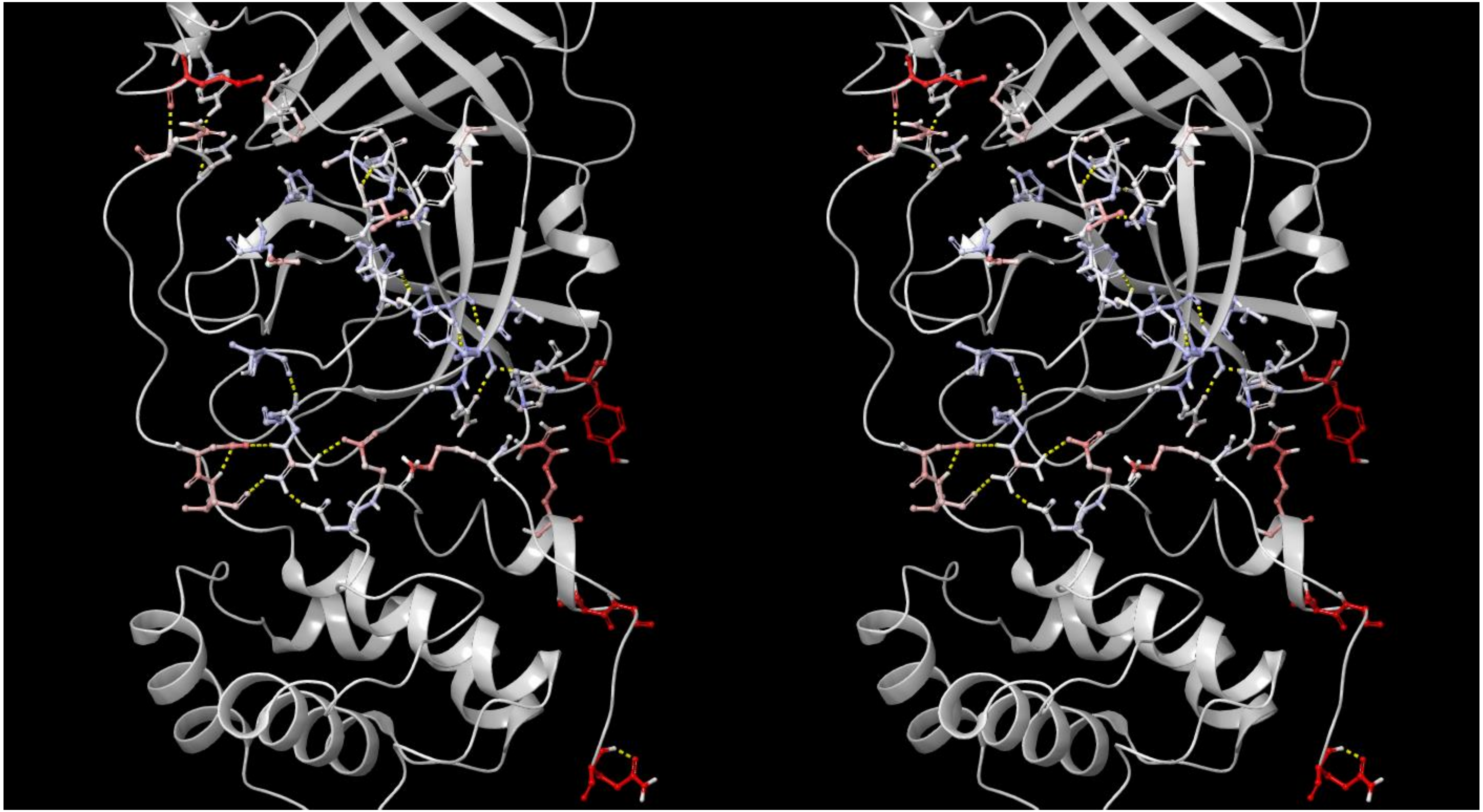

Overlays of representative crystallized ligands on the WATMD-calculated solvation hotspots of 2QCY (same as Figure 26)

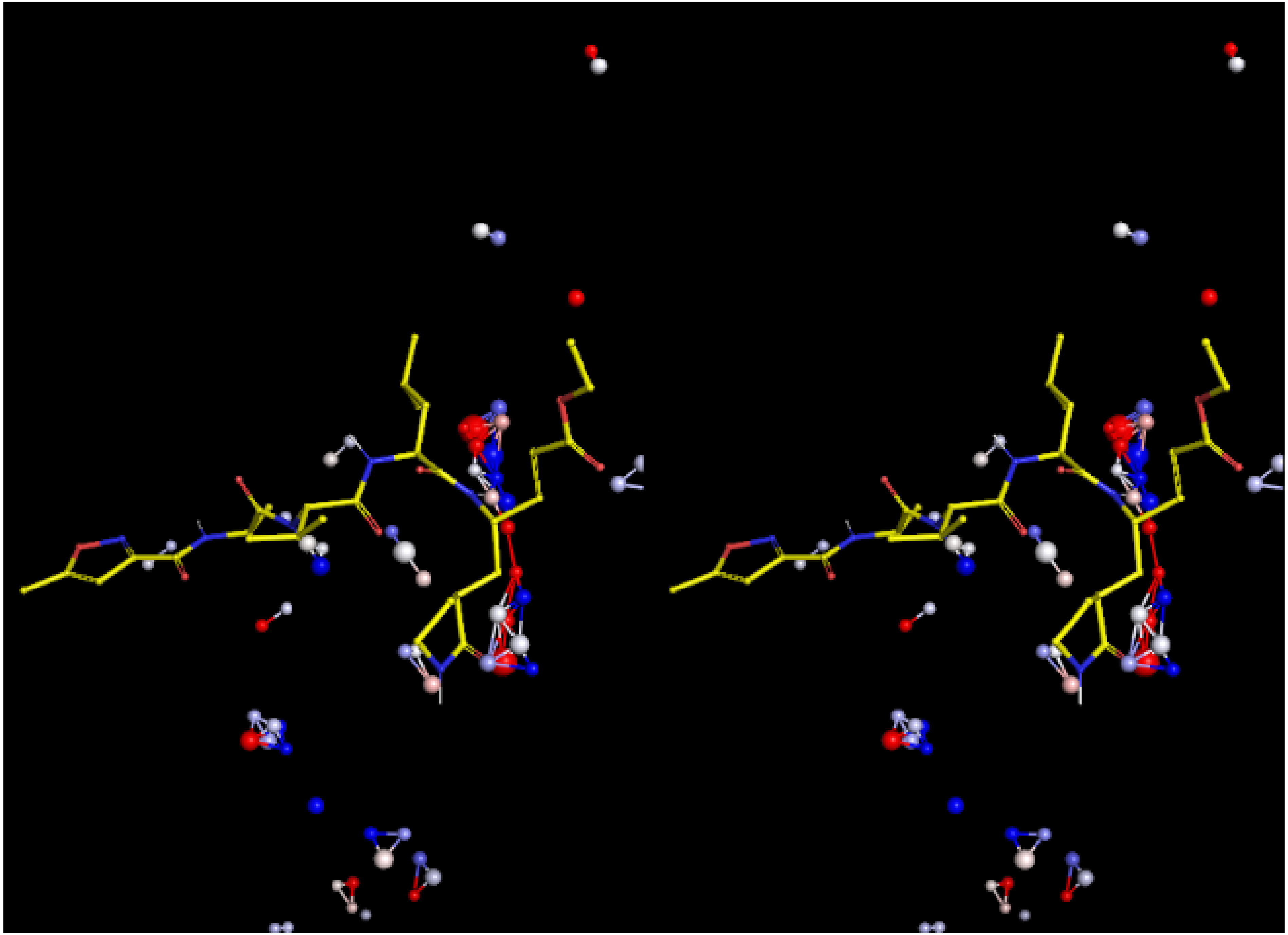

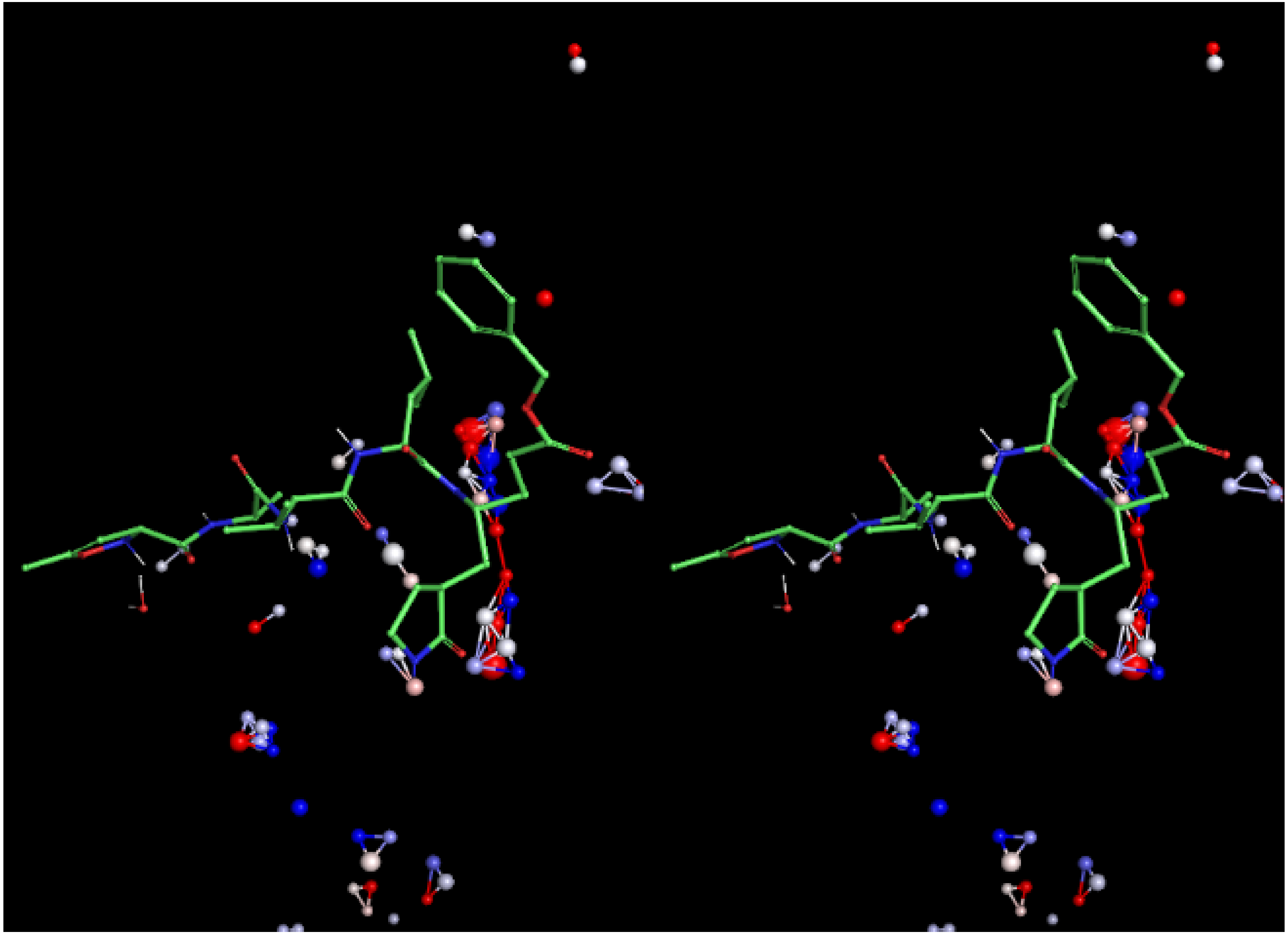

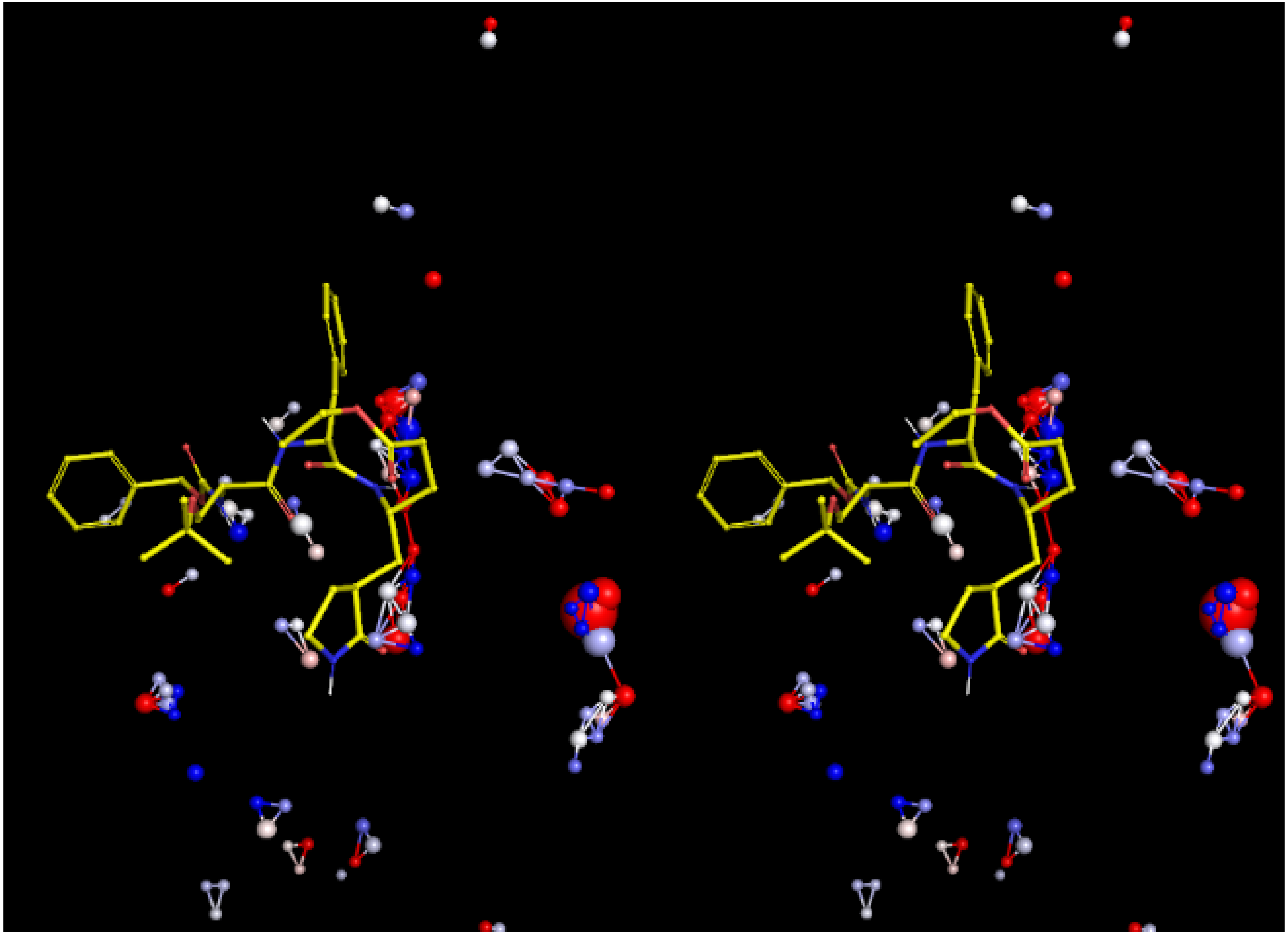

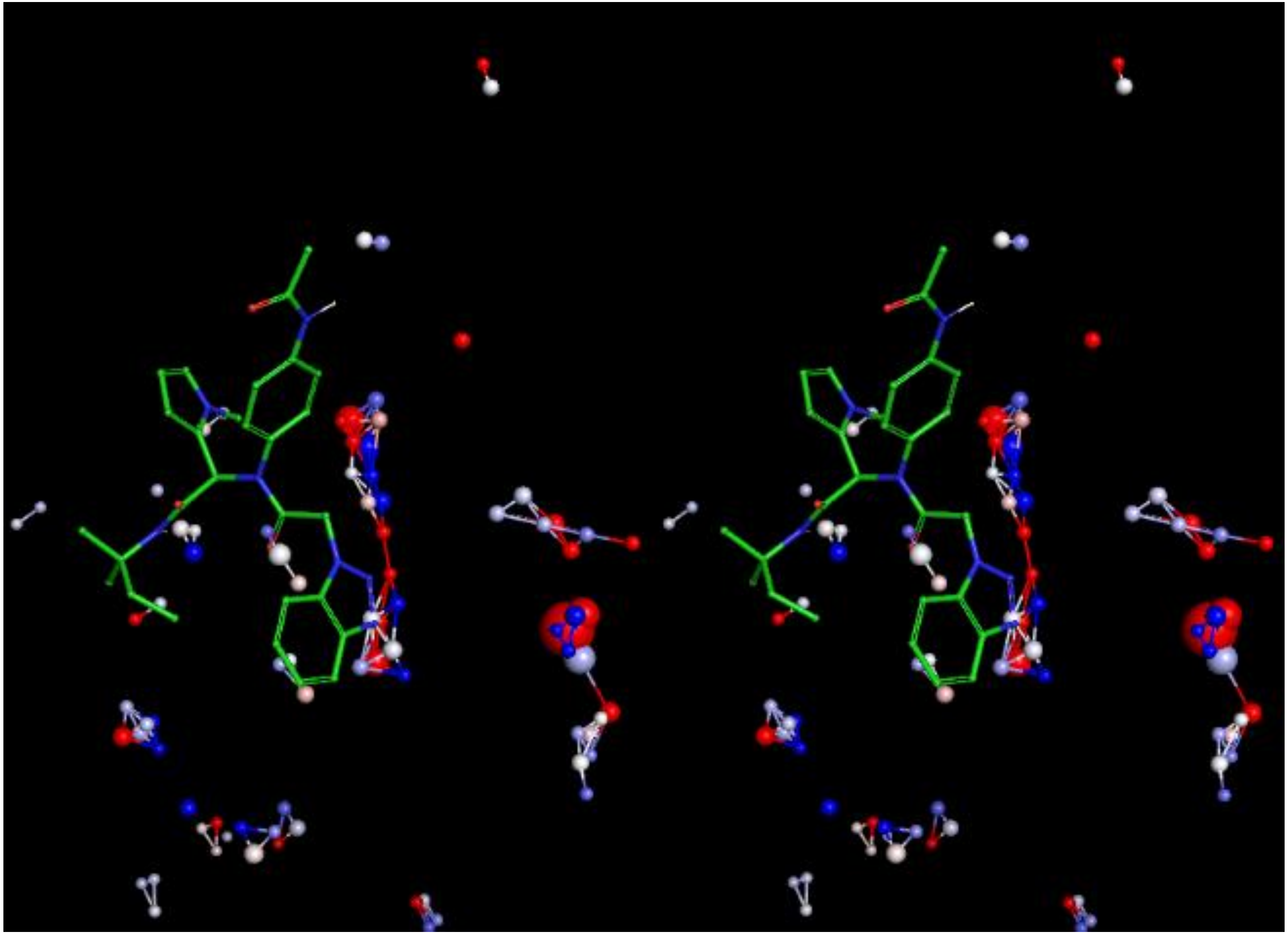

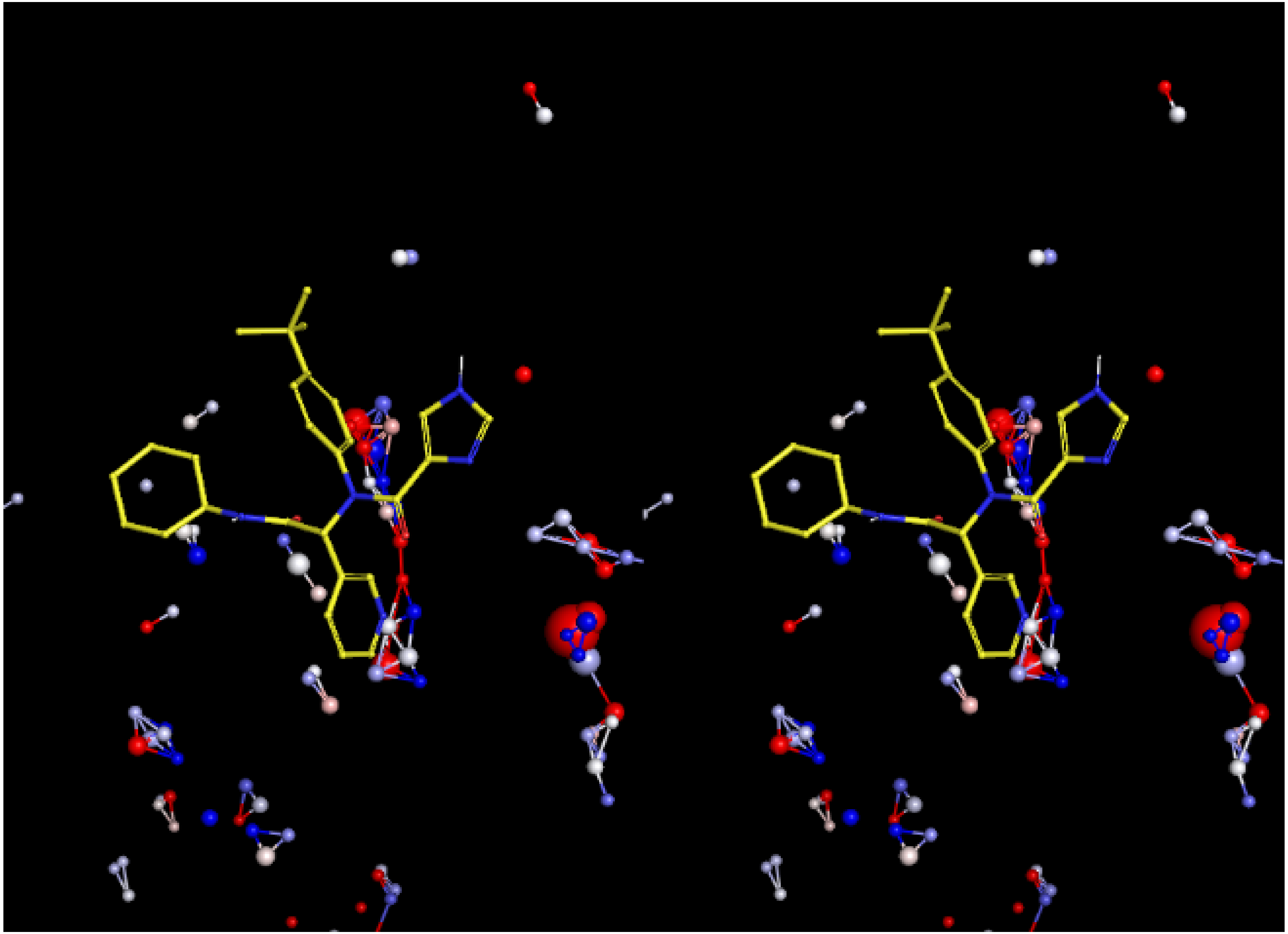

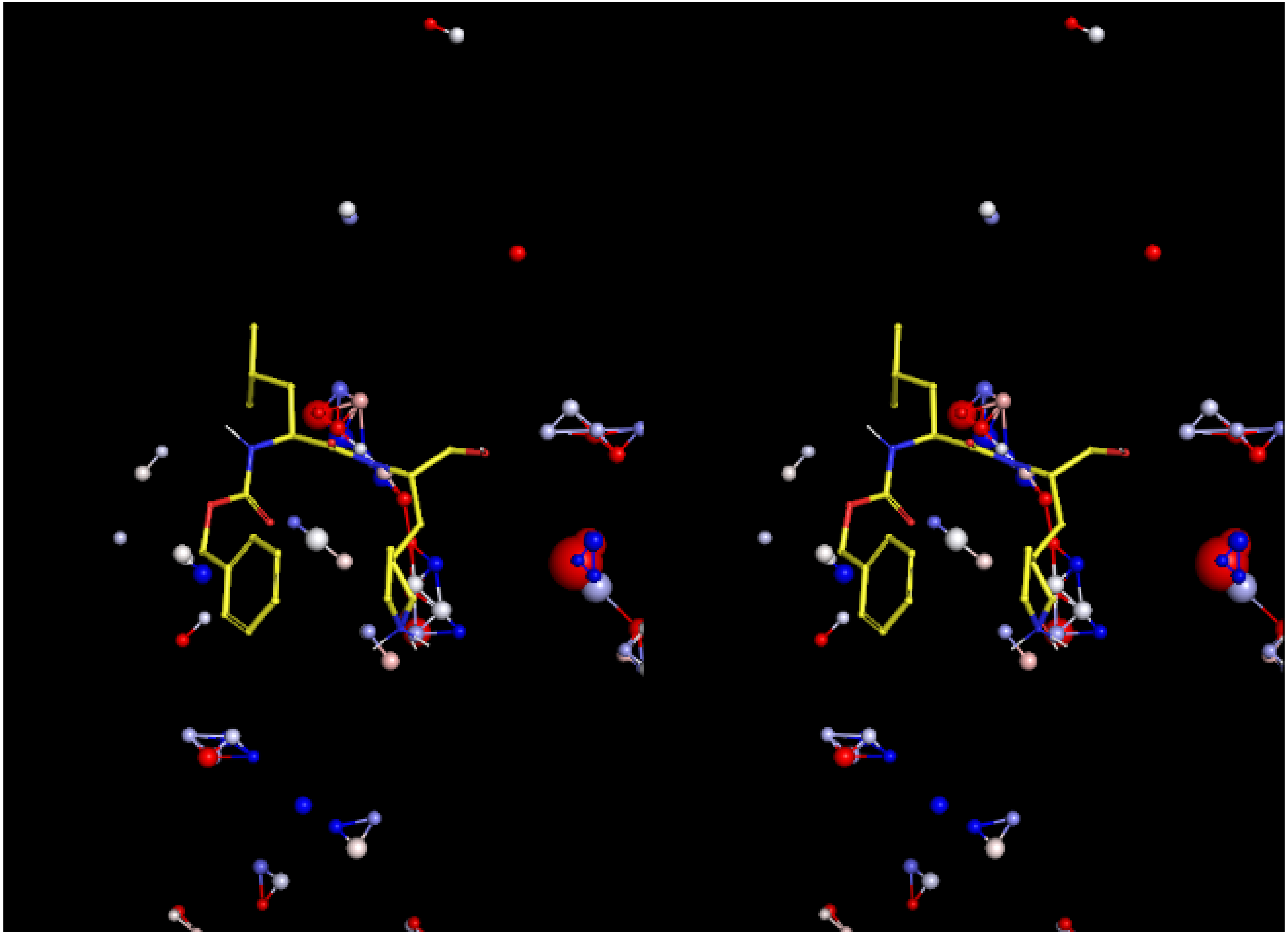
